# Population dynamics of choice representation in dorsal premotor and primary motor cortex

**DOI:** 10.1101/283960

**Authors:** Diogo Peixoto, Roozbeh Kiani, Chandramouli Chandrasekaran, Stephen I. Ryu, Krishna V. Shenoy, William T. Newsome

## Abstract

Studies in multiple species have revealed the existence of neural signals that lawfully co-vary with different aspects of the decision-making process, including choice, sensory evidence that supports the choice, and reaction time. These signals, often interpreted as the representation of a decision variable (DV), have been identified in several motor preparation circuits and provide insight about mechanisms underlying the decision-making process. However, single-trial dynamics of this process or its representation at the neural population level remain poorly understood. Here, we examine the representation of the DV in simultaneously recorded neural populations of dorsal premotor (PMd) and primary motor (M1) cortices of monkeys performing a random dots direction discrimination task with arm movements as the behavioral report. We show that single-trial DVs covary with stimulus difficulty in both areas but are stronger and appear earlier in PMd compared to M1 when the stimulus duration is fixed and predictable. When temporal uncertainty is introduced by making the stimulus duration variable, single-trial DV dynamics are accelerated across the board and the two areas become largely indistinguishable throughout the entire trial. These effects are not trivially explained by the faster emergence of motor kinematic signals in PMd and M1. All key aspects of the data were replicated by a computational model that relies on progressive recruitment of units with stable choice-related modulation of neural population activity. In contrast with several recent results in rodents, decision signals in PMd and M1 are not carried by short sequences of activity in non-overlapping groups of neurons but are instead distributed across many neurons, which once recruited, represent the decision stably during individual behavioral epochs of the trial.

## Introduction

When navigating in traffic, a driver constantly integrates evidence about the outside world that must inform upcoming decisions: stay on the throttle, press the brake, switch gears, etc. This process of deliberating on available sensory evidence to reach a commitment to a specific proposition or action is termed perceptual decision-making (Hanks and Summerfield, 2017, Brody and Hanks, 2016, Murakami and Mainen, 2015, Shadlen and Kiani, 2013, Gold and Shadlen, 2007). In the driver example, the actions involve limb movements, and in such contexts, primary motor cortex (M1) and dorsal premotor cortex (PMd) are thought to be involved in the decision-making process (Guo et al., 2014, Thura and Cisek, 2014, Cisek, 2012, Cisek and Kalaska, 2010, Cisek, 2007, Cisek and Kalaska, 2005, Wise, 1985, Vaadia et al., 1988, Wise et al., 1997). In particular, lesion (Passingham, 1985), inactivation (Kurata and Hoffman, 1994, Sasaki and Gemba, 1986) and electrophysiological studies (Cisek and Kalaska, 2005, Song and McPeek, 2010, Hoshi, 2013) suggest an important role for PMd and M1 in action selection and visuomotor association. Recent studies have employed more sophisticated perceptual discrimination tasks with arm movements as the operant response (Thura and Cisek, 2014, Chandrasekaran et al., 2017, Coallier et al., 2015) and shown that firing rates of a diverse neural population in PMd covaries with choice, stimulus difficulty, and reaction time (RT) well before the movement onset. These results are consistent with a role for PMd and M1 in “somatomotor” decisions and also suggest the presence of a candidate DV, organized by cortical laminae, in these brain areas (Thura and Cisek, 2014, Thura and Cisek, 2016, Coallier et al., 2015, Palmer et al., 2005, Wang et al., 2016, Chandrasekaran et al., 2017).

With few exceptions (Bollimunta et al., 2012, Kaufman et al., 2015, Kiani et al., 2014b, Ponce-Alvarez et al., 2012, Rich and Wallis, 2016), neurophysiological studies of decision mechanisms have focused on *average* decision-related signals at the single neuron level (Roitman and Shadlen, 2002, Churchland et al., 2008, Kiani and Shadlen, 2009, de Lafuente et al., 2015) and at the population level (Mante et al., 2013, Machens et al., 2010, Raposo et al., 2014). Key questions remain unanswered about the single-trial dynamics and spatiotemporal structure of neural population responses in perceptual decision formation in PMd and M1. We therefore trained macaque monkeys to perform fixed as well as variable-duration random-dot motion direction discrimination tasks (Kiani et al., 2008) using an arm movement as the operant response while simultaneously recording hundreds of neurons using Utah arrays implanted in PMd and M1. We used decoding techniques to estimate single-trial DVs from PMd and M1 firing rates, and examined how the dynamics of these decoded DVs changed with parameters such as stimulus difficulty and uncertainty about expected stimulus duration. Our analyses focused on three interconnected questions.

First, we analyzed the relationship between single-trial dynamics of the DV and sensory stimuli that inform the choice, and determined whether these dynamics differ between PMd and M1. We then tested whether the neural dynamics change when subjects transition from a context of temporal certainty to high temporal uncertainty about stimulus duration (Shadlen and Newsome, 2001, Murphy et al., 2016).

Second, we used computational modeling of behavior and neural responses to identify mechanisms that can explain the observed dynamics of choice representation in PMd and M1 under different task conditions. Bounded accumulation of evidence is a widely used modeling framework for perceptual decisions in the direction discrimination and similar sensory tasks (Ratcliff and McKoon, 2007). For a binary choice, the model assumes that two accumulators integrate sensory evidence in favor of the two competing options until one of the accumulators reaches a decision threshold or bound (Vickers, 1970, Link, 1992, Beck et al., 2008, Shadlen and Kiani, 2013, Kiani et al., 2014a). We considered three variants of this model that could offer a theoretical account for accelerated representation of choice under temporal uncertainty: increased urgency or reduced bound (Churchland et al., 2008, Heitz and Schall, 2012, Purcell and Kiani, 2016, Hanks et al., 2014), increased input gain (Cisek et al., 2009, Thura et al., 2012), and a novel framework based on progressive recruitment of choice representing neurons which suggests that the fraction of neurons carrying choice related signals increases throughout the trial, leading to increasingly more stable choice representation over the course of the trial.

Our final goal was to understand how a stable choice representation is implemented by a population of PMd and M1 neurons. We considered two competing hypotheses: sustained representation of choice by: 1) a stable population of neurons, or 2) a sequence of transient responses in non-overlapping groups of neurons. Sustained responses are commonly observed in the frontoparietal cortex of the primate brain and are implicated as a substrate for working memory, decision-making, and higher cognitive functions (Chandrasekaran et al., 2017, Churchland et al., 2008, Cisek and Kalaska, 2005, Kiani and Shadlen, 2009, Machens et al., 2010, Mante et al., 2013, Roitman and Shadlen, 2002, Shadlen and Newsome, 2001, Thura and Cisek, 2014, Goldman-Rakic, 1995). An alternative mechanism has recently emerged from optical imaging studies in rodents (Harvey et al., 2012, Morcos and Harvey, 2016) and later expanded to electrophysiological and computational studies (Rajan et al., 2016, Scott et al., 2017), suggesting that decision-related activity may be carried by transient sequences of small subsets of neurons at different points in time. The “sequence” model predicts that in a given epoch in the trial, only a small subset of neurons represents the decision, and this subset will be largely non-overlapping with the subset of decision related neurons for any other epoch in the trial.

We found that, in the fixed duration task, single-trial DVs are represented in both PMd and M1 shortly after stimulus onset but are stronger and faster in PMd compared to M1. On single trials, estimated DVs exhibit ramp-like growth during stimulus presentation and the slope is steeper for easier coherences compared to harder coherences. In the variable duration task, for both PMd and M1, DV dynamics are strikingly accelerated and are nearly identical in the two areas throughout the entire trial, although PMd responses still lead M1 responses by ∼15 ms. Even though single trial DVs ramp up faster in the variable-duration task, the co-variation of ramp slope with stimulus difficulty was preserved and behavioral accuracy remained largely stable. Control analyses show that these results are not explained by accelerated representation of motor variables such as reach speed, other kinematics of the arm and eye, or eventual RT, nor could the combined physiological and behavioral data be modeled accurately by changes in gain or urgency. Instead, the data are consistent with a progressive recruitment of choice-related neurons that is accelerated under uncertainty conditions. Consistent with the progressive recruitment model, the empirically observed population choice signals become increasingly stable throughout the stimulus presentation as more units with sustained choice selectivity are recruited. This stabilization happens more rapidly in the variable duration task, in which there is a high premium for quick decisions (Murphy et al., 2016).

## Results

### Monkeys discriminate stimulus motion better for higher coherence and longer duration trials

We trained monkeys in a variant of the classical random dot motion discrimination task (RDM, (Britten et al., 1992)), in which animals report the net direction of motion in a random dot kinematogram (Shadlen and Newsome, 2001, Kiani et al., 2008, Britten et al., 1992) presented on a LCD touchscreen (Fig. 1a). In our variant of the RDM task, the monkeys used an arm reach to one of two targets corresponding to the perceived direction of motion to report their decision (Fig. 1a-b). In the fixed duration version of the task the stimulus was always presented for 1000 ms followed by a random delay period (400-900 ms) after which the monkey was provided with a “go cue” to report its decision. Eye fixation was enforced from the beginning of the trial until appearance of the go cue to impose additional behavioral control and avoid interpretational confounds since PMd activity can be modulated by eye-hand relative position (Pesaran et al., 2006).

**Figure 1.**
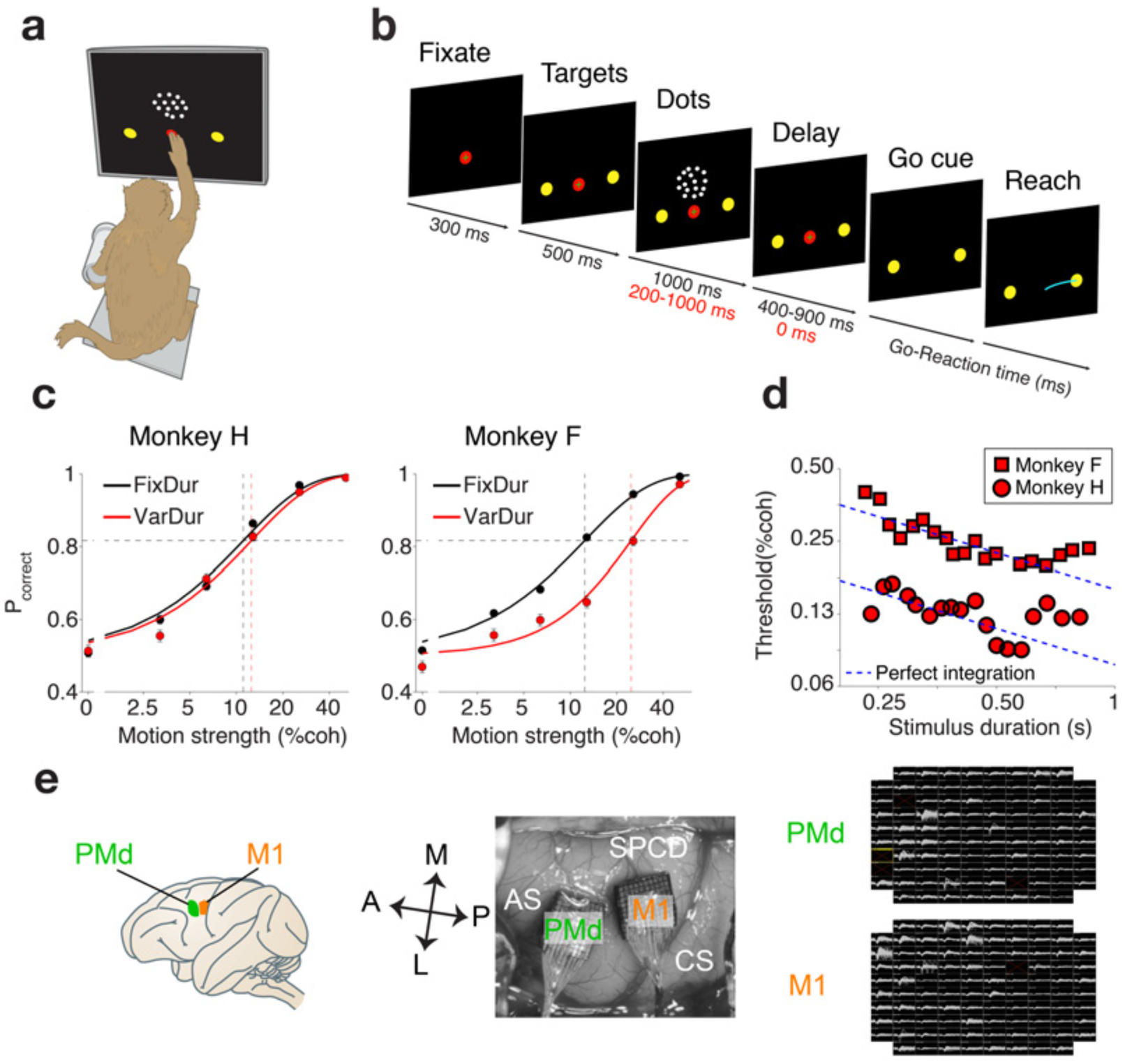
Motion discrimination task, psychophysical performance and recording locations and techniques. **a) Behavioral setup -** The monkey performed the motion discrimination task on a touchscreen using one arm, while the other arm remained gently restrained. Eye position was continuously tracked using an infrared mirror placed in front of the monkey’s eyes. **b) Direction discrimination task structure** - Trials start with the onset of a fixation point on the touchscreen. Once both eye and hand fixation are acquired two targets appear on the screen. The motion stimulus was shown after a short delay (500 ms) and lasted 1000 ms (200-1000 ms) for the fixed (variable) duration version. The dots offset was followed by a 400-900 ms delay in the fixed duration version whereas no delay was present for variable duration version. At the end of the delay, the offset of the fixation point cued the monkey to report his decision by making a hand reach movement to the appropriate target. **c) Psychophysical performance in the motion discrimination task.** Percentage correct is plotted as a function of motion coherence for the fixed duration version (black) and the variable duration task (red) for monkey H (left panel) and monkey F (right panel). Observed data points (+/- SEM) are represented by the red and black markers. The data for each task was independently fit with Weibull curves (red and black curves). 17167/ 17440 trials for the fixed duration task and 4923/5381, trials for the variable duration task for monkey H/F, respectively. **d) Psychophysical thresholds in the variable duration motion discrimination task.** Psychophysical threshold is plotted as a function of stimulus duration for monkey H (circles) and monkey F (squares). Dashed blue lines show predicted perfect accumulation for each subject. The observed performance deviates from perfect accumulation for stimuli longer then 533/682 ms for monkey H/F respectively. **e) Location of the two multielectrode arrays.** Two 96 channel Utah arrays were implanted in primary motor and dorsal premotor cortex as judged by anatomical references (left and middle panels). Example waveforms collected from PMd and M1 for the same experimental session (monkey H, right panel). White squares denote ground pins in the four corners of the arrays.

Monkeys displayed excellent behavioral performance in this task, achieving close to ceiling levels of accuracy (99% for both monkeys) for the highest coherence stimuli (Fig. 1c, black curves). The accuracy decreased smoothly with stimulus difficulty (lower coherence) and remained above chance for the lowest (non-zero) coherence stimulus (3.2% coherence, 59%-62% accuracy for monkey H-F). Psychophysical thresholds (α) (estimated at 81.6% accuracy by fitting a cumulative Weibull function to the performance curves) were 12.1% and 12.4% stimulus coherence for monkey H and F, respectively (Fig.1c, black dashed vertical lines).

After data collection was concluded in the fixed duration task, monkeys performed a variable duration RDM task, (Fig. 1c, red curves). In these experiments, stimulus duration was randomly selected on each trial (200-1000 ms exponentially distributed, median 435 ms) and the delay period was eliminated, requiring subjects to report their decision immediately after stimulus offset. Psychophysical thresholds for both monkeys decreased as stimulus duration increased up to ∼500 msec (Fig. 1d) indicating that monkeys performed better for longer stimuli. This improvement in performance suggests that additional visual evidence was utilized to improve decisions as a result of integrating the sensory evidence for a longer duration. The improvement in thresholds occurred at the rate expected from a perfect integrator model (slope, ∼ -0.5) for stimulus durations up to approximately 533 and 682 ms for monkey H and F, respectively, with little or no improvement for longer stimuli (Kiani et al., 2008, Kiani and Shadlen, 2009).

The two tasks enabled us to probe the dynamics of decision-related signals in PMd/M1. The fixed duration task provided temporal separation between evidence integration (dots period), action planning (hold period), and action execution (post-go period). In contrast, the variable duration task provided the ability to query the subject’s choice as soon as the stimulus is terminated.

### Single trial choice signals in PMd and M1 are compatible with the neural representation of a decision variable

We recorded neural activity in PMd and M1, using two chronically implanted 96-channel Utah arrays (Fig. 1e), while subjects performed each motion discrimination task. Consistent with prior studies in PMd (Cisek and Kalaska, 2005, Chandrasekaran et al., 2017), we found diverse responses at the single cell level, which may reflect multiple functions being implemented in this area (Supp. Fig. 1). The same observation was true for M1 (Supp. Fig. 2).

Our primary goal was to understand the dynamics of these diverse neural responses in PMd and M1 at the population level—both on average and on single trials. We trained a regularized logistic classifier to predict right and left choices on individual trials based on short periods of neural activity (50 ms windows), using a method we developed in a recent study (Kiani et al., 2014b). Classification was based on features of the simultaneously recorded activity from ∼100-200 units from each area (Kiani et al., 2014b) (median number of units/session: 153 for PMd, 147 for M1).

For the fixed duration task, choice prediction accuracy before and immediately after the onset of the targets was near chance (Fig. 2a, Supp. Fig. 3a and 4a) suggesting that choice biases prior to motion onset were negligible and the monkey’s choices were shaped by the motion stimulus. Choice prediction accuracy rose above chance at 187 ± 12 ms (mean±s.d.) after stimulus onset for PMd and 240 ± 14 ms (mean±s.d.) for M1. The median latencies for PMd were significantly shorter than for M1 (Wilcoxon rank sum test, p<0.001) and were similar to other reports of decision-related signals in PMd (Thura and Cisek, 2014, Chandrasekaran et al., 2017) and MIP (de Lafuente et al., 2015). Although our monkeys were extensively trained on this version of the task that allowed them 1000 msec to evaluate the stimulus motion and at least another 400 msec of delay period to prepare the operant response, both PMd and M1 still responded in a choice predictive manner less than 250 ms after the stimulus onset and over a second before the initiation of the action was cued. Thus, choice predictive activity in these (pre)motor structures, similar to their parietal counterparts (Shadlen and Newsome, 2001) is not contingent on having to prepare for immediate action, but is also present when that action is delayed ∼1-2 seconds into the future.

**Figure 2.**
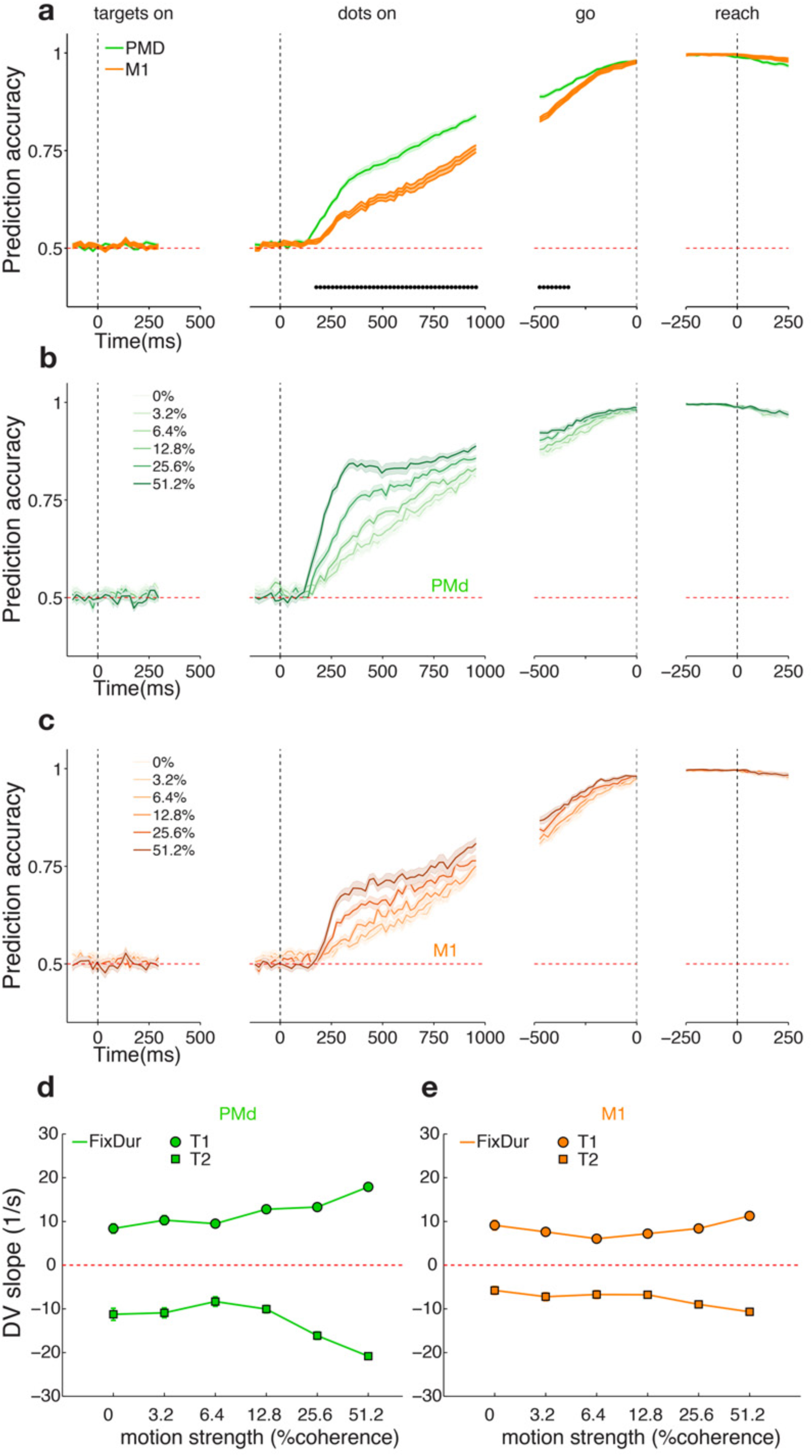
Neural population choice prediction accuracy on single trials in the fixed duration task (pooled results across 2 monkeys). **a) PMd is more choice predictive than M1 during the stimulus presentation but not later in the trial.** Average prediction accuracy (see Methods) over time +- SEM for both monkeys. PMd (M1) data are plotted in green (orange). Black dots denote time bins for which the prediction accuracy was significantly different between the two areas (Wilcoxon signed rank test, p<0.05 Holm-Bonferroni correction for multiple comparisons). **b) Choice prediction accuracy in PMd rises faster for easier trials during stimulus presentation.** Average choice prediction accuracy as function of stimulus difficulty. Easy stimuli are represented in darker tones while harder stimuli are plotted in lighter tones, and shading corresponds to +- SEM. Same data as **a)** (green trace), except prediction accuracy is calculated individually for each stimulus difficulty. **c) Choice prediction accuracy in M1 rises faster for easier trials during stimulus presentation.** Same data as **a)** (orange trace), except prediction accuracy is calculated individually for each stimulus difficulty. Figure conventions as in **b).** **d) Single-trial decision variable slopes in PMd co-vary with stimulus coherence.** Average of single trial DV slopes are plotted as a function of stimulus coherence and choice. Positive values (circles) correspond to T1 (right) choices and negative values (squares) to T2 (left) choices (correct trials only). Error bars indicate +/- SEM across trials. **e) Single-trial decision variable slopes in M1 co-vary with stimulus coherence. Same as d) for M1.**

Choice prediction accuracy rose steadily for both areas as the trial proceeded, but was significantly higher for PMd than for M1 (P<0.05 Wilcoxon Sign rank test, Holm-Bonferroni corrected) during most of the motion-viewing epoch (Fig. 2a, Supp. Fig. 3a and 4a). This difference was observed in both monkeys and did not result from a higher number of recorded units in PMd (Wilcoxon rank sum test comparing median number of units, p=0.41). At the end of the visual stimulus period prediction accuracy reached 84.5% ± 1.3% and 83% ± 0.5% (mean±s.e.m.) for PMd and 72% ± 0.9% and 78% ± 0.9% (mean±s.e.m.) for M1 of monkeys H and F, respectively (Fig. 2a; Supp. Fig. 3a and 4a). These classification accuracies roughly matched (in M1) or exceeded (in PMd) those previously reported for neural population recordings in prearcuate cortex using similar recording and analysis techniques (Kiani et al., 2014b) (and could be even further increased by adjusting the window size Supp. Fig. 5), confirming the possibility of obtaining single trial read-outs of a decision state from these areas.

Highly reliable choice predictive activity with short latencies is expected from standard accumulation-to-bound models of decision formation (Mazurek, 2003, Cisek et al., 2009). The second expectation is that the rate of increase of choice predictive activity should depend on stimulus difficulty (Gold and Shadlen, 2007). Consistent with this expectation, classification accuracy on easier trials rose faster and attained higher values compared to harder trials (Fig. 2b-c, Supp. Fig. 3b-c and 4b-c). This feature was present in both areas though the separation between stimulus difficulties was stronger in PMd than M1 between 200-600 ms aligned to stimulus onset (Wilcoxon sign rank test, P<0.005, Supp. Fig. 6). The third expectation is that the relationship between classification accuracy and motion coherence be stronger during the first half of the dots period and becomes smaller as the trial unfolds (Shadlen and Newsome, 2001, Roitman and Shadlen, 2002, Kiani et al., 2008, Wang, 2002) (Supp. Fig. 6, 200-600 vs 600-1000 ms periods). Finally, during the delay and peri-movement periods prediction accuracy is high with little or no separation by stimulus difficulty, suggesting that a categorical decision is made by the end of the delay period in the vast majority of trials. Overall, the dynamics of the choice prediction accuracy matched accumulation of evidence and commitment to a choice when adequate evidence was accumulated.

To better understand the dynamics of decision-related activity, we calculated a continuous readout of the strength of the model’s prediction for the subject’s choice, which is critical for single trial analyses to follow. We calculated the logistic model’s log odds ratio for the two choices for each time point on every trial. This variable is equal to the distance of the neural population activity from the classifying hyperplane (Supp. Fig. 7a). As in our previous study (Kiani et al., 2014b), we interpreted this distance as the model’s decision variable (DV) and used it as a proxy for the internal cognitive state of the animal, representing a preference for a given choice. Because the DV is continuous (unlike predicted choice which is binary) and can differ even between correctly predicted trials (Supp. Fig. 7a), it provided a continuous metric for quantifying the internal cognitive state and its dependency on stimulus difficulty. Our convention was that positive values of the DV reflect increased likelihood of right choices and negative values reflect left choices. As expected the average decision variable showed the same effects found for choice prediction accuracy: (i) its magnitude increases with time and with stimulus coherence (Supp. Fig. 7b), (ii) the coherence-dependent separation of the DVs depends on the time in the motion viewing period, and (iii) this separation vanishes around the time of Go cue and motor response.

We next investigated whether these effects of stimulus difficulty held on *single trials.* If the DV traces truly ramped up on single trials, their slopes should increase as a function of coherence. Alternatively, if the entire system synchronously stepped from an uncertain state to a committed state at different points in time for different trials (with ramping being an artifact of averaging multiple trials) the slopes should not depend on stimulus coherence. Our results cannot exclude the possibility that population-level ramping is implemented by asynchronously stepping neurons (Latimer et al., 2015) whose step times are coherence dependent.

To quantify stimulus coherence effects on the single-trial DV we focused on the first half of the stimulus presentation interval (500 msec). This time window was consistent with a conservative estimate of the motion integration times from psychophysical data for both monkeys (Fig. 1d). We used a tri-linear fit to single-trial DV traces. The fitted function consisted of an initial interval of zero slope, reflecting the finite latency between dots onset and initial modulation of PMd/M1 activity. The slope during the second interval captures a period of rapid DV change following dots onset, while the third interval reflects a general slowing of DV change that occurs by the middle of the dots period (Supp. Fig. 7b-d, see Methods). The tri-linear fit enables us to focus on the rate of rise of the DV during decision-making (Shadlen et al., 2016). Consistent with the ramping representation, at the population level, of a decision variable on single trials in these areas, higher coherence trials are associated with steeper DV slopes (second slope of the tri-linear fits, Figure 2d-e, Supp. Fig. 3d-e and Supp. Fig. 4d-e). The results are significant for both areas and choice directions (for both models: slope vs coherence and slope vs log_2_(coherence); see Methods, statistical analyses Supp. Table 1).

### Stimulus duration uncertainty increases and accelerates choice predictive activity in both areas

So far, we focused on the neural activity from the fixed duration task for which the animals consistently had 1 second of visual evidence to deliberate and decide upon. However, in the real world, subjects rarely know the precise timing of visual information relevant to making a choice. Thus, after experiments on the fixed duration task, we introduced the variable duration task and trained the monkeys to report their decision immediately upon termination of the stimulus (Fig. 1a,b; Methods). Prior to these recordings the monkeys had never been exposed to short duration stimuli (< 1000 ms).

Since subjects could not predict the duration of the stimulus on single trials and most trials were short (median 435 ms, see Methods), the variable-duration task incentivized accurate assessment of sensory evidence early in the stimulus presentation period: the first 200 ms of dots motion were guaranteed to be shown but stimulus presentation could be terminated at any point thereafter. We asked whether the dynamics of decision-related signals in PMd and M1 were different in the variable duration task. In both areas, we found that classification accuracy increased faster in the variable duration task leading to much higher accuracy values during the stimulus presentation period (Fig. 3a, Supp. Fig. 8a, Supp. Fig. 9a). This acceleration in prediction accuracy was most apparent in M1, where choice predictive neural responses emerged much faster in the variable duration task (193 ± 12 ms compared to 240 ± 14 ms in the fixed duration task). This earlier onset also happened in PMd (177 ± 8 ms compared to 187 ± 12 ms in the fixed duration task), though to a lesser extent. Consequently, the difference in the onset time of choice-related activity between PMd and M1 diminished substantially in the variable duration task (latency difference= 13.2 ms in variable duration versus 41.6 ms in fixed duration task, p=1.6 x 10^−13^, Wilcoxon rank sum test). Moreover, the difference in absolute prediction accuracy between the two areas was significant for only 80 ms during the entire dots period (P<0.05 Wilcoxon Rank Sum test, Holm-Bonferroni corrected), confirming that these areas represent the upcoming choice with very similar strength in the variable duration task.

**Figure 3.**
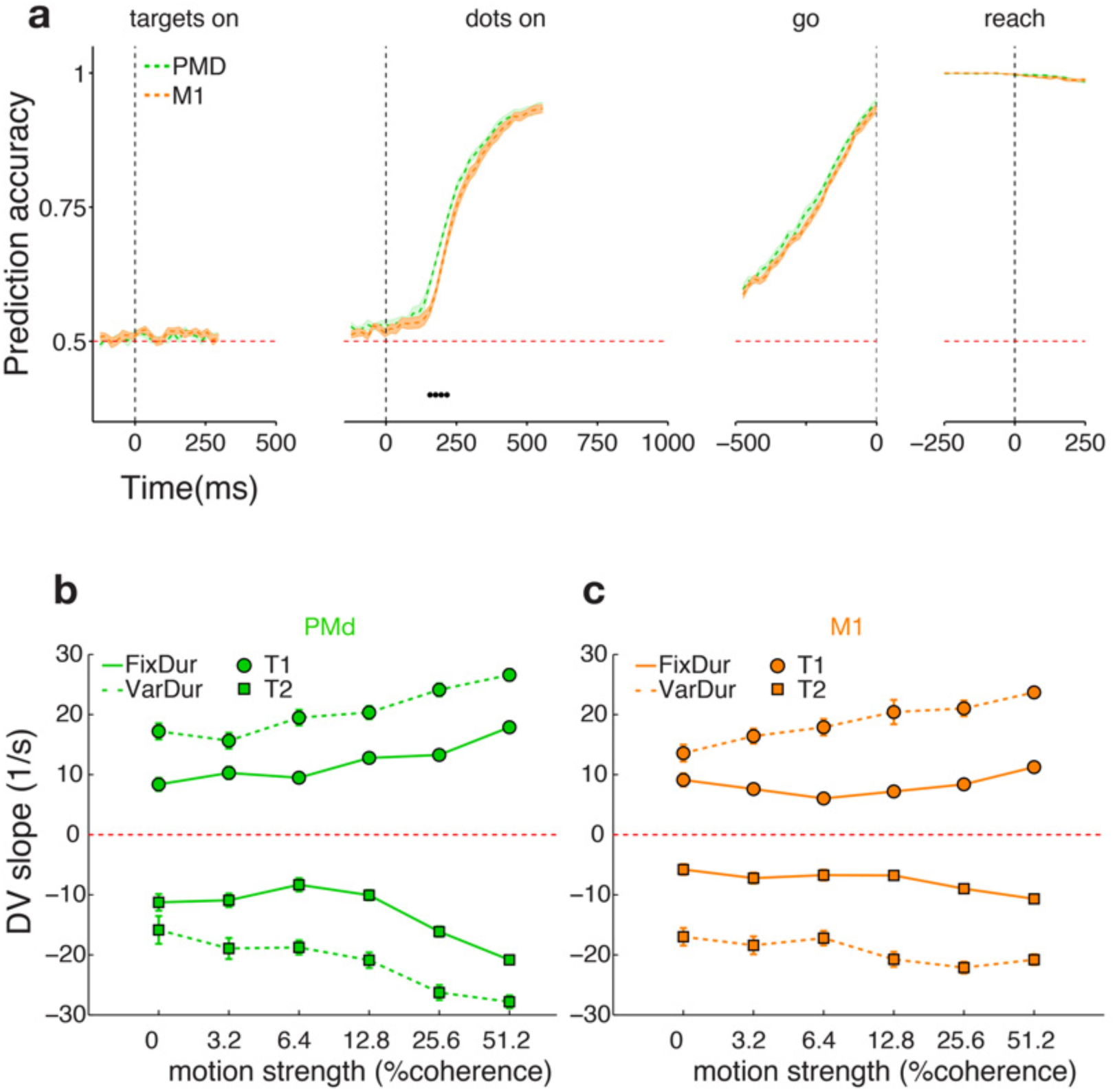
Effects of stimulus duration uncertainty on choice prediction accuracy and model decision variable (pooled results across 2 monkeys). **a) Average prediction accuracy for variable stimulus duration sessions increases during the stimulus presentation.** Equivalent to Figure 2a, but for variable stimulus duration sessions (same figure conventions). In the “dots on” panel, data were only included prior to the offset of the stimulus, ensuring that peri-movement activity did not affect the firing rates in this epoch. Because the visual stimulus varied in duration, fewer trials contribute to the trace as time progresses. In contrast to Figure 2a, prediction accuracy rises much faster and reaches higher values when the stimulus duration is uncertain. Differences between PMd and M1 dynamics are highly reduced under conditions of uncertainty. **b) Single-trial DV slopes for PMd increase for the variable duration task while maintaining co-variation with stimulus coherence.** Data points show average single-trial DV slopes as a function of stimulus coherence and choice for variable stimulus duration (dashed lines) and fixed stimulus duration (solid lines, same data as Figure 2d) sessions. Error bars indicate +/- SEM across trials. **c) Single-trial DV slopes for M1 increase for the variable duration task while maintaining co-variation with stimulus coherence.** Same as **b)** for M1. Figure conventions as in b).

The slope analysis of single-trial DVs in the variable duration task showed that the coherence effects were largely conserved on single trials in both PMd and M1 despite the accelerated dynamics (dashed lines in Fig. 3b-c, Supp. Fig. 8b-c, Supp. Fig. 9b-c, statistical analyses, Supp. Table 1, all parameters, Supp. Fig 10 Supp. Fig 11). The DV slopes are overall larger for the variable duration task compared to the fixed duration task (vertical shift between the solid and dashed lines in Fig. 3b-c, Supp. Fig. 8b-c, Supp. Fig. 9b-c). This difference was significant for all areas and target directions (p<10^−18^, see Methods and Supp. Table 4).

### Changes in decision-related dynamics are not due to contamination by motor signals

The data in Fig. 3b,c suggested that dynamics of decision-related activity accelerated under conditions of temporal uncertainty, as indicated by the overall larger slopes in the variable duration task. Before testing model predictions, we addressed a potential confound that could lead to misinterpretation of the DV slope data. When the duration of the stimulus is uncertain and the operant response is required immediately after offset of the stimulus (i.e. no delay period), it is possible that motor planning is accelerated and that kinematic signals related to the operant movement invade the visual stimulus period, contaminating our DV estimates from PMd and M1 recordings. If the accelerated dynamics were exclusively due to motor preparation signals contaminating the early dots period, we would expect the coherence effects on single trial DVs to diminish or disappear altogether in the variable duration task. This was not the case as shown in Figures 3b-c, Supplementary Figures 8b-c and 9-bc, and Supp. Tables 1-4 (effect of coherence on the DV slopes, p<10^−4^). Nonetheless, we formally tested the motor kinematics hypothesis by measuring the extent to which neural activity during the stimulus viewing period predicts motor kinematic variables in both tasks. Our analysis focused on movement onset time after the Go cue (reaction time, RT) and hand velocity, both of which known to be reflected in the activity of PMd and M1 neurons (Afshar et al., 2011, Kubota and Hamada, 1979, Churchland et al., 2006a, Churchland et al., 2006b). We performed Ridge regression of both kinematic variables onto population neural activity to determine the time when kinematic signals appear in PMd and M1.

For the fixed duration task, our prediction of RT from neural population data was poor in the targets and dots epochs as expected from the task design (Fig. 4 a-b). Only late in the delay period (∼last 50 ms), when the monkey was presumably planning the arm movement, did we observe a very small rise in RT variance explained by neural activity. This rise became significant for both areas and targets within 60 ms after the go cue (Wilcoxon signed-rank test p<0.01, Holm-Bonferroni correction for multiple comparisons). We observed a wide range of RTs in both tasks, which lead to a strong dynamic range in firing rates that correlated with RT after the go cue and thus lead to high R^2^ values, which are expected for (pre)motor brain regions. Crucially, the results for the variable duration task were similar in terms of temporal profile, with significant R^2^ values only present after the go cue but not during dots (Fig. 4 c-d, Supp. Figs 12 and 13 show model performance for example sessions). Repeating the same analysis for hand peak velocity, we observed only modest R^2^ values after the go cue and around the time of the response for both tasks (Fig. 4 e-h). The absence of significant R^2^ values during the stimulus presentation period in our two tasks (Fig. 4 a-h) confirmed that hand motor signals were not contaminating our DV estimates.

**Figure 4.**
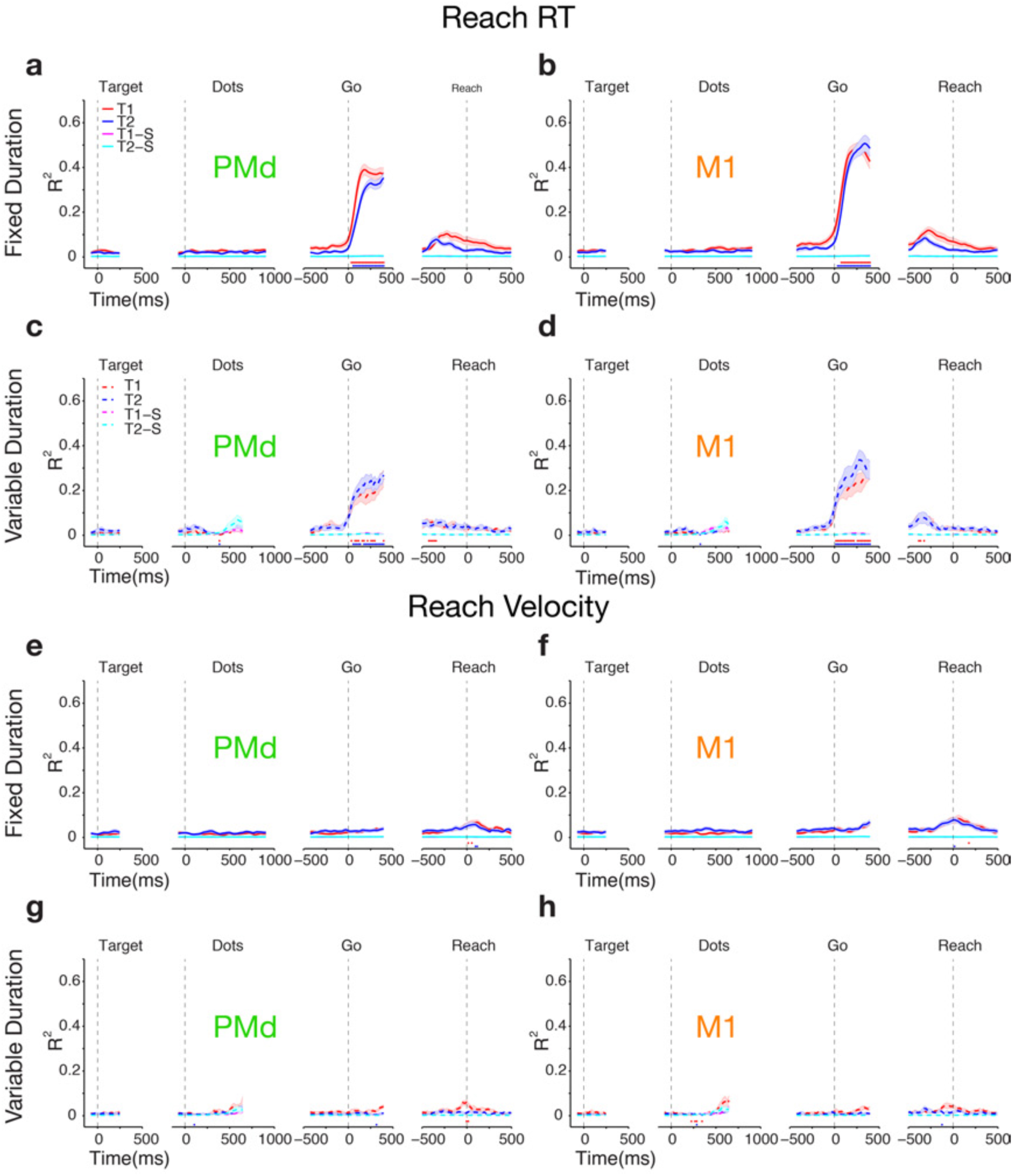
Neural activity in PMd and M1 only becomes predictive of RT and hand velocity around the go-cue in both tasks (pooled results across 2 monkeys). **a) Fraction of variance explained by a linear model regressing unit activity in PMd against reaction time for the fixed duration task only increases after the go cue.** Red traces represent average fraction of variance for rightward choices and blue traces for leftward choices ± SEM (shaded areas). Across the population, neural activity only becomes a reliable RT predictor on or after the time of the go cue. Magenta and cyan lines (highly overlapping) show results from a model trained on shuffled data as a control. Red (Blue) dots above the x-axes denote time points for which the explained variance was significantly different from baseline (defined as the average for [-100, 200] aligned to targets onset) for T1 (T2) according to Wilcoxon signed-rank test (p<0.01 Holm-Bonferroni correction for multiple comparisons). Median RTs for fixed duration task monkey H: 361 ms, monkey F: 424 ms. **b) Fraction of variance explained by a linear model regressing unit activity in M1 against reaction time for the fixed duration task only increases after the go cue.** Same as a) for M1. Figure conventions as in **a)** **c) Fraction of variance explained by a linear model regressing unit activity in PMd against reaction time for the variable duration task only increases after the go cue.** Figure conventions as in **a).** Median RTs for variable duration task monkey H: 335 ms, monkey F: 427 ms. **d) Fraction of variance explained by a linear model regressing unit activity in M1 against reaction time for the variable duration task only increases after the go cue.** Figure conventions as in **a)** **e)-h) Same as a)-d) for hand peak velocity.** Across the population, neural activity is a poor predictor of hand peak velocity throughout the trial; a slight increase in variance explained occurs only around reach initiation.

Finally, and to rule out the contamination from additional variables associated with the eye movement, we performed the same analyses on the analogous saccade parameters: saccade RT and saccade peak velocity. Similar to our results for hand movement kinematics, we could predict a significant fraction of variance of saccade RT only during and following the go cue (but not before) (Supp. Fig. 14 a-d). The R^2^ peaks for saccade RTs were significantly lower than those for hand RTs (Fig. 4 a-d vs Supp. Fig. 14 a-d) (Wilcoxon signed-rank test for peak R^2^ Hand vs Eye RT: p<5e-4 for all areas, tasks and target directions). Further, saccade peak velocity was not explained by PMd or M1 neural data at any point in the trial (Supp. Fig. 14 e-h). The weaker representation of eye kinematics after Go cue is consistent with the expected role of PMd and M1 in controlling arm movements.

In summary, our results showed that regardless of task timing, motor parameter representation in PMd and M1 was reliable only around and after the go cue and not while the visual evidence was presented. Thus, the accelerated dynamics of choice predictive activity early in the stimulus presentation period of the variable duration task was not due to a contamination by motor signals.

### Progressive recruitment of choice selective neurons underlies accelerated dynamics of the DV in the variable duration task

In the previous sections, we showed that the dynamics of choice related neural activity on single trials is flexible, being strongly influenced by the expected statistics of stimulus duration. In conventional evidence integration models of decision formation (Ratcliff and McKoon, 2007, Lo and Wang, 2006, Mazurek, 2003) changes in the dynamics of the DV are implemented through parameters that govern the accumulation of sensory evidence. We tested whether these models could replicate our observation that the DV buildup rates are higher in the variable duration task, and that the size of the increase is independent of motion coherence.

We focused on a simple formulation of integration-to-bound models, in which two competing integrators accumulate noisy evidence about motion energy over time toward a decision bound (Kiani et al., 2014a). As soon as one of the integrators reaches the bound, a choice is made and the two integrators maintain their state (integrated evidence) until the end of the trial (Kiani et al., 2008, Shadlen and Newsome, 2001). We simulated two pools of spiking neurons whose mean firing rates represent the state of integrators. Finally, we trained our logistic classifier on the spike counts of the simulated neurons and calculated the DV buildup rates, just as we did for our recorded neurons.

We envisioned three possible mechanisms for changes in the DV dynamics, each of which can account for the observed psychophysical data with appropriate parameter adjustments (Fig. 5a). The first two are based on known phenomena: urgency (Churchland et al., 2008, Purcell and Kiani, 2016) and sensory gain (Cisek et al., 2009). Urgency is an evidence-independent signal that drives both integrators toward their bounds. In principle, an overall increase of urgency in the variable-duration task might mimic a coherence-independent increase in the DV buildup rates as observed in the physiological data (Fig. 3b,c, vertical shift in DV slope vs. coherence traces) through faster commitment to choice. However, because urgency affects both accumulators, and the DV depends largely on the *difference* in the activity of our two pools of neurons, the effect of urgency on the DV slope is small unless very large urgency signals are imposed (Fig. 5b, note that the cyan, red, and black data points are almost completely superimposed in this figure). A large urgency signal is mathematically equivalent to a reduction in the decision bound and would lead to less accurate choices and, consequently, to a sizeable increase in psychophysical thresholds in the variable duration task (Fig. 5c). Although the data of monkey F provided some support for increased threshold, monkey H was incompatible with the prediction of the urgency model (Supp. Fig. 15).

**Figure 5.**
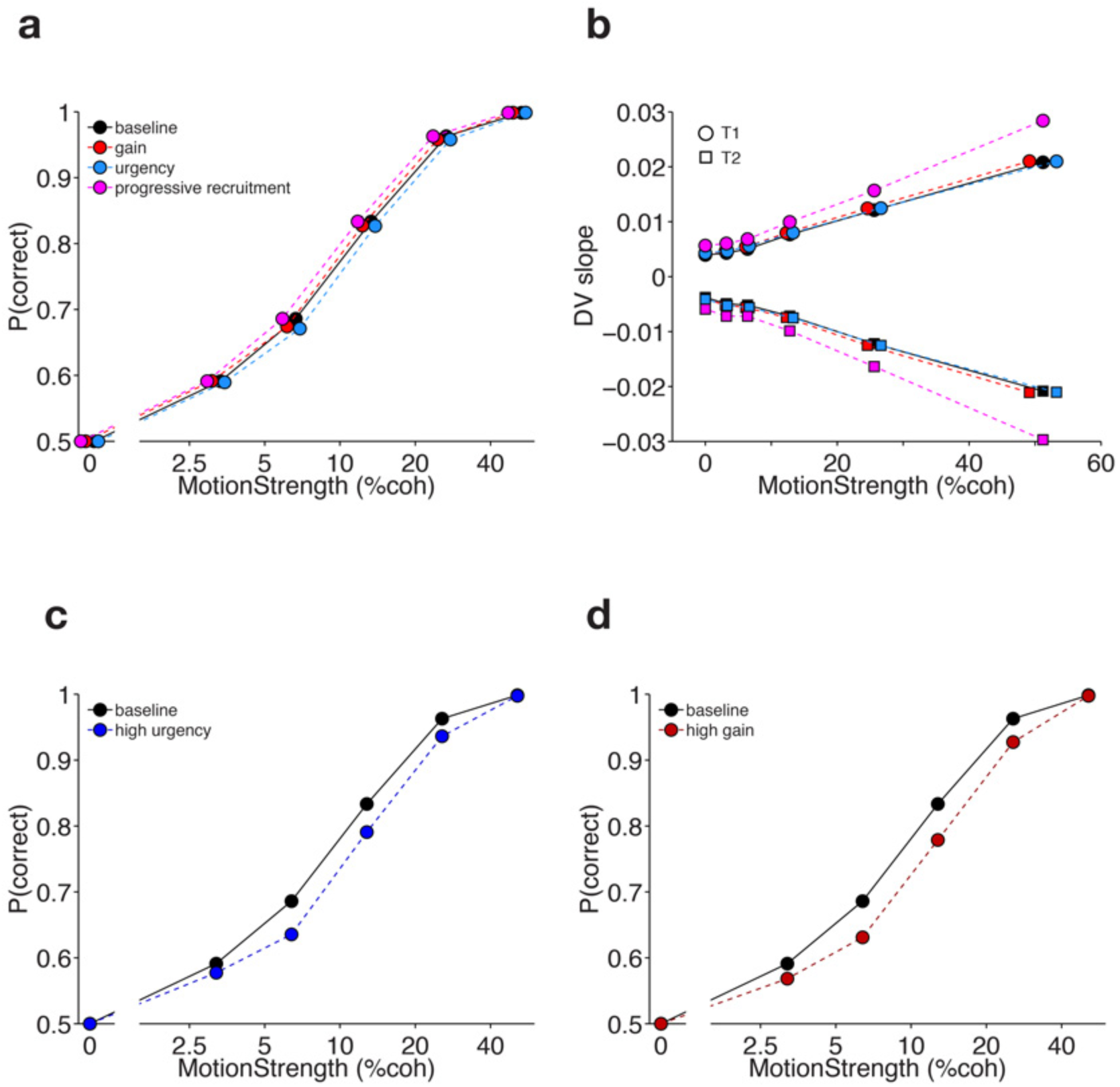
Modeling results suggest that the progressive recruitment model can explain the increase in the DV slopes for the variable duration task. When the parameters of the urgency, gain, and progressive recruitment models are adjusted to match the psychometric function under two task conditions (**a**), only the progressive recruitment model could replicate the increased DV slopes observed in the data (**b**). Because the curves were highly overlapping, data points in (**a**) were offset on the x-axis by multiplying the x-coordinates by a factor c for each curve (*c_baseline_* = 1.04, *c_gain_* = 0.96, *c_urgency_* = 1.08, *c_PRM_* = 0.92) to aid interpretation. Same procedure was performed in (**b**) with *c_baseline_* = 1, *c_gain_* = 0.96, *c_urgency_* = 1.04, *c_PRM_* = 1. Increasing urgency or gain to match the increased DV slopes leads to significant changes in the psychometric function, inconsistent with the observed data (**c** and **d**).

The sensory gain model builds on the proposal that later sensory evidence is progressively amplified with a gain factor before integration (Cisek et al., 2009). The original proposal by Cisek and colleagues assumes a leaky integration process with a short time constant (Cisek et al., 2009) but in our tests the time constant of integration was not a key factor. Similar to increased urgency, increased gain on sensory inputs might lead to a coherence-independent increase in DV slope by accelerating bound crossing and causing earlier commitment to choice. However, just as we showed for the urgency signal, modest increases in gain do not generate a significant change in the DV slopes as these largely depend on the difference between the accumulators, both of which are affected by the increase in gain (Fig. 5b, red data points are largely superimposed on cyan and black). While large gains can increase the DV slope, they also lead to reduced performance accuracy (Fig. 5d) because the bound crossing is accelerated and effective integration time is shortened, similar to what we observed for the urgency signal above. The data from monkey H reduce the likelihood that the urgency or sensory gain mechanisms are the only causes of the accelerated DVs in the variable duration task.

Our third hypothesis proposes that the accelerated dynamics of the DV is due to progressive recruitment of additional neural signals under conditions of temporal uncertainty that represent choice outcome, which we term “categorical choice”, but not the accumulation of sensory evidence that leads to the choice. We postulate that these categorical choice signals appear in each pool of accumulator neurons with increasing frequency as each accumulator nears its bound (see below). In our implementation, the strength of the categorical choice signal varies across single neurons and is independent of the strength of the accumulated evidence signal in each neuron (see Methods, *γ_i_* values). In effect, the weighted contributions of these neurons constitute a subspace of the neural population activity carrying coherence-independent choice signals. This “categorical choice subspace” would not contribute to the formation of the decision but might be necessary for translating the output of the integration process into preparation for a specific operant action. Hereafter, for brevity, we’ll refer to this subspace as the “choice subspace”.

In principle, this choice subspace could be encoded by a population of dedicated neurons that transition from an uncommitted state (baseline firing rate) to a committed state for choice 1 or choice 2, with the transition becoming more probable as each accumulator approaches its decision bound. We, however, favor the alternative implemented in our simulations—the same neurons that represent integration of evidence also represent the categorical choice. This mixed selectivity at the level of single neurons leads to the representation of choice and evidence accumulation in separable subspaces at the level of population responses. These two methods for implementation of the categorical choice signal have similar consequences for the behavior and calculated DVs, but the latter is more in line with previous observations in frontal cortex (Mante et al., 2013).

As suggested above, we simulated this mixed selectivity in a population of model neurons whose responses were weighted sums of accumulated evidence and a categorical choice signal (Methods, Integration Models). The choice signal was a nonlinear monotonic function of the distance of the accumulated evidence from the decision bound and can be thought of as a readout of the accumulation process in preparation for commitment to a choice. We call this the progressive recruitment model (PRM) for representation of choice signals. In the variable duration task, acceleration of the choice signal enhances the representation of the upcoming choice and boosts the model DV, leading to a coherence independent increase in DV slopes (Fig. 5b, note that the magenta points are offset vertically from cyan, black and red).

PRM captures the behavioral data well because accelerated recruitment of coherence-independent choice signals does not cause perturbations in the underlying integration process and does not change psychophysical performance (Fig. 5a). Thus the PRM neatly captures the key behavioral (Fig. 5a) and physiological data (Fig. 5b) in monkey H. In contrast to monkey H, psychophysical thresholds for monkey F increased under conditions of temporal uncertainty, implying changes in the underlying integration process (e.g., increased urgency or sensory gain). Importantly, accelerated choice representation could happen simultaneously with changes in the integration process that could cause an increased psychophysical threshold for monkey F. Overall, our modeling results suggest that accelerated choice representation, either in isolation or mixed with urgency or sensory gain, plays a key role in enhanced response dynamics in PMd and M1 in the variable duration task.

### The progressive recruitment model makes specific predictions at the population and single unit levels

The PRM, as implemented in our simulations, makes specific, testable predictions about the spatiotemporal features of the neural responses in both the fixed and variable duration tasks. First, at the population level, the choice representing subspace should be stable during a trial as more units are recruited to maintain a representation of choice. Such stability facilitates decoding by downstream areas in the presence of timing differences in our tasks. Second, this stabilization should happen faster in the variable duration task due to the accelerated recruitment of choice representing neurons. Third, the choice subspace in the population responses should be shared across the two tasks. Fourth, at the single unit level we should observe the progressive onset of choice representing units, some during the psychophysical integration window (Fig.1d) and some only later in the trial. These units should have stable choice preference (left or right) and stable or increasing choice modulation and their recruitment should be accelerated in the variable duration task.

For simplicity in our simulations, we assumed no categorical choice representation in the fixed duration task (Methods, Integration Models). Similar results would have been obtained, however, if categorical choice signals were also recruited in this task (a non-zero average *γ_i_* parameter) as long as they remained substantially lower than in the variable duration task. We expect this to be a more plausible scenario, and the extent to which progressive recruitment is present in the fixed duration task can be tested empirically.

In the following two sections we test these predictions first at the population level and then at the single unit level.

### Choice signal stabilizes during stimulus presentation in PMd and M1

To test our predictions, we started by examining the structure of the temporal representation of choice across the entire trial. For each time point in each task we defined a “choice axis” that best represents modulation of neural responses with choice. Neurons with strong choice modulation at that time had a large weight in the choice axis and neurons with smaller modulation had smaller weights (see Methods). By comparing the similarity of choice axes at different times and in different tasks, we could quantify the stability of choice-representation in the population. Fig. 6 shows the inner product of choice axes at different times. A large inner product suggests good alignment of the choice axes and high stability of the choice-representing subspace. Conversely, a low inner product suggests a rotation in the choice axis, which could happen if two different sub-populations of neurons represent the choice at different times or if the relative contribution of neurons to the representation of choice changes over time. Mathematically, the projection of a choice axis on itself would be 1, making the diagonals uninformative. We therefore calculated two choice axes for each point in time for two independent halves of each session’s data and measured the projection of these two axes onto each other (see Methods). This way, the diagonal elements of the projection matrix were not set to 1 but instead provided a measure of self-consistency of the choice axis. Armed with these stability and self-consistency metrics we investigated our model predictions.

**Figure 6.**
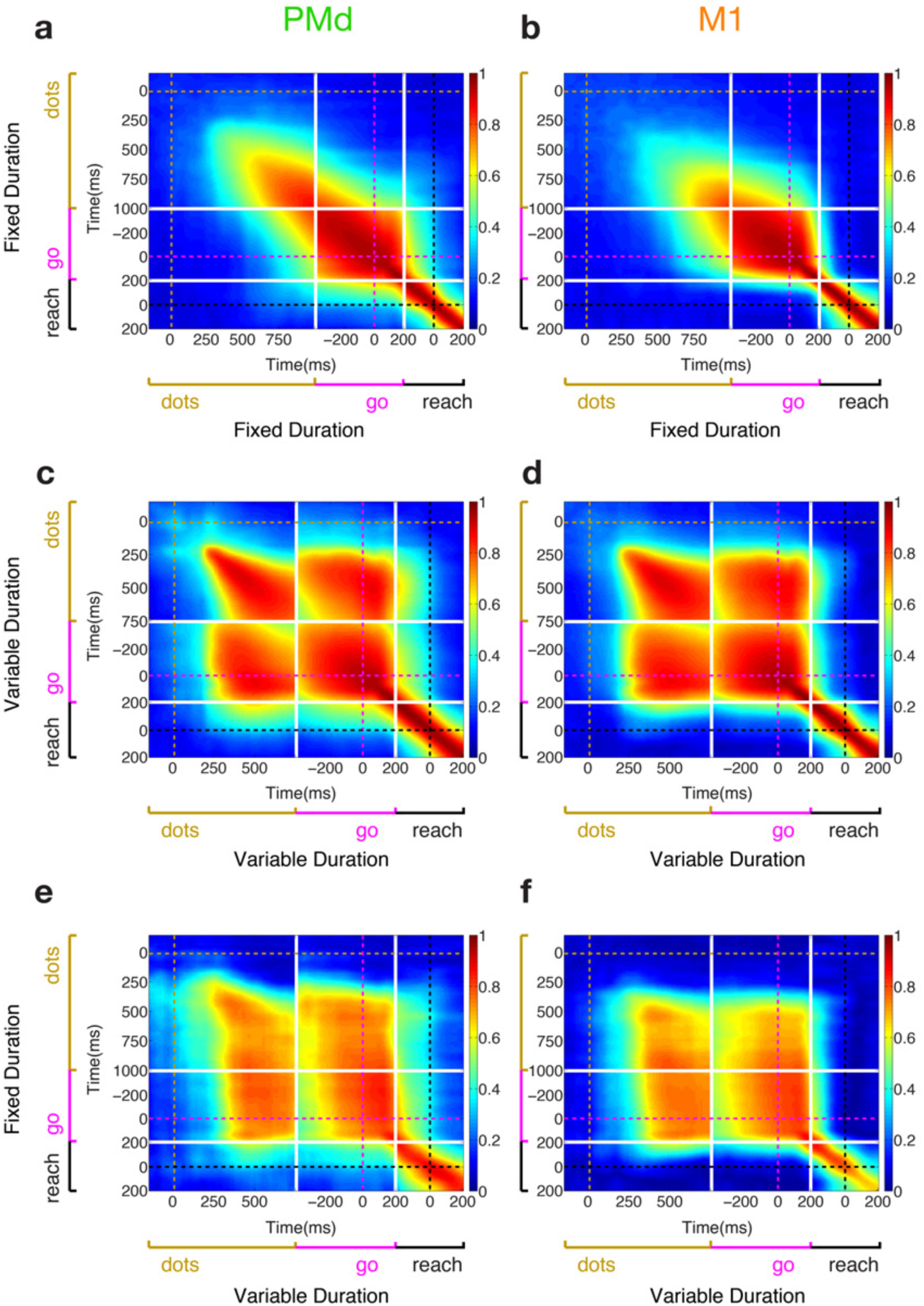
Stability of choice representation during dots is dependent on the statistics of stimulus duration (pooled results across 2 monkeys). **a) Choice representation only becomes stable late in the dots presentation period in the fixed duration task.** Heat map shows the dot product of the choice vector (vector of beta values for choice) across time. Vectors were obtained using 50 steps of 2-fold cross-validation. Warm colors correspond to very high overlap between vectors whereas cool colors denote little projection between vectors. In PMd (left panel) choice representation becomes stable ∼600 ms after stimulus presentation. In M1 (right panel) this phenomenon occurs even later around ∼750 ms. Note how broadly stable the choice signal is during the delay period and how locally stable it is around the reach. White solid lines denote the separation of epochs (dots end, and go cue +200 ms), golden, magenta and black dashed lines mark the dots onset, go cue and reach initiation, respectively. Data from both monkeys. **b)** Same as **a)** for M1. **c) Choice representation becomes stable very early in the dots presentation period in the variable duration task.** Heat map shows the dot product of the choice vector (vector of beta values for choice) across time. The choice signal is already very stable only 300 ms after the stimulus presentation both in PMd (left panel) and M1 (right panel). Unlike in **a)** and **b),** the pre-go period in the variable duration task overlaps with the dots period because of the absence of a delay period; thus correlations in the off-diagonal quadrants should be interpreted with caution since the same data can be correlated against themselves. Figure conventions as in **a)**. Data from both monkeys. **d)** Same as **c)** for M1. **e) Choice representation is stable across tasks.** Heat map shows the dot product of the choice vector (vector of beta values for choice) across time between the fixed duration task (y-axis) and the variable duration task (x-axis). Note that the representation of choice in the second half of the dots presentation and delay period on the fixed duration task strongly overlaps with the early choice representation in the variable duration task. Figure conventions as in **a)**. Data from monkey F. **f)** Same as **e)** for M1.

Starting with PMd in the fixed duration task (Fig. 6a, Supp. Figs 16a and 17a), we observed three important features. First there was a gradual emergence, rotation and stabilization of the choice axis (emergence of square structure in the heat map) that started ∼350 ms after dots onset and unfolded over the remainder of the dots presentation. Second, the dots period was followed by a highly stable choice signal in the delay period. Importantly, the choice axes late in the dots period were largely overlapping with the choice axes early in the delay period (up until the go cue) indicating that the representation of choice was largely maintained even in the absence of additional visual evidence. Third, the choice signal around the initiation of the reach, despite being extremely strong, was also very transient in its direction in neural state space. These three main features were recapitulated for M1 (Fig. 6b, Supp. Figs 16b and 17b), the main difference being the latency for stabilization of the choice axis during the dots presentation, which happened faster for PMd (∼350ms after dots onset) compared to M1 (>500 ms after dots onset). The temporal ordering between the two areas was consistent with our earlier analysis of choice predictive activity in the fixed duration task (Fig. 2a). These results are consistent with the first prediction from the PRM regarding stability of the choice subspace during dots and delay period.

For the variable duration task the rotation and stabilization of the choice axis happened much faster (∼250 ms after dots onset) than in the fixed duration task, consistent with the second prediction of PRM. In the variable duration task, in fact, the temporal stabilization of the choice axis was virtually indistinguishable between PMd and M1 (Fig. 6c-d, Supp. Figs 16c-d and 17c-d).

The heat maps provide a qualitative description of stability of the choice subspace. We quantified stability *within* and *across* epochs using decoding analyses (see Methods). Our results show substantial choice representation stability within the target, dots and pre-go epochs, but not the peri-movement epoch (Supp. Fig. 18 and 19). In addition, our results demonstrate choice representation stability across the dots and pre-go epochs but not across other pairs of epochs (Supp. Fig. 20).

Finally, our results also suggest a stable choice representation across tasks. Taking advantage of sessions in which we recorded the same units in each brain area while the monkey performed both tasks, we compared alignment of choice axes across time and tasks (Fig. 6e-f). The choice axis measured in the stimulus presentation for the variable duration task was largely consistent with choice axes at later times in the fixed duration task (and vice versa), in agreement with the third prediction of the PRM. This implies that the same transformation from integration of evidence to stable choice signal occurs in the two tasks and is being carried out through the recruitment of the same units, only at different rates that reflect the cognitive demands imposed on the subject.

### Stabilization of population choice axes occurs through progressive recruitment of neurons with sustained choice modulation

We next examined the choice modulation at the single unit level to test the fourth prediction of the PRM. This analysis provides a bridge between our population analyses, modeling results, and single unit properties. We first calculated the cumulative fraction of units that display significant choice modulation as stimulus presentation progresses. Consistent with the PRM, the cumulative fraction rises much faster over the course of stimulus presentation in the variable duration task (dashed lines) compared to the fixed duration task (solid lines) for both PMd (Fig. 7a, Supp. Fig. 21a and 22a) and M1 (Fig. 7b, Supp. Fig. 21b and 22b).

**Figure 7.**
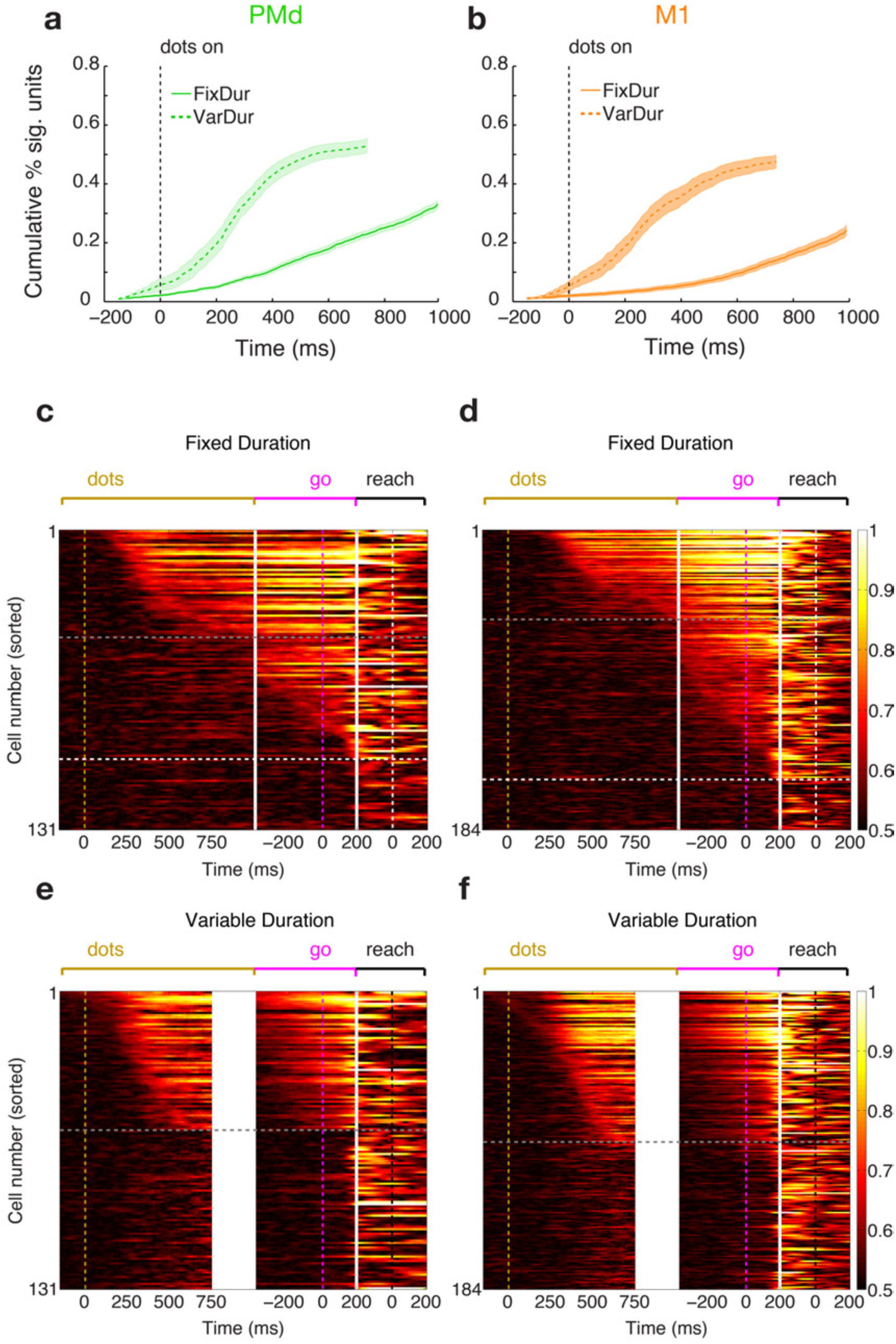
Recruitment of choice predictive cells accelerates for both brain areas under uncertainty conditions. **a) Fraction of units carrying significant choice signals increases faster in the variable duration task.** The cumulative fraction of units with significant choice signals is plotted as a function of time aligned to dots onset for PMd. Solid line shows the results for the fixed duration task and dashed line the variable duration task. Shaded areas denote ± SEM (across sessions). Data from both monkeys. **b)** Same as **a)** for M1. **c) Individual unit choice predictive activity is stable during dots presentation and builds up slower in the fixed duration task.** Area under ROC traces for all units recorded in one session in PMd. Traces we sorted by onset of significant choice modulation during the dots presentation (one row for each unit). White solid lines denote the separation of epochs (dots end, and go cue +200 ms); golden, magenta and black dashed lines mark the dots onset, go cue and reach initiation, respectively. Horizontal dashed gray line separates cells with significant choice modulation during dots (above) from cells with significant choice modulation that only starts during the delay period. Horizontal dashed white line separates the latter group from the remainder of the population (below). Data from monkey F. **d)** Same as **c)** for M1. **e) Individual unit choice predictive activity is stable during dots presentation and builds up faster in the variable duration task.** Figure conventions as in **b)**. Data from monkey F. **f)** Same as **e)** for M1.

Further support for a mechanism of progressive recruitment came from persistent activity of choice-selective neurons. We used area under the ROC (Shadlen and Newsome, 2001) (auROC) to quantify how well single neuron responses represented choice at each time point during the trial. If the representation of choice at the single neuron level is stable over time, a heat map of auROC of individual neurons, ordered by onset of choice representation, should show an upper triangular structure. In contrast, transient representation of choice in individual neurons should be evident as a diagonal structure in the heat map (Harvey et al., 2012, Morcos and Harvey, 2016). Fig. 7c-d showed a strong upper triangular structure for units with significant choice modulation (above the gray dashed line), indicating persistent choice modulation over the course of the trial. The existence of units that only become choice modulated late in the dots period or even during the delay period for both PMd (Fig. 7c) and M1 (Fig. 7d) matches our expectation that progressive recruitment of choice signals is also present in the fixed duration task.

Consistent with our logistic regression results, the emergence of persistent choice representation in the individual units was faster and more widespread in PMd (Fig. 7c Supp. Fig. 23a) than M1 (Fig. 7d, Supp. Fig. 24b) in the fixed duration task.

For a direct comparison across areas and tasks we also calculated the area under ROC traces for sessions in which fixed and variable duration tasks were run in the same experimental session (while putatively recording from the same units, see Methods). The results show that for both areas (Fig. 7e-f, Supp. Fig. 24c-d), units with stable modulation were recruited earlier during the trial, and just as in the fixed duration task, maintained their modulation strength until close to the time of the arm movement. Not only did the same units represent choice in both areas during the stimulus presentation period (Fig. 6e-f), but also their recruitment ordering was consistent across tasks for both areas (Spearman correlation between latencies across tasks for PMD: rho = 0.869, p = 1.37×10^−11^ and M1: rho = 0.579, p = 3.79×10^−5^ for the example session shown in Fig. 7e-f), further suggesting that the same transformation of signals is happening in both tasks at different rates. Finally, for the variable duration task, just as in the logistic regression analysis (Fig. 3a), the differences between the two areas largely vanished in the variable duration task, both in terms of fraction of significant units and rate of recruitment. Taken together these results corroborate the fourth prediction from the PRM and show that temporal stability of choice predictive signals inferred at the population mechanism is present at the level of individual units as well.

### Choice signal is distributed across the neural population

The stability of the choice axis over time (Figs. 6, 7) suggests that there is little relay of information between different ensembles of neurons (sequence mechanism: (Harvey et al., 2012, Morcos and Harvey, 2016, Rajan et al., 2016, Scott et al., 2017)) once the choice signal appears in the PMd and M1 populations. To further test whether a sequence mechanism might be compatible with our results, we quantified the distribution of choice-related neurons in the population as a function of time during the trial. If the choice representation is generated by a sequence mechanism, the neural representations at a given time during the trial should critically depend on a small number of key neurons. Removing these neurons from the population should result in a drastic degradation in the quality of the neural representations (Haxby et al., 2001, Kiani et al., 2007). We tested this possibility by performing a unit dropping analysis that calculates how prediction accuracy is impacted by exclusion of the best units (Kiani et al., 2015).

We illustrate our results by focusing on two points in time: the end of the stimulus presentation period (last 50 ms) and go cue presentation (50 ms before go), because a strong choice related signal is present at these times in both tasks. Our results (Fig. 8a, Supp. Fig. 24a and 25a) show that predictive accuracy decayed smoothly as the best units were removed for both areas and both monkeys. We did not observe any precipitous drop in prediction accuracy that might suggest a special role for a small group of transiently active neurons. PMd remained more predictive than M1 at the end of stimulus presentation (Fig. 8a, Supp. Fig. 24a and 25a), as expected from previous sections of this paper. This discrepancy vanished around the go cue. Also, in the variable duration task the decay in performance was shallower (up to only ∼10% for the best 70 units) compared to the fixed duration task (Fig. 8b, Supp. Fig. 24b and 25b) due to the higher number of strongly tuned units (Fig. 7). The key observation is that representations in both areas and in both time points show remarkable robustness to exclusion of the best predictive single units. Even after dropping 70 best units (corresponding to a median 46%/33% of units in PMd and 48%/31% of units in M1 in the fixed/variable duration tasks respectively) choice prediction accuracies remained well above chance.

**Figure 8.**
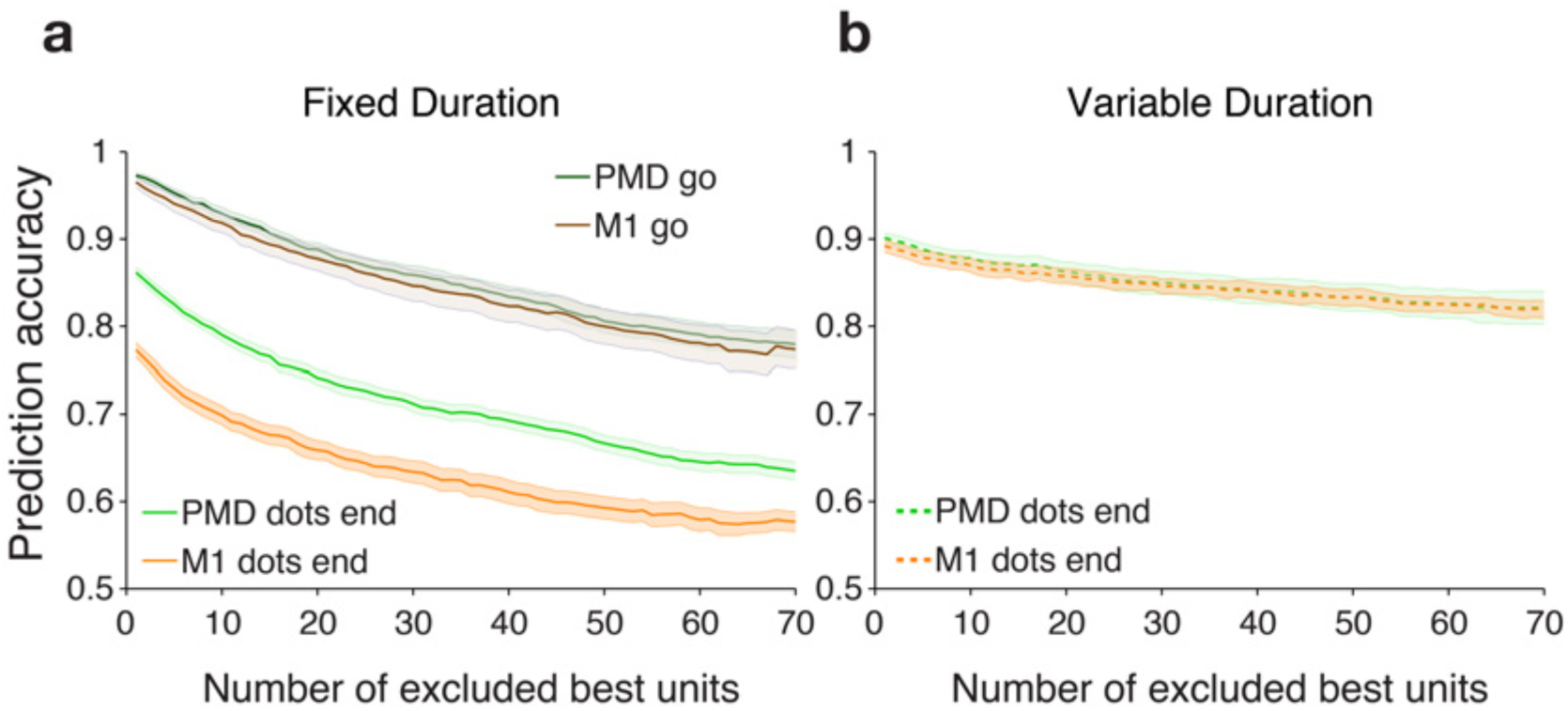
Choice signal is robust and distributed across the population of cells in both areas and both tasks (pooled results across 2 monkeys). **a) Choice signal is robust in the fixed duration task.** Average prediction accuracy curves for PMd (green) and M1 (blue) ± SEM (shaded areas) as a function of the number of best units excluded for the fixed duration task. The unit dropping curves were calculated for two separate time points: end of dots presentation (light shades) and go cue presentation (dark shades) Curves were calculated for each session/area separately and then averaged across sessions. The decay in performance is smooth, demonstrating that the choice signal is distributed across many cells. As expected from figure 2 a), the initial accuracy for the go cue period is higher than for the end of dots. **b) Choice signal is extremely robust in the variable duration task.** Same as **a)** for variable duration task. The choice signal is even more robust in this task as evidenced by the very small decline in prediction performance (<10%) after dropping the 70 best units. Figure conventions as in **a)**. The end of dots and go cue coincide in this version of the task so only one curve is shown for each area.

## Discussion

The primary goals of this study were: 1) investigate whether single-trial, decision-related activity in PMd and M1 has the properties of a DV and determine whether and how DV dynamics depend on uncertainty about stimulus timing, 2) test computational models to identify mechanisms that can explain the observed choice behavior and neural dynamics, and 3) examine the spatio-temporal features of decision-related signals, under conditions of both temporal certainty and uncertainty, with a specific goal of differentiating between stable vs. sequential representations of choice-predictive signals. To achieve these goals we employed two variants of a classical motion discrimination task, fixed and variable duration (Kiani et al., 2008, Shadlen and Newsome, 2001), and combined them with multi-electrode recordings and decoding techniques to obtain reliable single trial estimates of decision-related dynamics at the level of the neural population.

### Population neural activity in PMd and M1 exhibit properties of a decision variable

Our neural population data are consistent with predictions of classic accumulation-to-bound models of the decision process. Specifically, choice predictive activity emerges quickly after stimulus onset in both PMd and M1 and increases with time and stimulus coherence as expected from evidence accumulation (integration) linked to the sensory stimulus. Critically, our simultaneous population recordings provided statistical power to test these predictions on single-trial activity as opposed to trial-averaged activity as in most previous studies. The build-up of choice predictive activity on single trials—as captured in the rate-of-rise of the logistic decision variable—varied systematically with stimulus coherence (Figs. 2d,e, 3b,c; Suppl. Tables 1-3) as expected from accumulator models (Shadlen and Newsome, 2001, Roitman and Shadlen, 2002, Bollimunta et al., 2012) and inconsistent with step like transitions in the neural states (Latimer et al., 2015, Miller and Katz, 2010).

Choice-predictive activity was present in both PMd and M1 even when action initiation was cued more than one second after termination of the visual stimulus. The most pronounced difference between the two areas occurred in the fixed duration task: significant choice related activity emerged faster and was stronger in PMd compared to M1 (Fig. 2b,c). These differences are consistent with the standard view of a greater cognitive role for PMd compared to M1 (Cisek and Kalaska, 2005, Coallier et al., 2015, Wise et al., 1997), and with the idea of a rostro-caudal gradient of visuomotor responses in PMd/M1 (Cisek and Kalaska, 2005) with stronger motor signals in caudal PMd/M1 and stronger target selection signals in pre-PMd/ rostral PMd.

This difference, however, essentially vanished in the variable duration task. Following stimulus onset, prediction accuracy increased at nearly identical rates in the two areas and plateaued at similarly high levels, the only difference being ∼20 msec longer latencies in M1 (Fig. 3a). The dynamics accelerated in both areas under conditions of temporal uncertainty, but the change was particularly dramatic in M1 (compare Fig. 2a with Fig. 3a). These results could not be explained by displacement of motor kinematic signals into the stimulus presentation period in the variable duration task (Fig. 4). Importantly, the accelerated dynamics were independent of the coherence of the visual stimulus; the DV slope vs. coherence curves for the variable duration condition are essentially vertically offset copies of those for the fixed duration condition.

To our knowledge only one other study (Shadlen and Newsome, 2001) employed both fixed and variable duration motion discrimination tasks while recording decision-related activity in individual units. Similar to the current study, the authors observed a larger and faster average increase in choice modulation in LIP neurons in the variable duration task, which they speculated could reflect increased urgency to make quicker decisions when the duration of the sensory evidence is uncertain. While this intuition is appealing, it begs the question as to the actual mechanism underlying the accelerated dynamics, which we explored through a series of quantitative models.

### Progressive recruitment of choice-selective neurons

The discovery of single-trial, coherence-independent acceleration of DV dynamics under conditions of temporal uncertainty provides a useful new constraint on mechanistic models of the decision process. Different variants of race models between accumulation processes have long been proposed to explain both behavior (Beck et al., 2008, Link, 1992, Vickers, 1970, Wong and Wang, 2006) and neural activity in premotor structures (Churchland et al., 2008, Hanks et al., 2014, Scott et al., 2017) during perceptual decision tasks. In this study we implemented three variants of race models that attempted to explain both the observed psychophysical behavior and the single trial DV dynamics across both tasks. The increased gain and increased urgency models could not replicate the DV dynamics for the variable duration task without unacceptable deterioration in psychophysical performance. In contrast, our novel progressive recruitment model (PRM) with an adaptable recruitment rate closely matched both the physiological and behavioral data. This model proposes that PMd and M1 population activity reflect recruitment of a second, coherence-independent “choice” signal in addition to the well-known coherence-dependent signal. Importantly, the PRM and accumulation-to-bound models are not incompatible. PRM simply adds a twist to how choice is represented while evidence accumulation proceeds during a trial.

### Temporal stability and latency of choice representation in PMd and M1

The analyses of choice axes suggested a stable representation of choice during the dots and delay period. However, we found that the choice axis early in the dots period (∼250 ms after dots onset) does not perfectly overlap with choice axis late in the dots period (∼750 ms after dots onset). Through our simulations, we posited that changes in the choice axis between the early and late epochs of the dots period occur due to the recruitment of signals associated with the categorical choice in addition to signals associated with the accumulation of evidence. Similarly, but to a much larger degree, the choice axis during the actual arm movement did not overlap with the choice axis from the dots period, probably reflecting the additional recruitment of signals associated with moving the arm. We believe that the shift in the choice axis across epochs is evidence for the existence of multiple choice subspaces in PMd/M1 (and other brain regions) that are engaged at different epochs in the tasks presented here (and for other tasks). In this study, we have exposed one aspect of these choice subspaces. Multiple choice subspaces will likely reflect the different behavioral demands for the monkey at different points in the task such as sensory evidence evaluation, motor preparation, movement execution, post-movement evaluation, reward expectation, and learning. Reorganization of neural activity into different subspaces has been previously observed in PMd for delayed reach tasks between the movement preparation and execution phases (Elsayed et al., 2016) and is compatible with the low projection between peri-movement and delay period axes we obtained in the fixed duration task (Figure 6a).

We also observed a difference in latency for choice representation to become stable between PMd and M1, and these latency differences depended on the task. For the variable duration task, the latencies for stabilization of the choice representation in both PMd and M1 were well within the estimated psychophysical integration windows for both monkeys (500-600 msec—Fig. 1d). In contrast, M1 data in the fixed duration task appear to stabilize at ∼750 msec (Fig. 6b), well outside the psychophysical integration window. Thus, the M1 delay can be highly variable depending on the expected time for the execution of motor action. In the variable-duration task, where the Go cue can happen any time, M1 responses reflect the choice much earlier. Note that similar progressive recruitment of the choice representing subspace in M1 and PMd would lead to similar reduction of latency in the two areas. Therefore, it is unlikely that a common input to the two areas underlies our results. An appealing hypothesis is that changes of latency in M1 are caused by changes of PMd dynamics. If the choice representation in PMd should reach a threshold level before it emerges in M1, the accelerated choice representation in PMd would cause both accelerated dynamics and significantly reduced latency of choice representation in M1 in the variable duration task. Overall, our results hint at a mechanism where PMd responses lead and furnish the choice representation in M1.

### Progressive recruitment accounts better for our data than a sequence hypothesis

Our results suggest that progressive recruitment of units with temporally stable choice modulation is a plausible mechanism for choice representation in PMd and M1. In contrast, evidence from recent optical imaging studies in rodents (Harvey et al., 2012, Morcos and Harvey, 2016) suggests an alternative mechanism: representation of choice by transient ensembles of neurons that are activated sequentially as the trial proceeds, effectively passing choice information from one ensemble to the next throughout a trial. Intrigued by this finding in rodents, we analyzed our neural population data to test the predictions of these two mechanisms on individual sessions in monkeys. Our analyses of the temporal stability of choice axes, within and across epoch decoding, and unit dropping all support a stable choice representation mechanism over a sequence mechanism. Our failure to detect sequences during the visual stimulus and delay periods does not reflect a problem with our analysis techniques; sequences of ensemble activity were strikingly present in the peri-movement interval for the operant arm movement, as shown by the diagonal structure in the lower right portion of each plot in Figure 6. This diagonal structure results from fast modulations around movement onset that are expected because of impending changes in limb position and kinematics during movement, as reported before (Churchland et al., 2010, Churchland et al., 2012). At the individual unit level our results also matched the progressive recruitment model predictions: the choice signal is carried by a large (and growing) fraction of neurons (Fig. 7a,b) and their modulation is largely stable over the stimulus presentation and pre-go cue period (ROC analyses, Fig. 7c,d).

The pronounced difference between stable choice representation in the primate cortex and sequential representation in the rodent cortex might simply reflect a species difference in neural mechanisms underlying choice behavior. However, a recent study of choice mechanisms in rodents supports stable accumulation of evidence in parietal cortex (Hanks et al., 2015). The key difference between the latter study and those that yielded evidence for sequences is that animals were actively locomoting on a track ball when sequences were observed. A more recent—and as yet not peer reviewed—study suggests that sequences of neuronal activity during track ball locomotion result not from choice-related signals per se, but from specific combinations of bodily position and head angle at successive times during locomotion (Krumin et al., 2017). Our best reading of the current literature is that the evidence for stable representations of choice in primate cortex is strong, whereas the sequence hypothesis that has emerged from rodent work requires further study to confirm, refine, or reject. Developing behavioral tasks that are as similar as possible for monkeys and rodents may help resolve some of these issues.

### Concluding remarks

We have focused on single trial estimation of decision variables in neural population data and development of mechanistic models that explain both the behavioral and physiological data. Like many studies in the contemporary literature, our comparison of models to data relies on regression analyses that produce vectors of weights on the responses of individual units, be they neurons or voxels. Importantly, we do not assert that a downstream brain area or deeper cortical layer (Chandrasekaran et al., 2017) actually performs a linear weighting of PMd/M1 activity in superficial layers to guide decisions. For present purposes, we simply use the DV as a proxy for the informational content about choice present at any given moment in these neural populations (Kiani et al., 2014b). Recordings across multiple brain regions and precise knowledge of projection pathways between them will be required to elucidate the actual mechanisms that transform this information to signals that trigger an action.

More broadly, our study integrates a small but growing body of literature that leverages simultaneous electrophysiological population recordings to explore the neural substrate of internal cognitive phenomena at the level of single trials (Bollimunta et al., 2012, Kaufman et al., 2015, Kiani et al., 2014b, Rich and Wallis, 2016). This approach promises to shed light on internal cognitive processes whose dynamics vary substantially from trial to trial (e.g. decision-making, attention). In the future, the strengths of this approach will be amplified by monitoring cognitive/attentional states in real time and probing the subject and circuit in a neurally contingent manner.

## Acknowledgments

We would like to thank Mr. Julian Brown for support with the data acquisition setup, Ms. Sania Fong for providing help with primate training and handling and Mr. Bora Erden for his work spike-sorting multiple electrophysiological datasets. We would also like to thank all other members of the Newsome Lab and Shenoy Lab at Stanford University for helpful discussions on the methods and results throughout the execution of the project. D.P. was supported by the Champalimaud Foundation, Portugal, and Howard Hughes Medical Institute. R.K. was supported by Simons Collaboration on the Global Brain grant 542997, Pew Scholarship in Biomedical Sciences, National Institutes of Health Award R01MH109180, and a McKnight Scholars Award. C.C. was supported by an NIH/NINDS K99/R00 grant NS092972. K.V.S. was supported by the following awards: NIH Director’s Pioneer Award 8DP1HD075623, Defense Advanced Research Projects Agency (DARPA) Biological Technology Office (BTO) “NeuroFAST” award W911NF-14-2-0013, the Simons Foundation Collaboration on the Global Brain awards 325380 and 543045, and the Howard Hughes Medical Institute. W.T.N. was supported by the Howard Hughes Medical Institute.

## Author Contributions

All authors contributed extensively to the conceptualization of the study, the experimental design and choice of methods for data analysis. D.P. trained animals, performed all electrophysiological experiments, collected and analyzed data. D.P., R.K. and W.T.N. wrote initial draft of the paper. S.I.R, D.P and R.K. performed the surgical procedures. All authors contributed analytical insights and commented on statistical tests, discussed the results and implications, and contributed extensively to the multiple subsequent drafts of the paper.

## Methods

### Subjects

Our experiments were performed on two adult male macaque monkeys (*Macaca mulatta*) trained to perform a direction discrimination task with reaching movements of the arm as operant responses. Neural activity was recorded from populations of neurons in dorsal premotor and primary motor cortex while monkeys performed the task. All training, surgery, and recording procedures conformed to the National Institutes of Health Guide for the Care and Use of Laboratory Animals and were approved by Stanford University Animal Care and Use Committee.

### Apparatus

Monkeys sat in a primate chair in front of a video touchscreen, with their head restrained using a surgical implant. The front plate of the chair could be opened, allowing the subjects to reach the touchscreen with the arm contralateral to the implanted hemisphere. The ipsilateral arm was gently restrained using a delrin tube and a cloth sling. Stimuli were shown on the video touchscreen (ELO Touchsystems 1939L), which allowed us to track hand position at 75Hz and was positioned approximately 35 cm away from the monkeys’ head. Eye position was continuously tracked with an optical eye tracker at 1kHz (EyeLink 1000, SR Research, Canada).

### Motion Discrimination Task

The task employed is a variation of the classical dots discrimination task, in which arm movement was the operant response (Fig. 1a). We used two variants of this task that differed based on the stimulus duration employed. The first version was a classical fixed duration task, in which every stimulus presentation lasted 1000 ms. We termed this version the fixed duration task. In contrast, we also employed a version in which the duration of the stimulus presentation varied from trial to trial. The stimulus duration ranged from 200-1000 (median 435 ms) and it was randomly chosen on each trial by sampling an exponential distribution. We termed this version, the variable duration task. For all variants, the trial starts with the onset of a fixation point (FP; 1.5 degree diameter) on a video touchscreen (Fig. 1a). To initiate the task, the monkey was required to maintain both eye and hand fixation within +/- 3 degrees of the FP as long as it remained on the screen. Importantly, throughout the entire trial, the monkey was required to always maintain direct hand contact with the screen, otherwise the trial would be aborted.

After 300 ms of fixation, two targets (1.5 degree diameter) appeared on opposite sides of and at same distance from the FP. After a 500 ms delay the random dot stimulus was presented for either 1000 ms (fixed duration) or 200-1000 ms (variable duration), depending on the task variant, after which it was removed from the screen. On each trial a fraction of the dots moved coherently along the horizontal axis in the 0 and 180 degree directions. The monkey was asked to report the net direction of motion by reaching to the target in the corresponding direction. The difficulty of the task was adjusted by changing the fraction of dots moving coherently in one direction (motion strength) (Britten et al., 1992). After the stimulus offset the monkey either entered a delay period during which it was required to withhold his response for 400-900 ms (for the fixed duration task) or was immediately presented the go cue (variable duration task). The go cue was then signaled by the offset of the FP at which point the monkey was free to gaze anywhere and report his decision by reaching to one of the two targets. Although gaze was monitored, reward acquisition depended solely on reaching to the correct target. Finally, for a response to be considered valid, the monkey was required to hold its hand position within +/- 4 degrees of the center of the target for 200 ms. The monkey was then rewarded with a drop of juice for correct choices and given a timeout (2-4 seconds) for incorrect ones. Zero coherence trials were rewarded randomly with a probability of 0.5 since there was no correct response on these trials.

### Random dots stimuli

The stimuli used in our psychophysical experiment were random dot kinematograms (RDK) generated using MATLAB and Psychophysics Toolbox. Stimuli were presented on a 19-inch LCD touch monitor (Elo Touchsystems) with 75 Hz frame rate and 800 x 600 pixels resolution positioned 30 cm away from the monkey. The details for generating the random dots stimuli have been described previously (Kiani et al., 2008). We used the same algorithm and parameters except: (1) the stimulus duration was fixed at 1 s for the fixed duration task and variable from 0.2s -1s (exponentially distributed) in the variable duration task; (2) the diameter of the stimulus aperture was 14 degrees, and (3) the speed of the coherent dots was 8 degrees / second. The dot density was 16.7 dots/deg^2^/s, and the dot size 2 pixels. The center of the dots stimulus was situated 12 degrees above the center of the fixation point. To create the impression of motion, the dots in the RDK were split into 3 consecutive sets with the same number of elements (1 set displayed for each individual frame) and displaced 3 frames (40 ms) later. The fraction of dots displaced coherently toward one of the two targets was determined by the coherence (motion strength), with the remaining dots being displaced randomly. For both monkeys, the motion strength could take one of 6 possible values: 0%, 3.2%, 6.4%, 12.8%, 25.6% and 51.2%. The direction and coherence of the motion were randomly assigned on each trial by sampling from a uniform distribution with replacement. For zero-coherence stimuli all dots were displaced randomly but, due to the stochasticity of that process, one obtains non-zero net motion toward the targets over a small number of frames.

### Behavioral Training

Training two monkeys to perform all versions of the dots discrimination task with excellent behavior required a thorough operant conditioning protocol. The protocol had to be adapted to the individual monkeys since they had very different training histories: monkey H was a naive monkey whereas monkey F had been trained on a saccade version of the motion discrimination fixed duration task. Monkey H started by being rewarded just for touching the touchscreen and then gradually progressed to an instructed reach task and from there to a delayed reach task. Once he was proficient in using the touchscreen, the dots stimulus was introduced, cueing the correct target to reach to at the end of the trial. Only easy coherences were used at first, with lower and lower coherences being introduced gradually until the final set was used. The final component of training was eye fixation. Eye fixation was trained by introducing blocks of trials for which the front plate of the primate chair was closed, cueing the monkey to perform the task with eye movements. The fixation window size was gradually decreased, and then eye fixation was also required during the reach blocks. By aborting trials if eye or hand fixation was broken the subject learned that both were required to perform the final task. Monkey F on the other hand was already proficient at discriminating motion so the main focus of training was achieving proficient use of the touchscreen with his hand. The same initial sequence of steps was used to train monkey F to perform delayed reaches. From that point on, the training was focused on combining knowledge about the dots task with the reaching response. Coherences were also introduced sequentially from highest to lowest but at much faster pace compared to monkey H. Recording sessions started when good psychophysical performance was achieved.

### Behavioral Analysis

Psychophysical performance was assessed in two ways: by describing the percentage of correct choices as a function of (unsigned) stimulus coherence and by describing the percentage of right choices as a function of signed stimulus coherence.

The percentage of correct choices as a function of motion strength (stimulus coherence) was fit by a cumulative Weibull distribution function (equation 1):

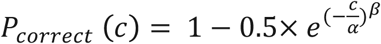

where *P_correct_* is probability correct, *c* is motion strength, *α* is the psychophysical threshold (the value of *c* that corresponds to ∼82% correct responses), and *β* is a parameter that controls the shape of the function, especially its steepness.

The percentage of rightward choices, *P_right_,* as a function of motion strength and direction was fit by a logistic regression (equation 2):

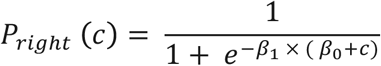

where *c* is signed motion strength, *β_1_* is the slope parameter and, *−β_0_* is the motion strength corresponding to the indifference point. This value was used to assess the monkey’s behavioral bias on each session.

To analyze performance as a function of stimulus duration (Fig. 1d) trials in the variable duration task were ranked by stimulus duration, binned and fitted with Weibull curves. In the y-axis we plot the *α* (threshold) parameter (log_2_ scale) obtained from the fits for each set of trials, and in the x-axes the median duration of the stimulus for each bin (log_2_ scale).

For a perfect integrator threshold should decrease linearly with stimulus duration (in log_2_ vs log_2_ plot) with a slope of -0.5 (Kiani et al., 2008). To assess at what point the observed decrease in threshold deviated from that expected for the perfect integrator we performed bi-linear fits to the data. We forced the first slope to be -0.5 and let the intercept, the second slope and the time of slope change as free parameters. The time of slope change obtained from the fits indicates the point at which behavioral improvement deviates significantly from the perfect integrator prediction. For the regression analyses we used independent bins of 500 trials. For Figure 1d we show bins of 500 trials that are incremented by 250 trials to interpolate the data and guide the eye.

In addition to psychophysical performance two behavioral metrics related to the arm reach itself were also quantified: reaction time (RT) and hand velocity. To obtain precise measurements of reaction times and maximum hand velocity we used the raw hand position data on each trial. We started by up-sampling the raw data by a factor of 13 to obtain artificial 1 ms resolution (since it had been acquired at 75Hz). Then we smoothed the up-sampled data by performing local linear regression to obtain smooth hand traces for each trial. The instantaneous velocity was calculated as the norm of the sum of vertical and horizontal speed components (the instantaneous derivative of the position). The peak hand velocity was calculated for each trial and reaction time was determined as the interval between the presentation of the go signal and the time point at which 20% of the peak velocity was reached.

### Electrophysiological recordings

Two multielectrode arrays (Blackrock Microsystems, Utah) with 96 electrodes each (1mm long platinum-iridium electrodes, 0.4 mm spacing, impedance 400 kOhm) were implanted in primary motor and dorsal premotor cortex of each monkey (Figure 1e). The arrays were placed anterior to the central sulcus, posterior to the arcuate sulcus and lateral but near the superior pre-central dimple (Churchland et al., 2010). Prior to the array implantation, single electrode recordings were performed (FHC, Maine) by lowering dura-piercing electrodes (tungsten, average impedance 6 MOhm) through burr holes, to determine the best location for the arrays. M-L position was determined by performing muscle palpation during recordings and searching for a strong upper arm representation; A-P position was determined by strong perimotor/delay activity in a delayed reach task for M1/PMd, respectively. The coordinates for the best sites were calculated with respect to the center of the chamber and verified during surgery using stereotaxic measurements. These coordinates were used to determine the final location of the arrays, subject to anatomical constraints (curvature of the cortex, blood vessels etc). Continuous neural data were acquired and saved to disk from each channel (sampling rate 30 kHz) and thresholded at -4.5 RMS. Waveforms corresponding to threshold crossings were sorted offline (Plexon Inc., Dallas) using both semi-automatic clustering methods and manual sorting. For all analyses presented in this study we did not differentiate between single-units and multi-units. Our goal was to maximize population predictive power and spatial coverage of the cortex and not just to select the very best isolated single-units. Only units with an average firing rate of 2 spikes/s or more during dots presentation were analysed in this study. The number of units meeting this criterion in each experimental session typically ranged from 100-180 per array.

### Datasets

For each task version and monkey we analyzed all datasets from each brain area that met two behavioral inclusion criteria: 1) over 750 trials and 2) a behavioral bias (|β0|) under 5%, as determined by a logistic regression fit. These criteria were imposed to guarantee that we have a sizeable number of trials per condition (6 coherence x 2 directions = 12 conditions) and that the behavior of the monkey is minimally biased, such that both neural and behavioral results are more easily interpretable. These criteria resulted in a selection of 9 (12) sessions of the fixed duration task and 6 (5) sessions of the variable duration task with no delay for monkey H (F), respectively. Data from both areas were collected simultaneously and the same recording sessions were used.

### Peri-stimulus time histograms (PSTHs)

PSTHs were generated by aligning spike trains of each trial to relevant task events: target onset, stimulus onset, go cue, and movement initiation. These spike trains were then convolved with a Gaussian kernel with a 50 msec acausal and a 50 ms causal component. The standard deviation of the Gaussian used was 30 msec. The resulting spike density functions were then sorted by experimental condition. Once the trials were selected for the specified condition, their spike density functions were averaged.

### Logistic Regression

For each session, the responses of all neurons in 90% of the trials were fit with a logistic model that attempted to separate rightward (T1) and leftward (T2) upcoming choices. The logistic model was fit in 50 ms windows, advanced in 20 ms steps over the entire trial duration (equation 3).

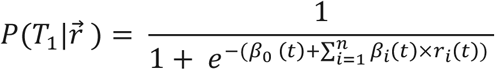

Where 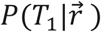 is the probability of observing a particular behavioral choice (*T_1_* or rightward choice in this case) given the population response 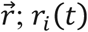 are the z-scored summed spike counts for each neuron and time window, *β_0_* is an intercept term and *β_i_(t)* are the classifier weights (one for each neuron and time window).

The remaining 10% of the trials were tested using the previously trained model and its accuracy was recorded. The same process was followed 10 times for each window (10-fold cross-validation) and the percentage of correctly predicted choices recorded. This process was repeated for consecutive windows displaced by 20 ms and yielding a prediction accuracy trace for each session and brain area. Both correct and error trials were included in this analysis to assure there would not be an imbalance between high coherence trials (more likely to be correct trials) and low coherence trials, which would bias the classifier to perform better on high coherence trials.

An L1- regularization technique (LASSO) was used to constrain the norm of the beta coefficients fitted by the model to prevent over-fitting (Kiani et al., 2014b). The lambda parameter that determines the strength of the penalty for the L1 norm was calculated for the 50 ms window preceding the go-cue by sweeping through 25 potential values and selecting the value with lower deviance by running 10-fold cross validation. This lambda value was then used for the model for all time points.

The exact same procedure was also followed using 150 ms windows (instead of 50 ms) to test whether choice prediction accuracy could still further improve, when applying an identical logistic regression method to the same datasets.

Finally, a slightly different procedure was used when training a single classifier over an entire epoch. The four epochs used for training the four corresponding classifiers were:

- Targets epoch: [-150, 350] ms aligned to targets onset;
- Dots epoch: [150, dots offset] ms aligned to dots onset. Dots offset was 1000 ms for fixed duration task and between 200-1000 ms for the variable duration task;
- Delay/Pre-Go epoch: [-600, 0] ms aligned to go cue;
- Peri-movement epoch: [-200, 600] ms aligned to reach;

All valid 50 ms samples of neural data during the selected period (above) for each epoch were used as a sample to train the corresponding classifier. The classifier was trained on 90% of the trials and tested on 10% of the trials using 10-fold cross-validation. As before, LASSO regularization was used to prevent over-fitting. The regularization parameter lambda was calculated individually for each epoch through cross-validation and chosen as the value with minimum expected deviance. Accuracy was calculated as fraction of test trials correctly predicted at every 50 ms long window (stepped in 20 ms increments).

### Coherence effects on prediction accuracy

For each dataset, coherence effects were assessed by measuring the difference in prediction accuracy between consecutive coherence levels: (51.2%-25.6%), (25.6%-12.8%), (12.8%-6.4%), (6.4%-3.2%), (3.2%-0%). Five 200 ms long periods during the dots presentation were considered. For each period, the five differences in prediction accuracy were averaged across time. Results for each period, brain area and task were combined across datasets. The Wilcoxon signed rank test (p<0.005) was used to assess if coherence accuracy differences were considered significantly larger than 0. The same criterion was used to assess significance of differences in coherence effects magnitude between brain areas within the same time period.

### Latency analysis

We determined the latency for choice predictive signals during the dots period as the first of three consecutive (and non-overlapping) 50 ms time steps for which the prediction accuracy is significantly larger than chance (0.5) according to a Wilcoxon signed rank test, p<0.001.

### Decision Variable

When performing logistic regression on the population activity, the set of weights associated with each neuron form the hyperplane that best separates leftward and rightward choices for the corresponding time window (50 ms width at a time). For each trial and time point, the distance of the population state to this hyperplane is given by the model choice log odds, i.e. it corresponds to the model’s certainty about the upcoming choice of the monkey—the further from the hyperplane, and thus the larger the distance, the higher the confidence of the model on its estimate of the eventual choice of the animal (equation 4):

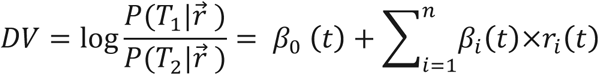

Where *r_i_(t)* are the z-scored summed spike counts for neuron *i* and time window *t*, *β_0_*(t) is an intercept term and *β_i_(t)* are the classifier weights (one for each neuron and time window). We use this distance as a proxy for an internal decision variable (DV) and study its dynamics as a function of stimulus difficulty and trial epoch. Using longer time windows of neural activity as predictors of choice increased the accuracy even further (Supp. Fig. 5a) at the expense of time resolution.

### Slope Analysis

To analyze the dependency of our putative decision variable on the stimulus strength we fit the single trial DV traces with a tri-linear curve. Data in the interval [0-500] ms aligned to dots onset was used to fit the curves. For the variable duration task no data after the go-cue was presented was used in this fit. We fix the first slope at zero since the stimulus does not influence the neural representation of choice within the first ∼100-150 ms following stimulus onset. The intercept, the 2nd and 3rd slopes, as well as the transition times are all free parameters. All free parameters were fit to minimize squared error. We used the value of the 2nd slope to quantify the DV initial rate of rise due to motion information. Since the subsequent analyses focused on this parameter, variable duration trials with stimulus duration in the [200,500] ms range were also used.

The curves were fitted independently for each trial and the fitting procedure was blind to stimulus coherence or task variant. Only correct trials were used in this analysis. Within each task we then tested if the slopes resulting from the fitting procedure had a statistically significant dependence on coherence. We did this in two ways: 1) by regressing slopes as a function of stimulus coherence and 2) by regressing slopes as a function of log2(stimulus coherence). The results were similar in both cases.

Finally we tested the effect of coherence, task variant, and their interaction on slopes across tasks for each brain area and target direction by fitting the following model:

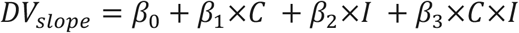

Where *DV_slope_* is the 2^nd^ slope, *C* is the magnitude of the stimulus coherence, normalized to 1 and *I* is the task identity (0 for fixed duration, 1 for variable duration).

The resulting *β*_0_ is the intercept term, *β*_1_ quantifies the effect of coherence on slopes (across tasks), *β*_2_ quantifies the shift in slope magnitude between tasks (across coherences) and *β*_3_ captures the coherence dependency of the offset of the slopes between tasks. A significant and positive/negative *β*_1_ value would indicate slopes increase/decrease as a function of coherence, a significant and positive *β*_2_ value would imply slopes are higher for the variable duration task compared to the fixed duration task (across coherences), and a significant and positive *β*_3_ value would imply the offset between variable duration and fixed duration slopes is coherence dependent.

All slope analyses were done on correct trials only to assure coherence effects were not a result of including higher number of incorrect trials for low coherences.

### Behavioral metrics prediction

We attempted to predict/explain four behavioral metrics based on neural activity throughout the trial: hand reaction time, eye reaction time, hand peak velocity and eye peak velocity.

To predict hand or eye Reaction Time (RT) based on neural activity we performed Ridge regression on the z-scored firing rates of all units within a 150 ms window according to:

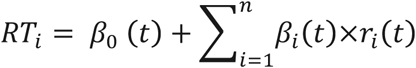

Where *RT_i_* is the behavioral reaction time on a given trial, *n* is the number of units, *r_i_(t)* is response of unit *i* at time *t* and the *β* coefficients are the fit model parameters. For each window, a different model was trained for each reach direction (left and right) on 90% of the correct trials that lead to the corresponding choice. Then the RTs on the remaining 10% of the trials were estimated using the trained model and the units firing rates. We performed this same process 10 times for each window (10-fold cross validation) and obtained a set of estimated Reaction Times. We then performed a linear regression between the estimated and the observed reaction times for all trials and recorded the R-squared value. Finally, we slid the window by 20 ms and repeated the process until all relevant epochs of the trial were tested. The adequate Ridge parameter was estimated independently for each dataset and reach direction for the window comprising [200, 350ms] after the Go cue, where the RT signal tended to be strongest. The estimation was performed using 10-fold cross validation over 20 potential values. The value corresponding to the smallest testing error was chosen and used to regularize the linear model in every window. The exact same procedure was followed when attempting to predict hand and eye peak velocity. Trials with hand RT lower than 150 ms or higher than 800 ms were excluded from this analysis.

### Integration models

We investigated which variations of the integration-to-bound models could potentially explain changes in the dynamics of single trial DVs in the fixed and variable duration tasks. To contrast quantitative predictions of these models, we implemented a basic integration model and its three key variants for acceleration of choice representation by changing input gain, urgency, and recruitment of choice-representing units. The basic model assumes that two large neural populations integrate sensory evidence in favor of the two competing choices. Each integrator receives two types of inputs. The first is the momentary evidence about motion:

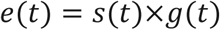

where *s*(*t*) is a random draw from a Gaussian distribution and *g*(*t*) is a gain term that scales the input of the integrator. The mean of the Gaussian distribution of *s*(*t*) is *k*×*C* for one integrator and −*k*×*C* for the second integrator, where *C* is the signed motion coherence in a trial, and *k* is the sensitivity coefficient for motion on the display monitor. The linear dependence of the mean of momentary evidence on motion strength is compatible with neural responses in motion selective areas MT and MST (Britten et al., 1996, Celebrini and Newsome, 1994). The standard deviation of the Gaussian distribution is 1. The second input to the integrators is an urgency signal that drives both integrators toward their bounds. The accumulated evidence is:

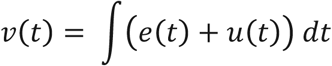

Each integrator has a lower reflective bound *B_l_* and an upper absorbing bound *B_u_* (Kiani et al., 2014a). The integration continues until one of the integrators reaches the upper bound. At that time, a choice is made and the two integrators maintain their states until the end of the motion presentation period. The model shows a monotonic improvement of choice accuracy with motion strength, consistent with the monkey’s behavior.

To mimic our recordings, we simulated 100 spiking units from each of the two integrator populations. The spikes were generated based on an inhomogeneous Poisson process. The instantaneous firing rate of each unit at each moment in a trial was a weighted average of the accumulated evidence and a choice-representing signal:

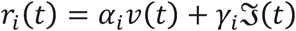

where *r_i_*(*t*) is the firing rate of unit *i* at time *t*. *α_i_* and *γ_i_* are weights between 0 and 1 and determine the tuning of the neuron for representing integrated evidence and choice. The choice representing signal, *ℑ*(*t*), is assumed to be a monotonic function created by a non-linear transformation of *v*(*t*). We chose an accelerating function based on distance of accumulated evidence from the decision bound (*B_u_*)

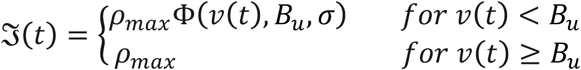

where Φ(.) is a cumulative Gaussian function, *ρ_max_* sets the maximum of *ℑ*(*t*), and *σ* determines the rate of acceleration. Introducing more realism by allowing response correlations, similar to those observed in electrophysiological recordings, or testing other monotonic functions did not significantly change our conclusions about the models.

We simulated 3000 trials for each motion direction and coherence, saved the spike times of the units, and used a logistic regression to calculate the single trial DVs, just as we did for the PMd and M1 neural responses. For the base model and its gain and urgency variants, *γ_i_* were set to 0, making the units represent only the accumulation of evidence. For the progressive recruitment model, *γ_i_* could take any value between 0 and 1, making the neurons represent a mixture of accumulated evidence and categorical choice signal. *α_i_* and *γ_i_* where chosen independently. For the simulations presented in this paper, the parameters of the base model were *k* = 0.3, *g*(*t*) = 1, *u*(*t*) = 0, *B_l_* = −5 and *B_u_* = 20. The same parameters were used for the progressive recruitment model, while *ρ_max_* was set to 40. For the model with high input gain, *g*(*t*) linearly increased from 1 to 3 over 1000ms. For the model with low gain, *g*(*t*) grew from 1 to 1.25. For both models *u*(*t*) was 0. For the models with low and high urgency, *u*(*t*) was 0.005 ms^−1^ and 0.025 ms^−1^, respectively, and *g*(*t*) = 1. Our conclusions are not critically dependent on these specific numbers and hold for a wide range of model parameters, as long as the upper absorbing bound is low enough to curtail the integration process, compatible with the monkeys’ behavior (Fig. 1). Simulations of the integration process and spiking of the model units were done with 1ms time steps.

### Stability of the population choice vector

For each dataset, we divided the trials in two disjoint sets. Within each set of trials, we then modeled the (z-scored) firing rate of each unit at each point as a linear combination of task related predictors (equation 9):

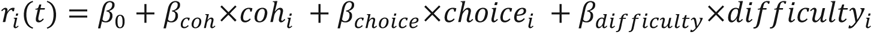

Where *β*_0_ is the intercept term, *coh_i_* is the signed stimulus coherence on trial *i.* (1 for 51.2% rightward motion and -1 for 51.2% leftward motion), *choice_i_* is the behavioral choice on trial *i* (1 right choice and -1 for left choice) and *difficulty_i_* is the absolute stimulus coherence on trial *i* (1 for 51.2% trials and 0 for 0% trials)

At every 10 ms interval, we obtained a *β*_choice_ coefficient for each unit. We concatenated these values for all units into a vector 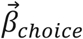 and normalized it. We repeated this same procedure across all time points and obtained a matrix *B*_1,*choice*_. We then followed the exact same steps for the second set of trials and obtained a second matrix *B*_2,*choice*_. Next, we projected *B*_1,*choice*_ onto *B*_2,*choice*_ to obtain a cross-validated measure of vector alignment (dot product) matrix. Because *B*_1,*choice*_ and *B*_2,*choice*_ were calculated using disjoint sets of trials the diagonal values are meaningful and not set to 1 by convention (Fig. 6, Supp. Figs 16 and 17). To further average out spurious vector alignment when the choice signal is small, for each dataset we performed the dot product matrix calculation 20 times always starting from different sets of trials. We then averaged the dot product matrices across iterations and finally across datasets within each condition (brain area, task - Fig. 6a-d, Supp. Figs 16 and 17). For the comparison across tasks we used data from 3 sessions in monkey F for which we collected data on the fixed duration task and the variable duration task back to back (Fig. 6e-f). In total, these sessions comprised 1425 trials in the fixed duration task and 2466 trials in the variable duration task. For these sessions, neural activity was hand sorted and only channels with waveform shapes deemed stable and easily identifiable in both blocks of trials were included in the analyses. The sorting procedure was done prior and without knowledge of the results of the choice stability analyses. The median average of channels excluded this analysis was only 8 out of up to 96 channels.

The analysis of the stability of the population choice vector could have been implemented using the discriminant hyperplanes obtained from the logistic regression analysis. However, we instead performed the linear regression described above for two reasons. First, the beta values for the discriminant hyperplane are obtained using aggressive L1 regularization, which pushes the lowest weights to zero to improve prediction accuracy and avoid overfitting. The logistic classifier could underestimate the contribution of neurons with small but significant choice representation especially late in the trial when strong choice selectivity arises in the population. This scenario would lead to “artificially” lower dot product of the choice axes across time. Instead we used three regressors in our linear model for choice, signed motion strength and absolute motion coherence. The choice regressor will capture the choice representation while the signed motion regressor will capture motion related signals that are not fully explained by choice. Finally, we included a stimulus difficulty regressor that captures non-directional motion coherence signals whose presence has been reported in some LIP cells (Meister et al., 2013).

### Choice predictive units

We applied the same linear model described above (equation 9) to model the (z-scored) firing rate of each unit at each time point and across all trials. For each time point we extracted a *β_choice_* and an associated p-value. We considered a unit to be significantly modulated by choice if *β_choice_* was significantly different from zero at five consecutive 10ms time points (p<0.05, Holm-Bonferroni corrected across all time points). The first of those data points was considered to be the onset of significant modulation for choice for that particular unit. We extended this analysis across all units within a dataset and calculated for each time point the cumulative fraction of units with significant choice modulation. The results were then averaged across datasets within the same condition (brain area, task).

To quantify how reliably each neuron predicted choice over time, we calculated the auROC (Shadlen and Newsome, 2001) metric for every 50 ms of the periods analyzed. According to our convention, right preferring neurons had 1 > auROC > 0.5 and left preferring neurons 0 < auROC < 0.5 (Supp. Fig.24). To collapse across both choice directions, we calculated | auROC -0.5| (Fig. 7c-f). The units analyzed were collected in a session with both a fixed and variable duration block (monkey F) from channels whose waveforms were deemed stable (see above). For this representative session (Fig. 7c-f) only data from one channel in PMd and eight channels in M1 (out of up to 96) were excluded.

### Unit dropping

For the unit dropping analysis we fit a logistic model (equation 3) to data obtained in the last 50 ms of dots presentation using 10-fold cross validation, just as before. The lambda regularization parameter however, was in this case fit to the same 50 ms epoch we would test (again using 25 potential values and 10-fold cross validation). The set of beta coefficients of the model corresponding to the lowest deviance lambda parameter was then chosen and ranked by magnitude. We removed from the data the unit with highest beta coefficient and re-trained and re-tested the model using 10-fold cross-validation and recorded the accuracy. This process was repeated 70 times until the 70 units with highest beta coefficients (ranked using the full model) were all dropped in descending order.

