## Supplementary Materials for "Population dynamics of choice representation in dorsal premotor and primary motor cortex"

### Supplementary Information

| 2 Monkeys | | PMD | | M1 | |
|---|---|---|---|---|---|
| | | T1 | T2 | T1 | T2 |
| Fixed Duration linear (coh) | p-value | 2.60E-19* | 8.30E-32* | 1.12E-05* | 3.56E-08* |
| | β | 17.11 | -23.44 | 7.46 | -8.93 |
| Fixed Duration $log_2$(coh) | p-value | 3.61E-14* | 1.08E-27* | 4.43E-06* | 1.09E-05* |
| | β | 1.94 | -3.17 | 1.02 | -1.06 |
| Variable Duration linear (coh) | p-value | 1.68E-14* | 3.32E-12* | 8.01E-07* | 6.56E-03 |
| | β | 19.153 | -21.78 | 15.23 | -6.95 |
| Variable Duration $log_2$(coh) | p-value | 3.02E-13* | 1.93E-10* | 1.01E-04* | 8.70E-03 |
| | β | 2.66 | -2.62 | 1.78 | -0.94 |

**Supp. Table 1 – Model fitting of single trial DV slopes as a function of coherence.** To assess the effect of stimulus coherence on the single trial DV slopes we fit two different linear models to the data. The first model tested the whether single trial DV slopes variance could be explained by a linear term on stimulus coherence and the second model tested single trial DV slopes variance could be explained by a linear term on $log_2$ of stimulus coherence. For each model the fits were performed separately for each brain area, choice and task variant. P-values denote the likelihood of wrongly rejecting the null hypothesis under which the linear terms (on coherence and $log_2$ (coherence)) are zero. All fits for a given model, task and area are significant at p=0.05 Bonferroni corrected for 8 comparisons (*). Data from both monkeys, correct trials only.

| Monkey H | | PMD | | M1 | |
|---|---|---|---|---|---|
| | | T1 | T2 | T1 | T2 |
| Fixed Duration linear (coh) | p-value | 3.11E-09* | 2.61E-17* | 1.54E-01 | 4.68E-03* |
| | β | 16.20 | -25.16 | 3.83 | -7.57 |
| Fixed Duration $log_2$(coh) | p-value | 1.82E-06* | 1.48E-14* | 5.91E-02 | 3.41E-02 |
| | β | 1.78 | -3.17 | 0.68 | -0.79 |
| Variable Duration linear (coh) | p-value | 1.78E-03* | 1.11E-05* | 4.06E-05* | 5.62E-02 |
| | β | 11.42 | -19.48 | 15.73 | -7.74 |
| Variable Duration $log_2$(coh) | p-value | 6.06E-04* | 2.25E-04* | 4.00E-04* | 2.65E-01 |
| | β | 1.83 | -2.28 | 1.99 | -0.64 |

**Supp. Table 2 – Model fitting of single trial DV slopes as a function of coherence.** Same as Supp. Table 1 but for fits to Monkey H's data alone.

| Monkey F | | PMD | | M1 | |
|---|---|---|---|---|---|
| | | T1 | T2 | T1 | T2 |
| Fixed Duration linear (coh) | p-value | 2.37E-12* | 9.05E-17* | 9.70E-06* | 5.51E-07* |
| | β | 18.47 | -22.55 | 9.69 | -10.26 |
| Fixed Duration $log_2$(coh) | p-value | 1.91E-09* | 3.46E-15* | 8.88E-06* | 2.95E-05* |
| | β | 2.10 | -3.21 | 1.24 | -1.33 |
| Variable Duration linear (coh) | p-value | 2.25E-14* | 6.52E-09* | 2.44E-03* | 3.59E-03* |
| | β | 25.84 | -25.47 | 14.35 | -8.43 |
| Variable Duration $log_2$(coh) | p-value | 1.67E-11* | 2.34E-09* | 3.16E-02 | 5.03E-05* |
| | β | 3.35 | -3.17 | 1.52 | -1.59 |

**Supp. Table 3 – Model fitting of single trial DV slopes as a function of coherence.** Same as Supp. Table 1 but for fits to Monkey F's data alone.

| 2 Monkeys | | PMD | | M1 | |
|---|---|---|---|---|---|
| | | T1 | T2 | T1 | T2 |
| Intercept | p-value | 7.32E-83* | 2.69E-46* | 3.00E-46* | 2.00E-34* |
| | $\beta_0$ | 9.227 | -8.882 | 6.894 | -6.204 |
| Coherence (C) | p-value | 1.08E-19* | 3.31E-29* | 9.01E-05* | 1.56E-07* |
| | $\beta_1$ | 8.76 | -12 | 3.82 | -4.572 |
| Task Identity (I) | p-value | 1.15E-21* | 2.57E-19* | 8.21E-28* | 6.86E-49* |
| | $\beta_2$ | 8.314 | -9.032 | 9.632 | -12.1 |
| Interaction (C*I) | p-value | 0.5177 | 0.6405 | 0.01526 | 0.4965 |
| | $\beta_3$ | 1.047 | 0.8519 | 3.977 | 1.013 |

**Supp. Table 4 – Model fitting of single trial DV slopes as a function of coherence across both tasks.**
To assess the effect of stimulus coherence, task identity and their interaction on the single trial DV slopes we fit a linear model to the data. The fits were performed separately for each brain area and choice. For each regressor, P-values denote the likelihood of wrongly rejecting the null hypothesis under which the linear term is 0 and (*) denotes significant regressors at p=0.05 Bonferroni corrected for 16. comparisons. Data from both monkeys, correct trials only.

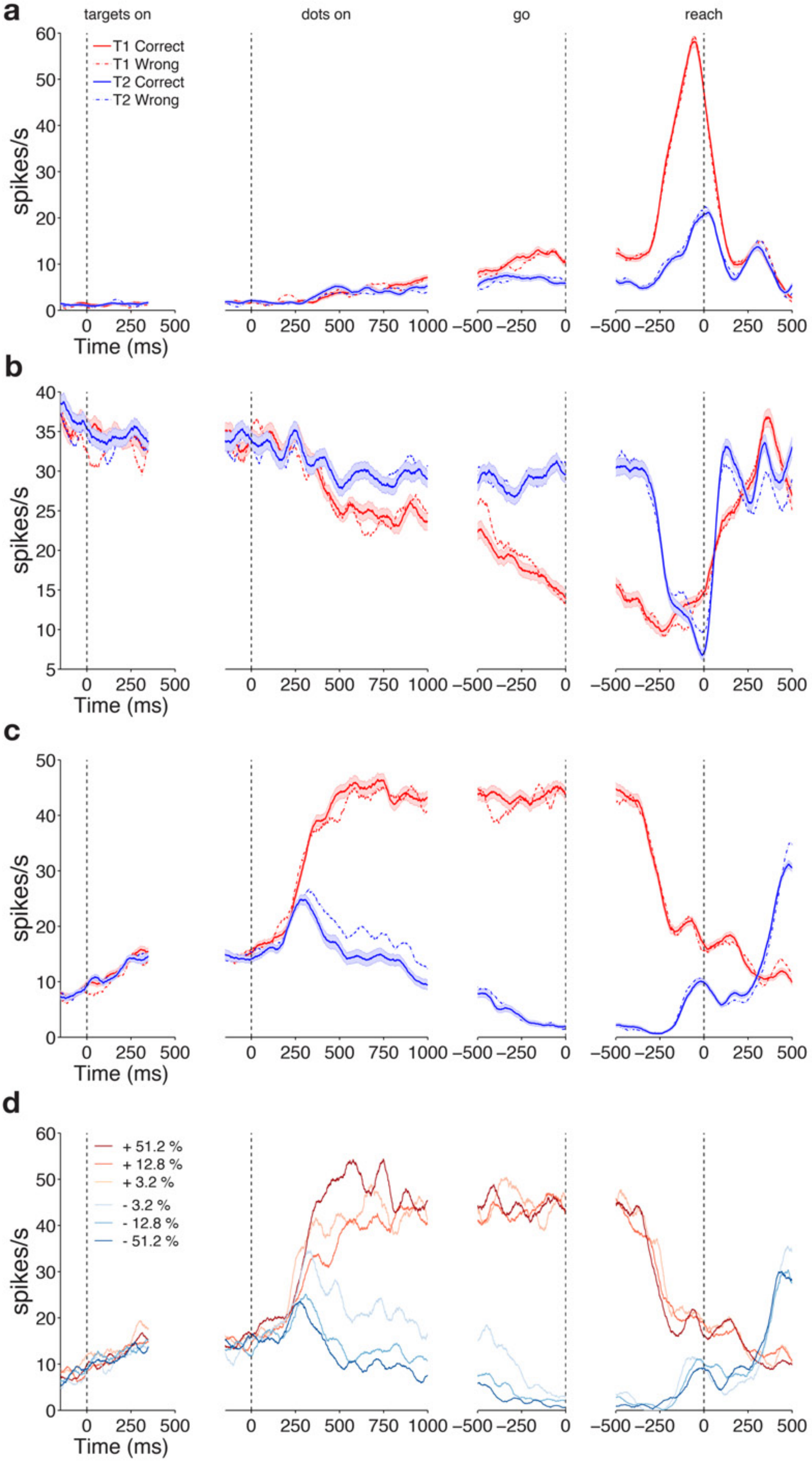

a
targets on
dots on
go
reach
T1 Correct
T1 Wrong
T2 Correct
T2 Wrong
spikes/s
Time (ms)
b
c
d
+ 51.2 %
+ 12.8 %
+ 3.2 %
- 3.2 %
- 12.8 %
- 51.2 %


71

**Supp. Figure 1 – Diverse single unit responses in PMd a)** Neural activity of a well isolated single neuron in PMd. Activity is aligned to four events in the task: targets onset, dots onset, go cue and response. Solid red (blue) lines show average activity level (+/- s.e.m.) for correct right (left) choices. Dashed lines show incorrect choices to the target of corresponding color: red for right and blue for left. **b-c)** Same as **a)** for two other PMd units recorded during the same session. **d)** Same unit as in **c)** but sorting the activity by choice (red for right, blue for left) and stimulus difficulty (dark for easy trials, light for hard trials). Only correct trials were included.

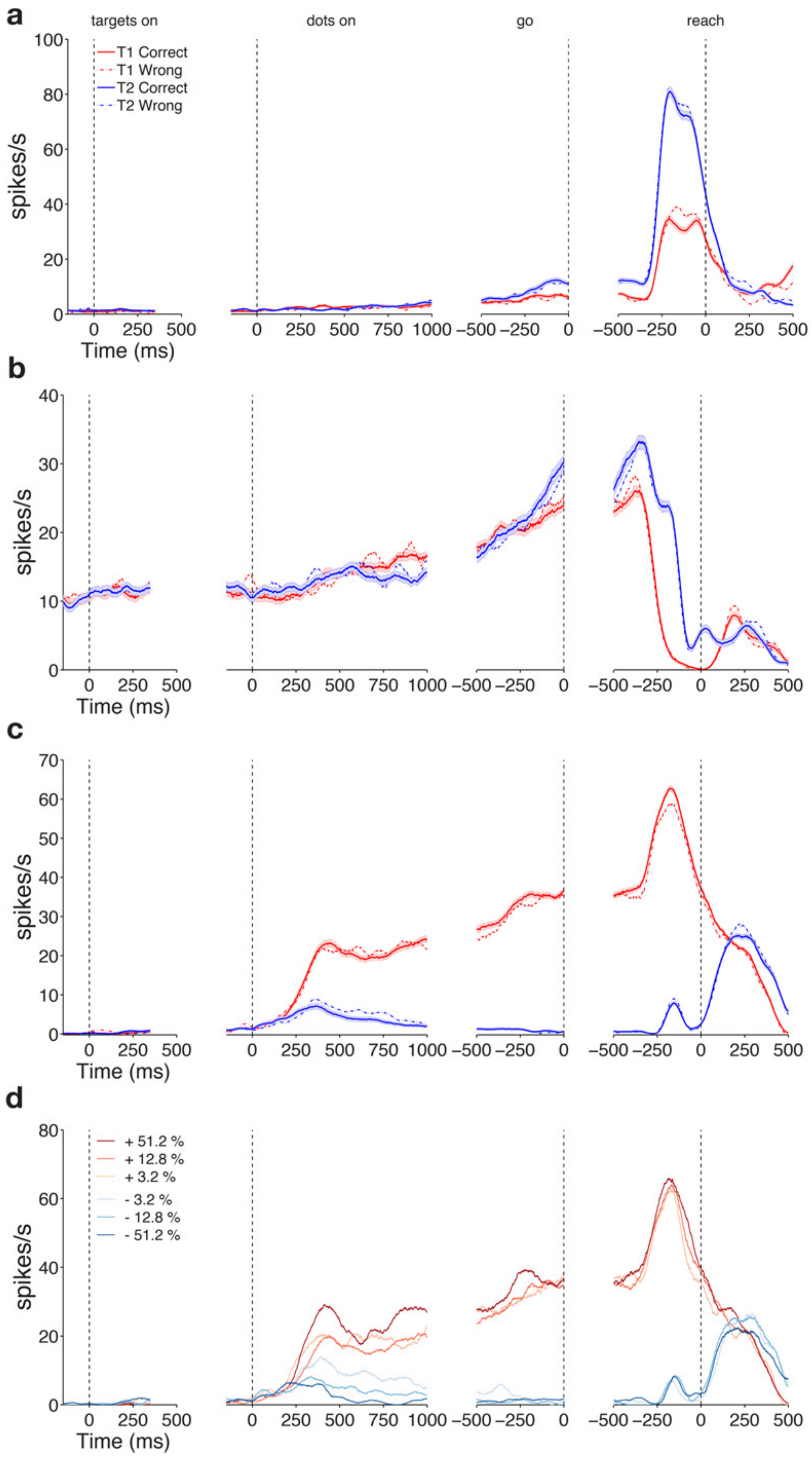
a
targets on
dots on
go
reach
T1 Correct
T1 Wrong
T2 Correct
T2 Wrong
spikes/s
Time (ms)
b
c
d
+ 51.2 %
+ 12.8 %
+ 3.2 %
- 3.2 %
- 12.8 %
- 51.2 %

83

**Supp. Figure 2 – Diverse single unit responses in M1 a)** Neural activity of a well isolated single neuron in M1. Activity is aligned to 4 events in the task: targets onset, dots onset, go cue and response. Solid red (blue) lines show average activity level for +/- s.e.m. for correct right (left) choices. Dashed lines show incorrect choices to the target of corresponding color: red for right and blue for left. **b-c)** Same as **a)** for two other M1 units recorded during the same session. **d)** Same unit as in **c)** but sorting the activity by choice (red for right, blue for left) and stimulus coherence (dark for high coherence trials, light for low coherence trials). Only correct trials were included.

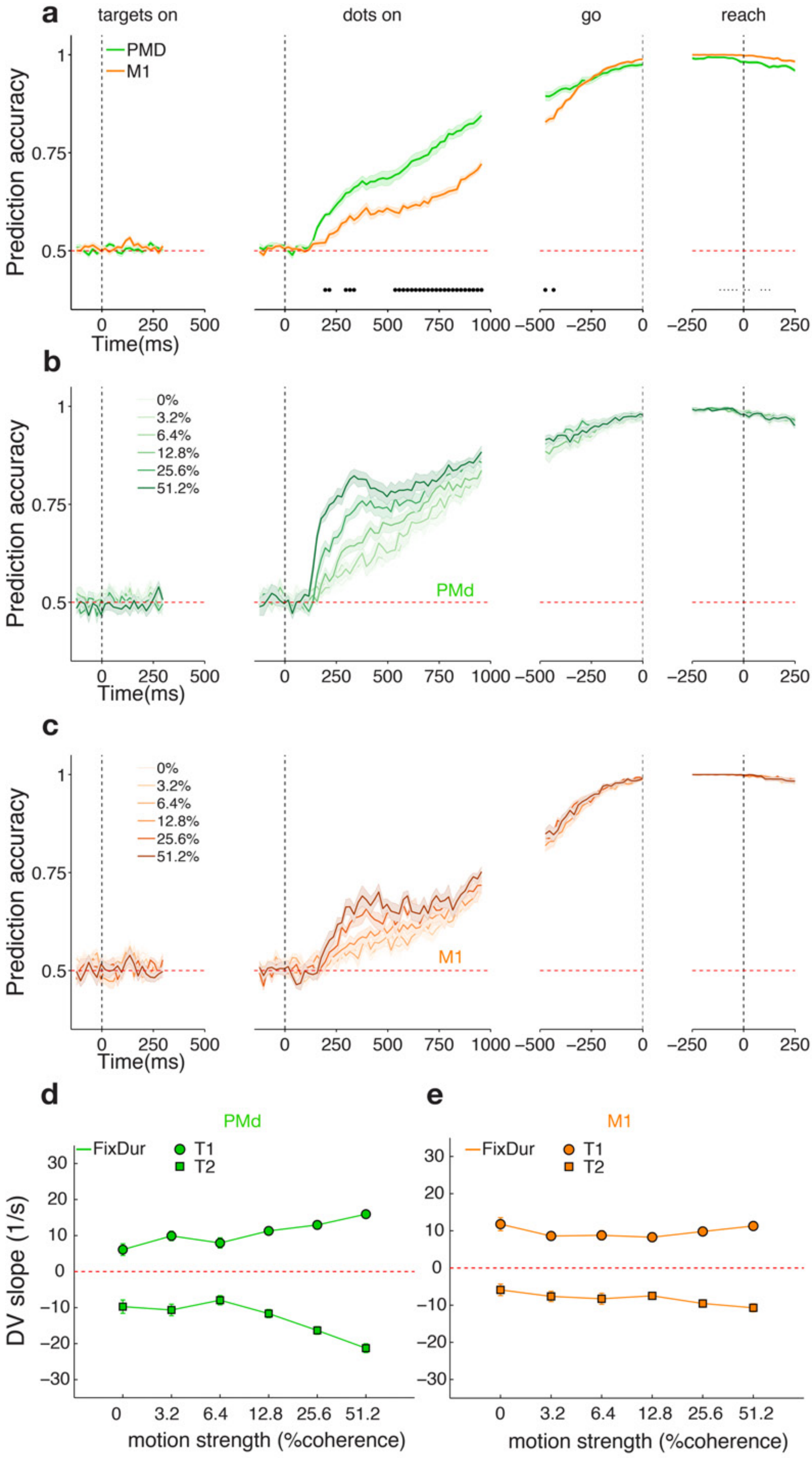
a
targets on
dots on
go
reach
PMD
M1
Prediction accuracy
Time(ms)
b
0%
3.2%
6.4%
12.8%
25.6%
51.2%
PMd
c
M1
d
e
PMd
M1
FixDur
T1
T2
DV slope (1/s)
motion strength (%coherence)

94

**Supp. Figure 3 - Neural population choice prediction accuracy on single trials in the fixed duration task. a-e)** Same format as Figure 2; data for Monkey H only.

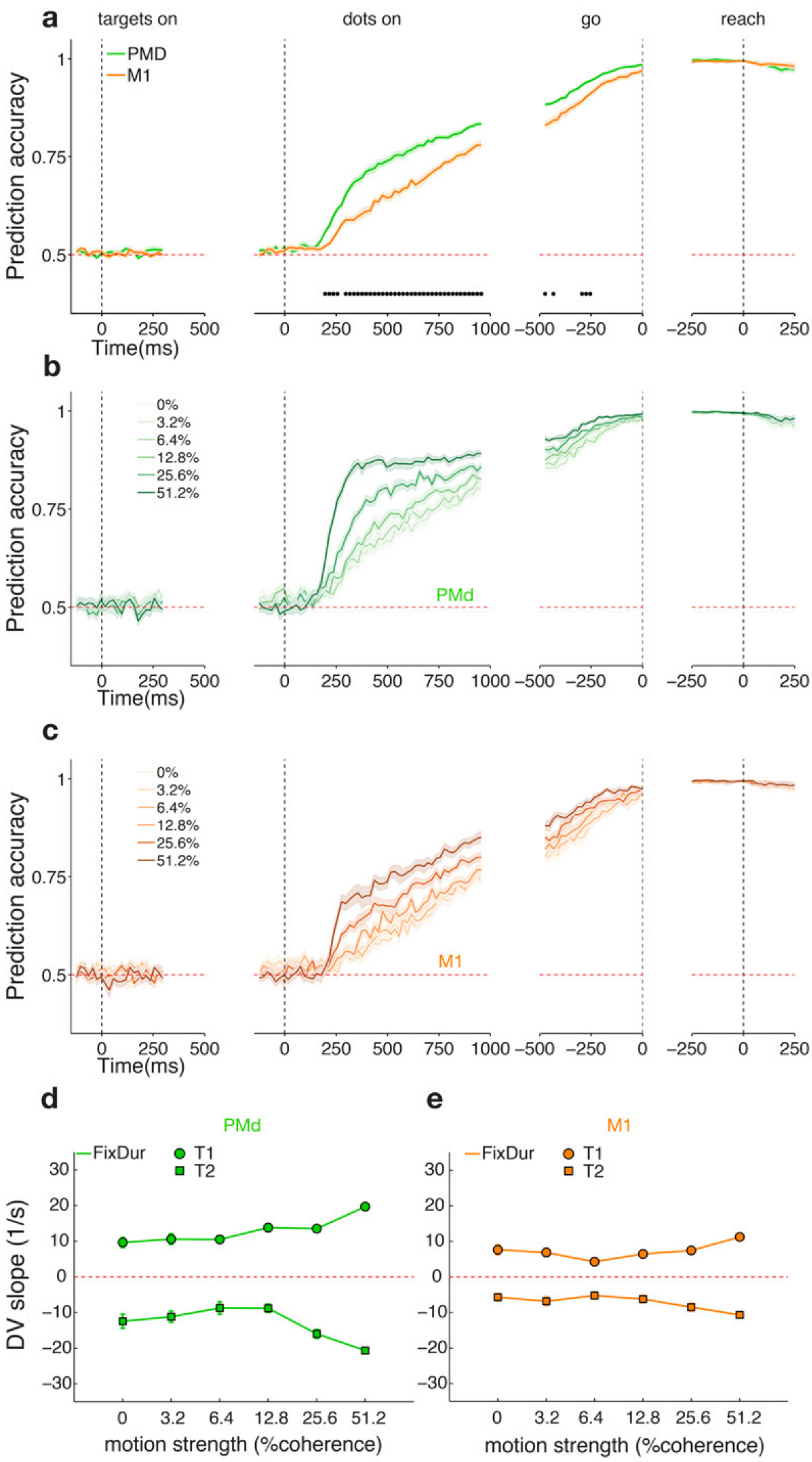

a
targets on
dots on
go
reach
PMD
M1
Prediction accuracy
Time(ms)
b
0%
3.2%
6.4%
12.8%
25.6%
51.2%
PMd
c
M1
d
e
FixDur
T1
T2
DV slope (1/s)
motion strength (%coherence)

**Supp. Figure 4 - Neural population choice prediction accuracy on single trials in the fixed duration task. a-e)** Same format as Figure 2; data for Monkey F only.

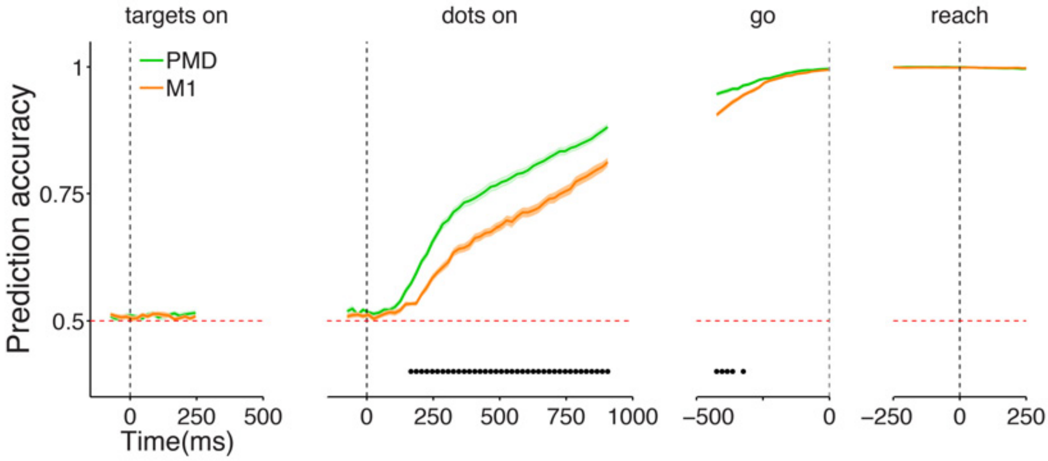


**Supp. Figure 5 - Neural population choice prediction accuracy on single trials (pooled results across 2 monkeys, using 150 ms window).**

**PMd is still more choice predictive than M1 during the stimulus presentation when using a larger window size.** Average prediction accuracy (see Methods) over time +- SEM for monkey H. PMd (M1) data are plotted in green (orange). Black dots denote time bins for which the prediction accuracy was significantly different between the two areas ($p<0.05$ Holm-Bonferroni correction for multiple comparisons). Prediction accuracy does reach higher values in the dots presentation period when using a 150 ms window compared to 50 ms window (88% vs 84% for PMd and 81% vs 76% for M1 at dots offset), demonstrating the values reported in the main text do not correspond to a higher bound on accuracy for linear classifiers on these data.

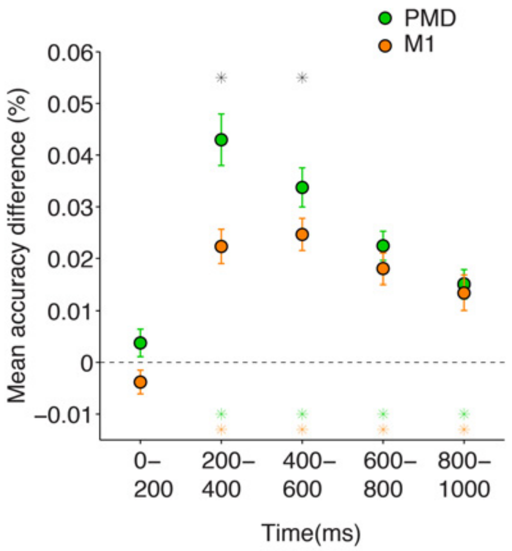


**Supp. Figure 6 – Coherence effects in PMd and M1 in the fixed duration task.**

Coherence effects were defined as the average difference in prediction accuracy between adjacent coherence levels for a given time window in the trial. 5 time windows of 200 ms duration were considered. Data for PMd/M1 is plotted in green/orange. Black asterisks denote windows for which the differences between PMd and M1 were significant (Wilcoxon signed-rank test, P<0.005). Orange/Green asterisks correspond to windows for which the coherence effects were significantly larger than zero (Wilcoxon sign rank test, P<0.005).
Coherence effects are nonexistent in the 200 ms of dots presentation and highest in the 200-400 ms period, after which they slowly decay but remain significantly larger than 0 for the remainder of the stimulus presentation.

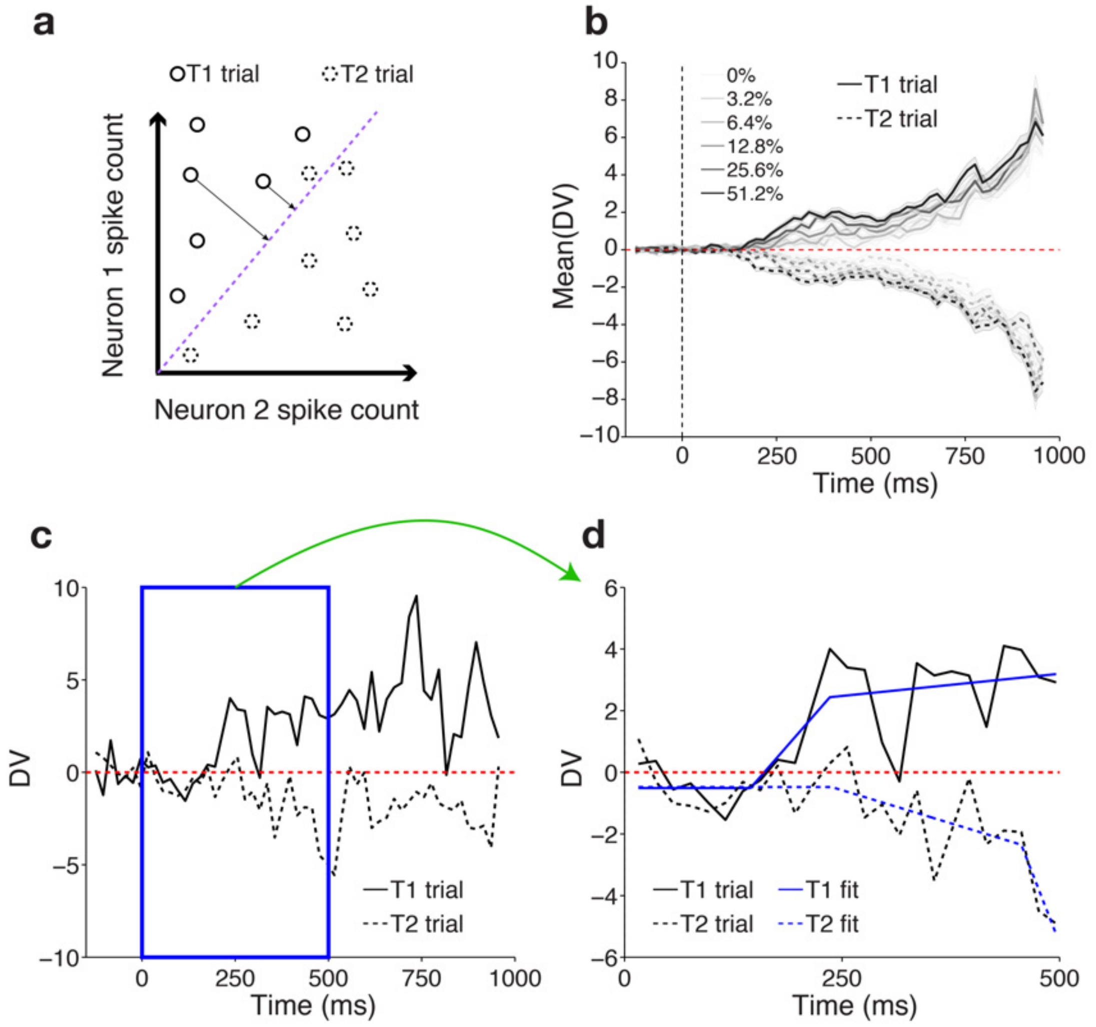


**Supp. Figure 7 – Single trial Decision Variable slope fitting procedure.**

**a) Two-neuron diagram of a linear classifier for choice and putative decision variable.** The spike count of neuron 1 is plotted as a function of the spike count of neuron 2 for a given epoch in the task. The different data points show combinations of neuron 1 and neuron 2 activity for the same epoch across different trials and are labeled based on the ultimate choice of the subject. The purple dashed line depicts the linear classifier boundary that best separates left and right trials based on the neural activity of these two neurons on a given set of training trials. In the illustrated diagram, all right choices (T1 trials) are above the boundary and most left choices (T2 trials) are below the boundary. The T2 trial above the boundary represents a left choice trial that was incorrectly predicted to be a right choice trial. For our logistic regression the confidence of the model in its own predictions can be calculated as the distance of the neural activity to the classifying boundary (length of the black arrows). Even correctly predicted trials will have different degrees of confidence associated with their prediction. This confidence is interpreted as a proxy for a decision variable (DV).

**b) Decision variable increases in magnitude as a function of time and stimulus coherence for both choices.** Average value of the model decision variable during the dots presentation as a function of time, stimulus coherence, and choice. Positive values correspond to higher likelihood of a right choice and negative values to a higher likelihood of a left choice at the end of the trial. Solid traces indicate trials that ended in a rightward choice; dashed traces indicate leftward choices. Darker tones corresponding to high coherence (easy) stimuli; lighter tones to low coherence (harder) stimuli. Only correct trials were analyzed and plotted. The shaded areas indicate +- SEM. Results for one example dataset from Monkey H PMd. As expected from Figure 2b, DV depends on stimulus difficulty in a lawful manner.

**c) Single trial decision variable traces.** Traces of the model decision variable during the dots presentation as a function of time are shown for two example trials. Solid trace indicates a trial that ended in a rightward choice; dashed trace indicates a leftward choice trial. Data from Monkey F PMd in the fixed duration task. To analyze how the initial DV slopes vary with coherence on single trials, we focused on window during which average coherence effects were strongest ([0-500] ms aligned to dots onset, blue box).

**d) Tri-linear fits to truncated single trial DV traces.** Same data as in **c)** truncated to [0,500] ms and aligned to dots onset, fit with tri-linear curves. The results of the fits for the corresponding trials are shown in blue. For our analysis we focused on the initial non-zero slope as the best signature of the rate of DV change following stimulus onset.

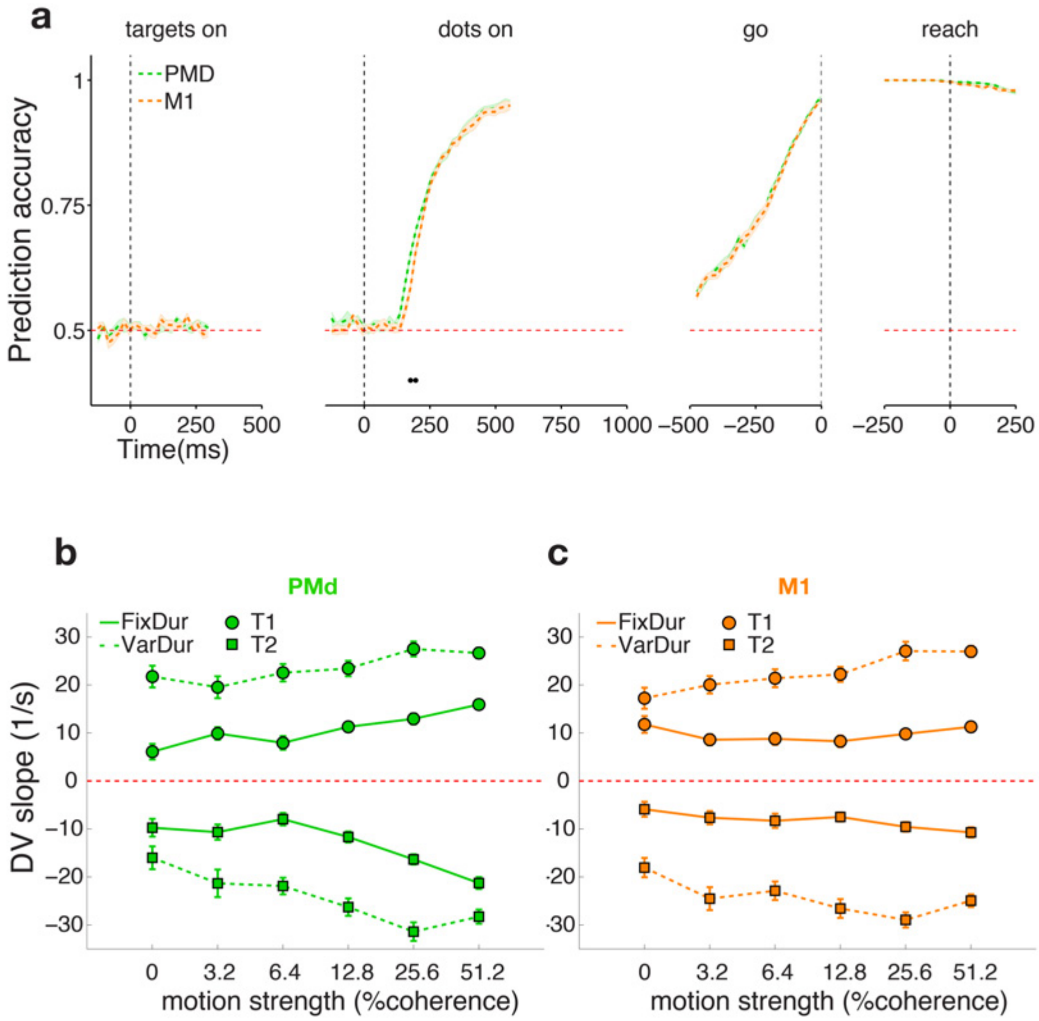


**Supp. Figure 8 - Effects of stimulus duration uncertainty on choice prediction accuracy and single-trial measurements of the model decision variable. a-c)** Same as Figure 3 for Monkey H only.

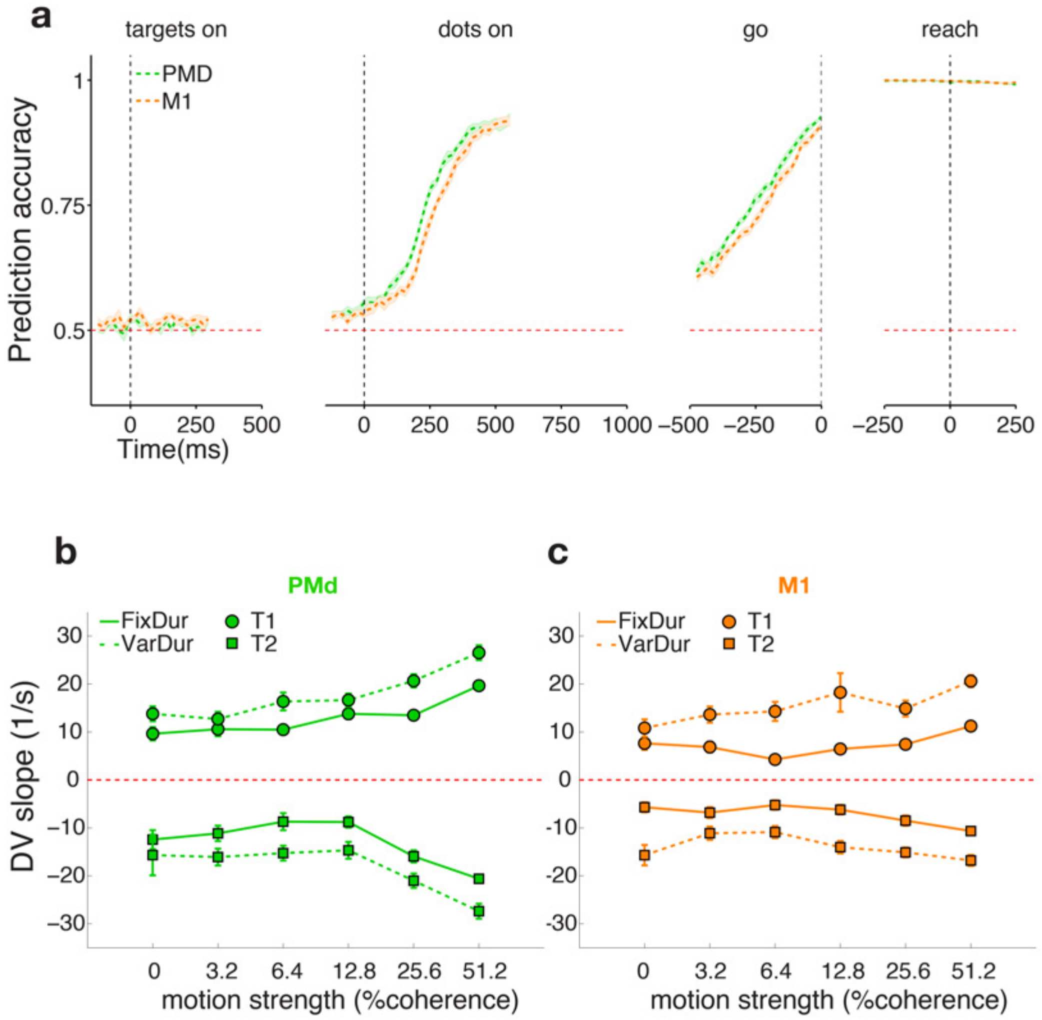


**Supp. Figure 9 - Effects of stimulus duration uncertainty on choice prediction accuracy and single-trial measurements of the model decision variable. a-c)** Same as Figure 3 for Monkey F only.

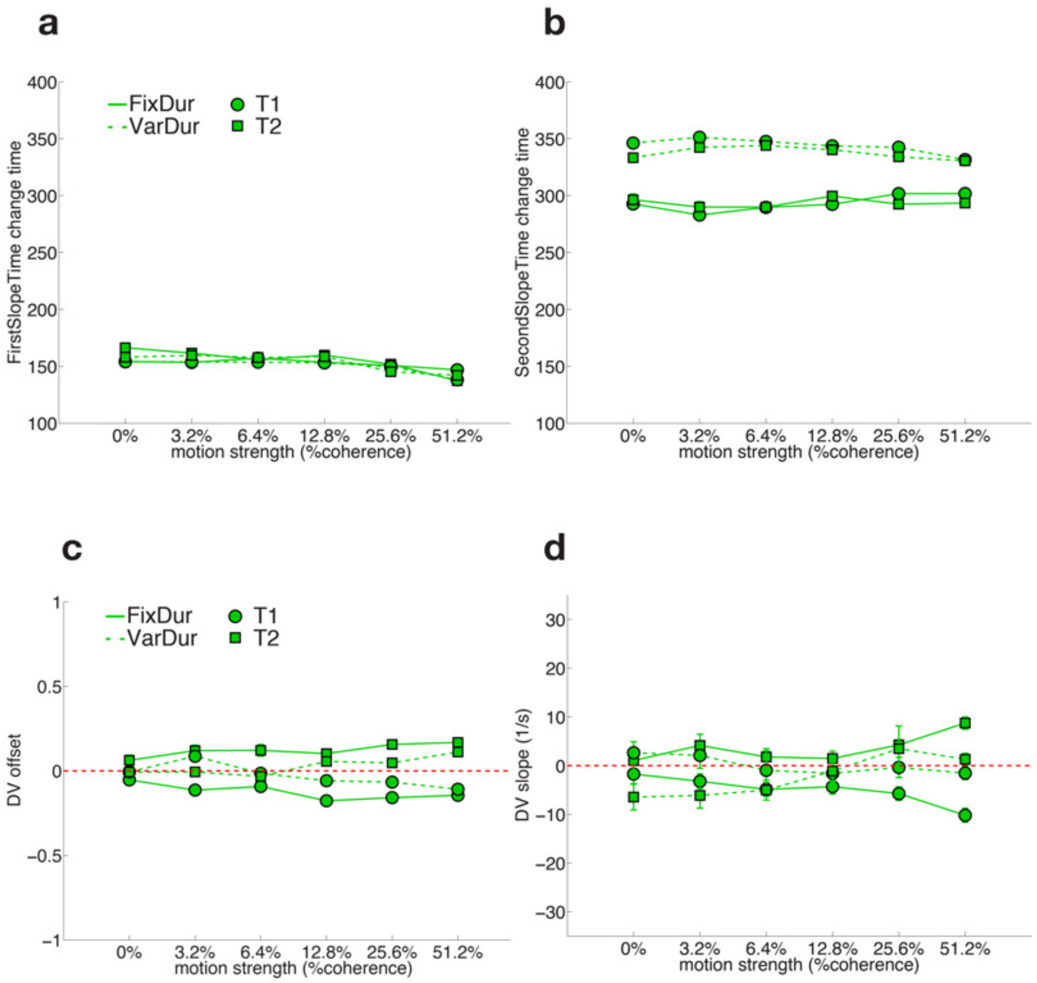


**Supp. Figure 10 – Additional parameters obtained for single trial DV fits in PMd for both targets and tasks. a)** Time of first slope change, **b)** Time of second slope change, **c)** DV Offset (average baseline DV at dots onset) and **d)** Second non-zero slope, as a function of coherence, choice and task

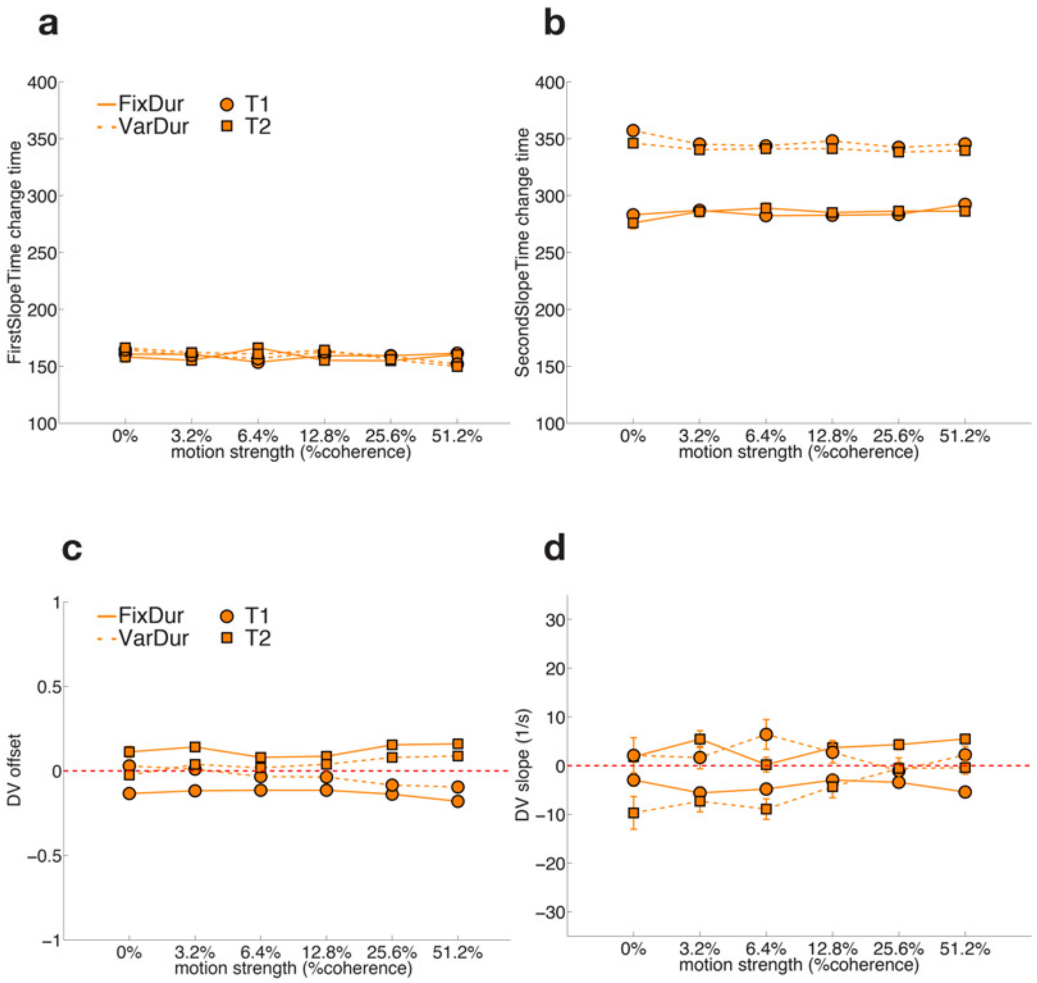


**Supp. Figure 11 – Additional parameters obtained for single trial DV fits in PMd for both targets and tasks. a)** First slope time change, **b)** Second slope time change, **c)** DV offset and **d)** Second non-zero slope, as a function of coherence, choice and task

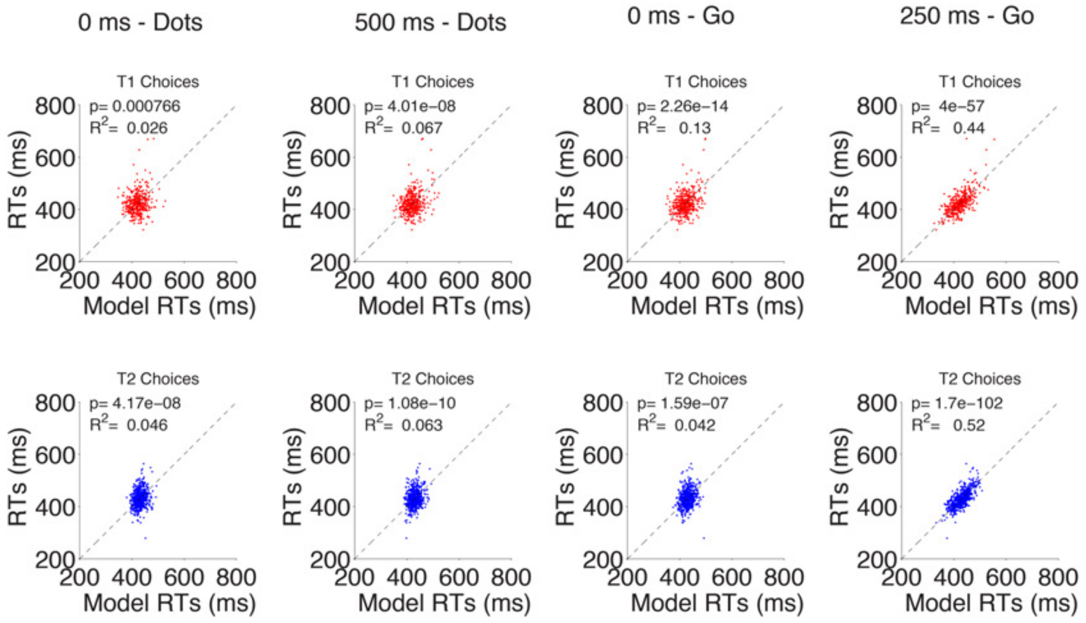

**Supp. Figure 12 – RT prediction model performance on individual time points for a representative session in the fixed duration task.** Scatter plot of measured RTs (y-axis) vs model RTs (x-axis) predicted from neural activity at different time points in the trial. Measured RTs are the same in each panel of a row. Top/bottom panels show results for T1/T2 choices. Each column corresponds to one time point in the trial: dots onset, 500 ms after dots onset, go cue onset, and 250 ms after the go cue onset (from left to right). Insets show $R^2$ and p-value for linear regression between measured and model RTs for each choice and time point. Each dot corresponds to one trial. The closer the dots get to the identity line the higher the model performance. Data for one session from Monkey F M1 in the fixed duration task.

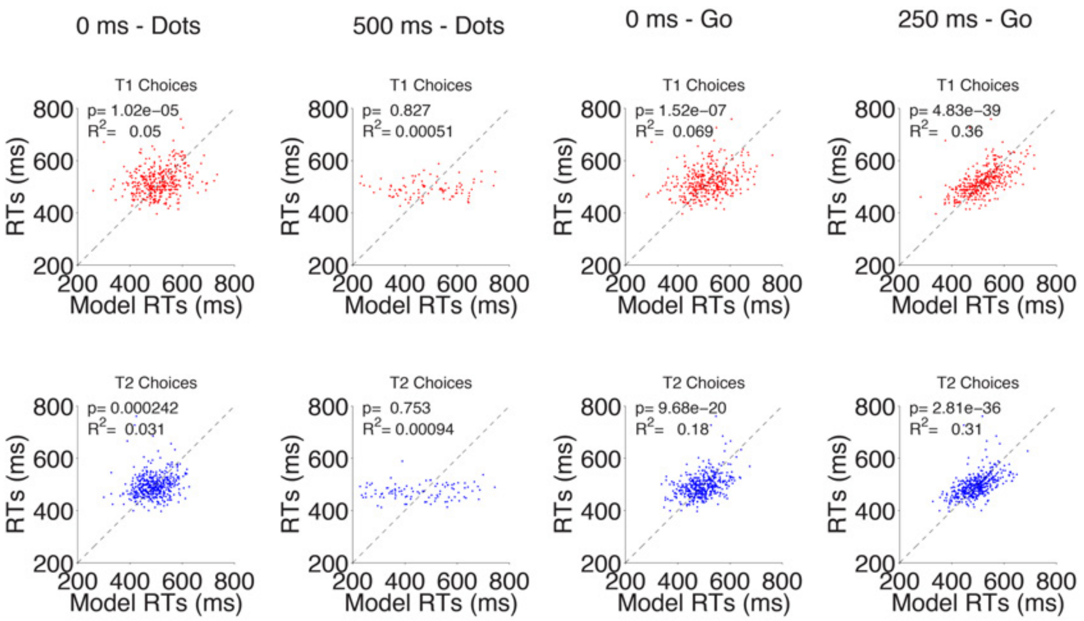


**Supp. Figure 13 – RT prediction model performance on individual time points for a representative session in the variable duration task.** Same as **Supp. Figure 12** but for data for one session from Monkey F M1 in the variable duration task.

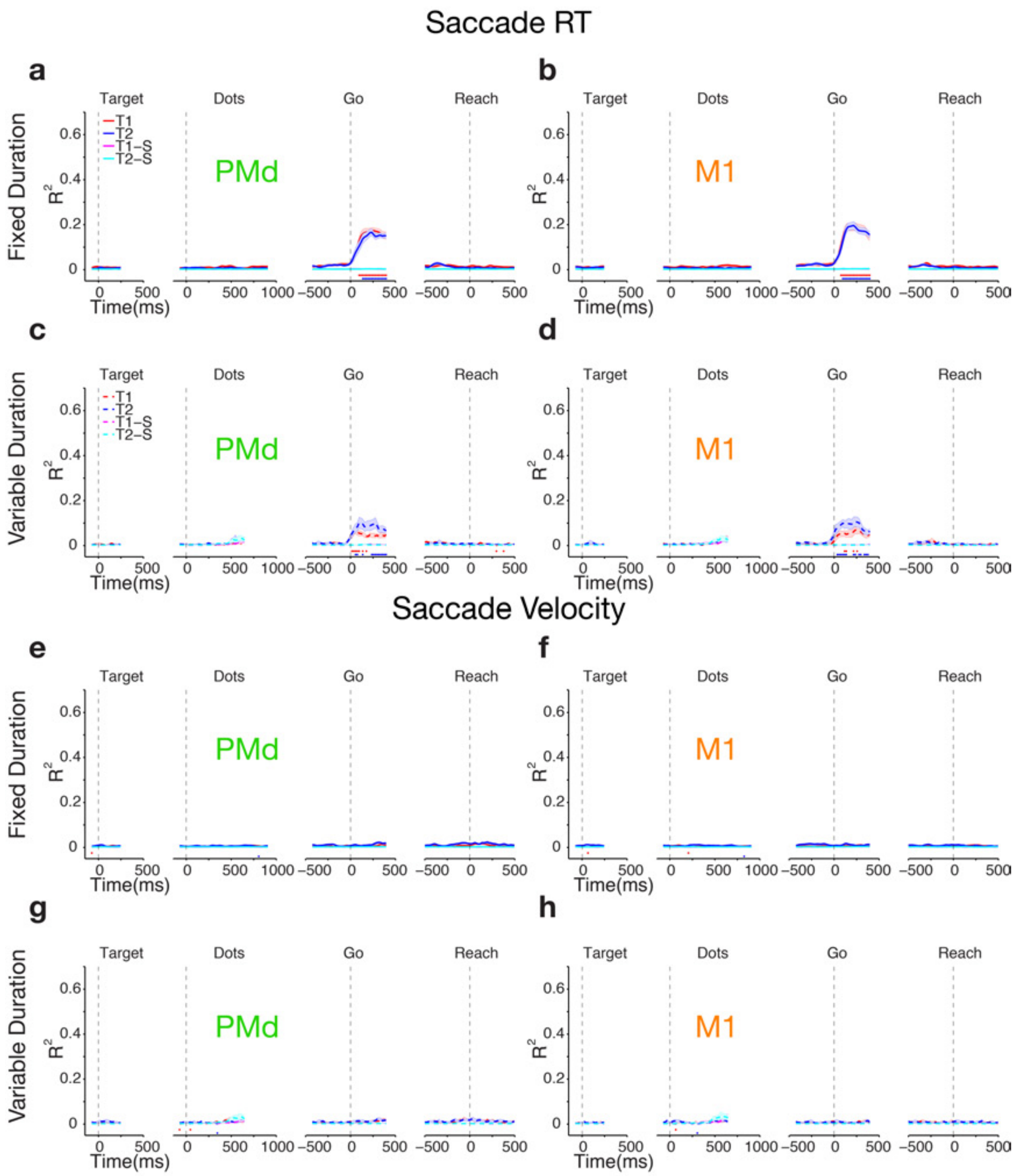


**Supp. Figure 14 - Neural activity in PMd and M1 only becomes predictive of saccade RT around the go-cue in both tasks and is never predictive of saccade peak velocity. a-d)** Same conventions as Figure 4 **a-d)** for saccade RT. **e-h)** Same as Figure 4 **e-h)** for saccade velocity.

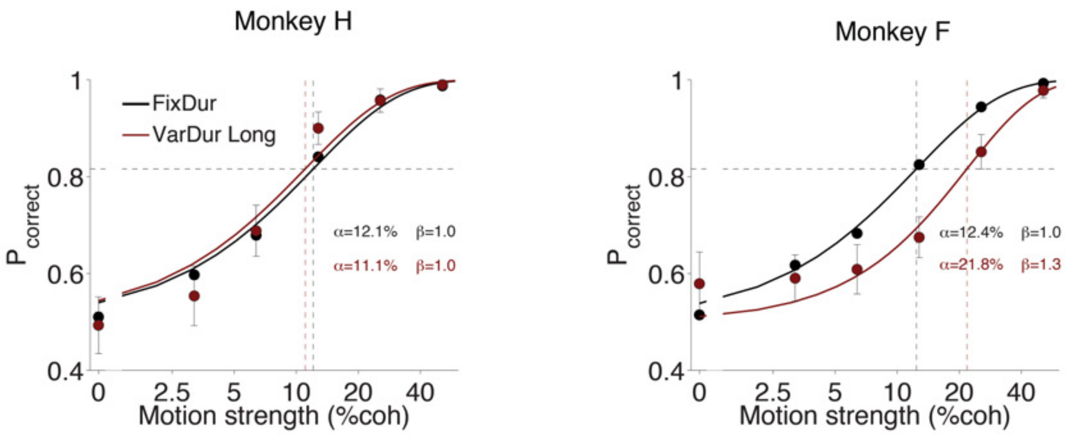


**Supp. Figure 15 - Psychophysical performance in the motion discrimination task for long trials**

Percentage correct is plotted as a function of motion coherence for the fixed duration version (black) for monkey H (left panel) and monkey F (right panel). Data are re-plotted from Fig. 1C. Percentage correct for long (>800 ms duration) trials in the variable duration task is plotted in dark red. Observed data points (+/- SEM) are represented by the dark red and black markers. The data for each task was independently fit with Weibull curves (red and black curves). 17167/ 17440 trials for the fixed duration task and 461/569, >800 ms duration trials for the variable duration task for monkey H/F, respectively. Insets show the fit parameters for the corresponding trials.

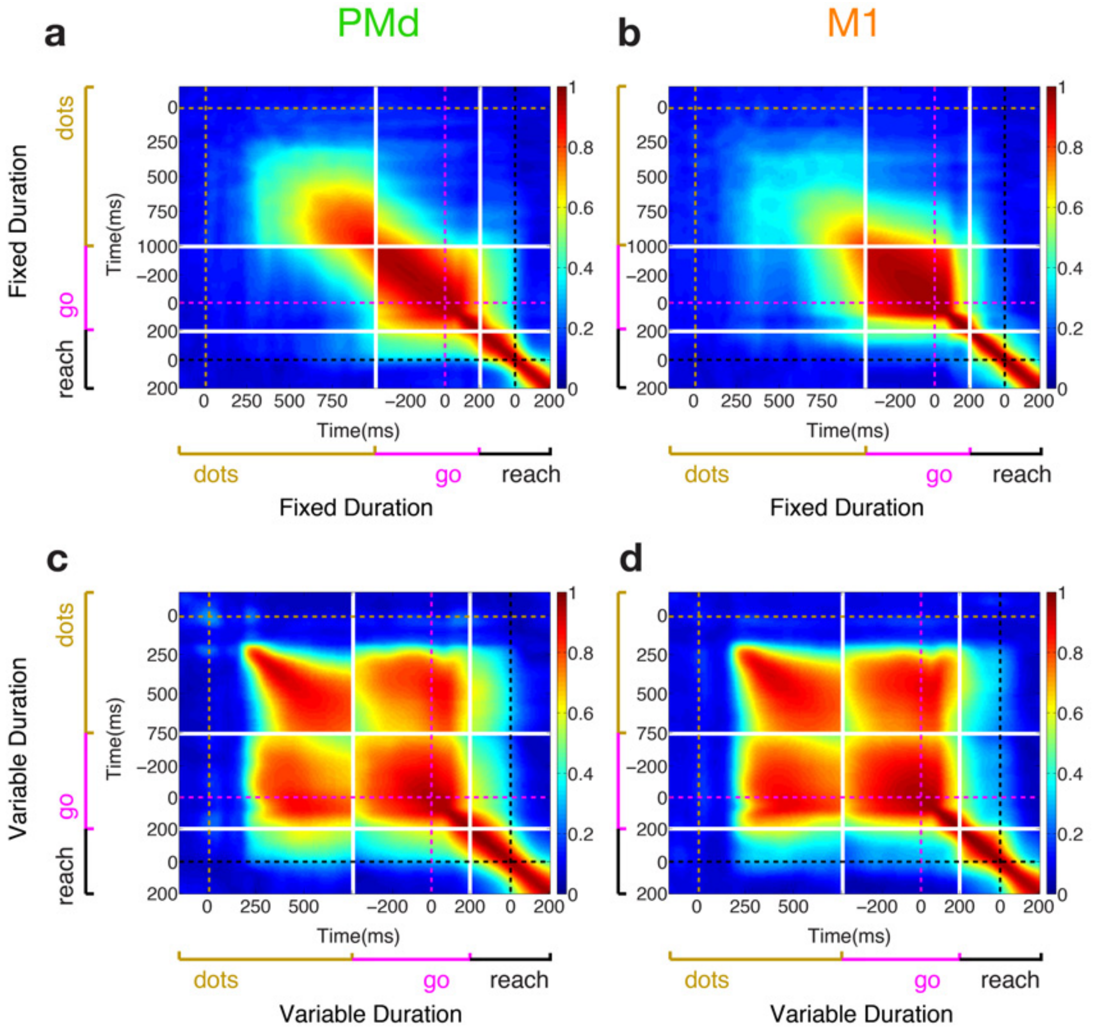


**Supp. Figure 16 – Stability of choice representation during dots is dependent on the statistics of stimulus duration. a-d)** Same as figure 6 **a-d)** for Monkey H.

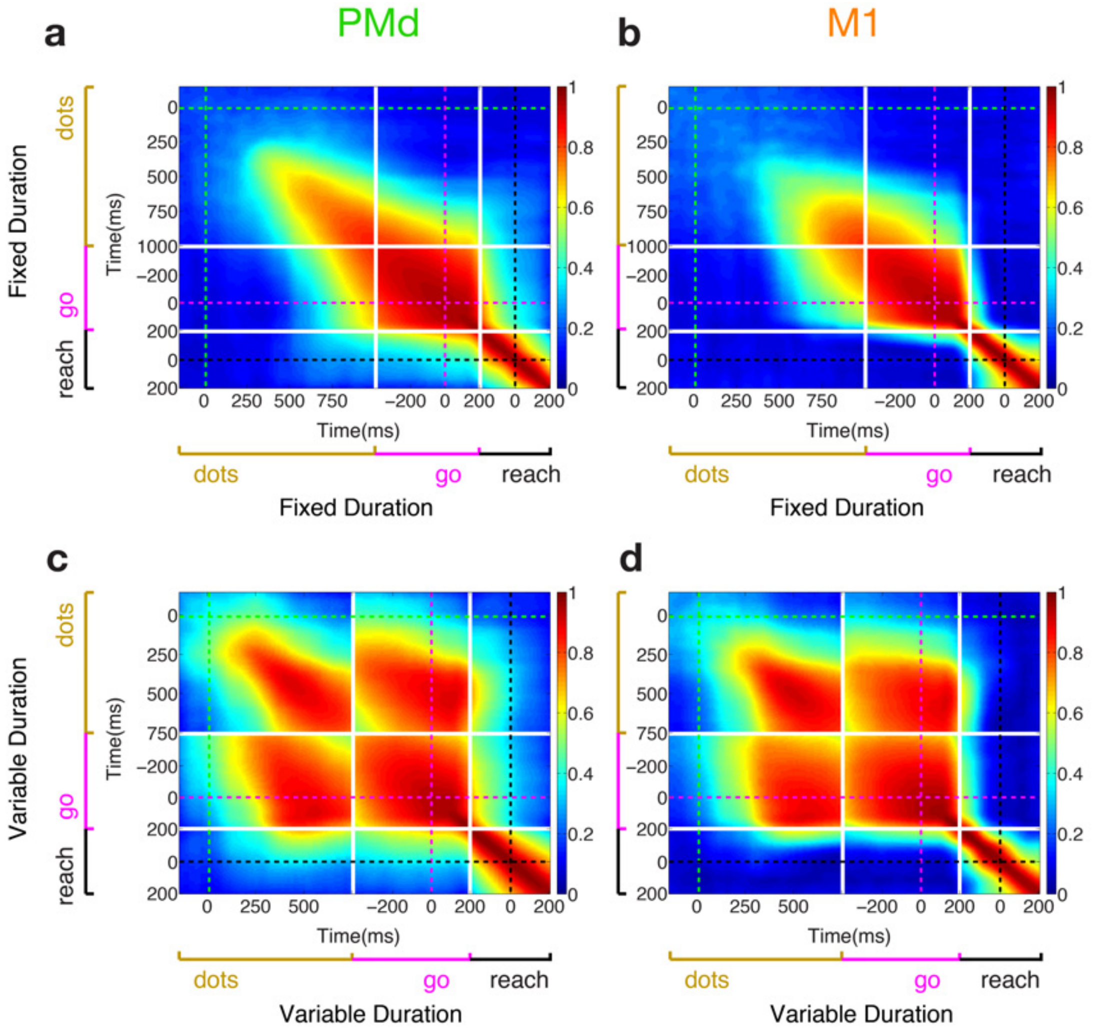


**Supp. Figure 17 – Stability of choice representation during dots is dependent on the statistics of stimulus duration. a-d)** Same as figure 6 **a-d)** for Monkey F.

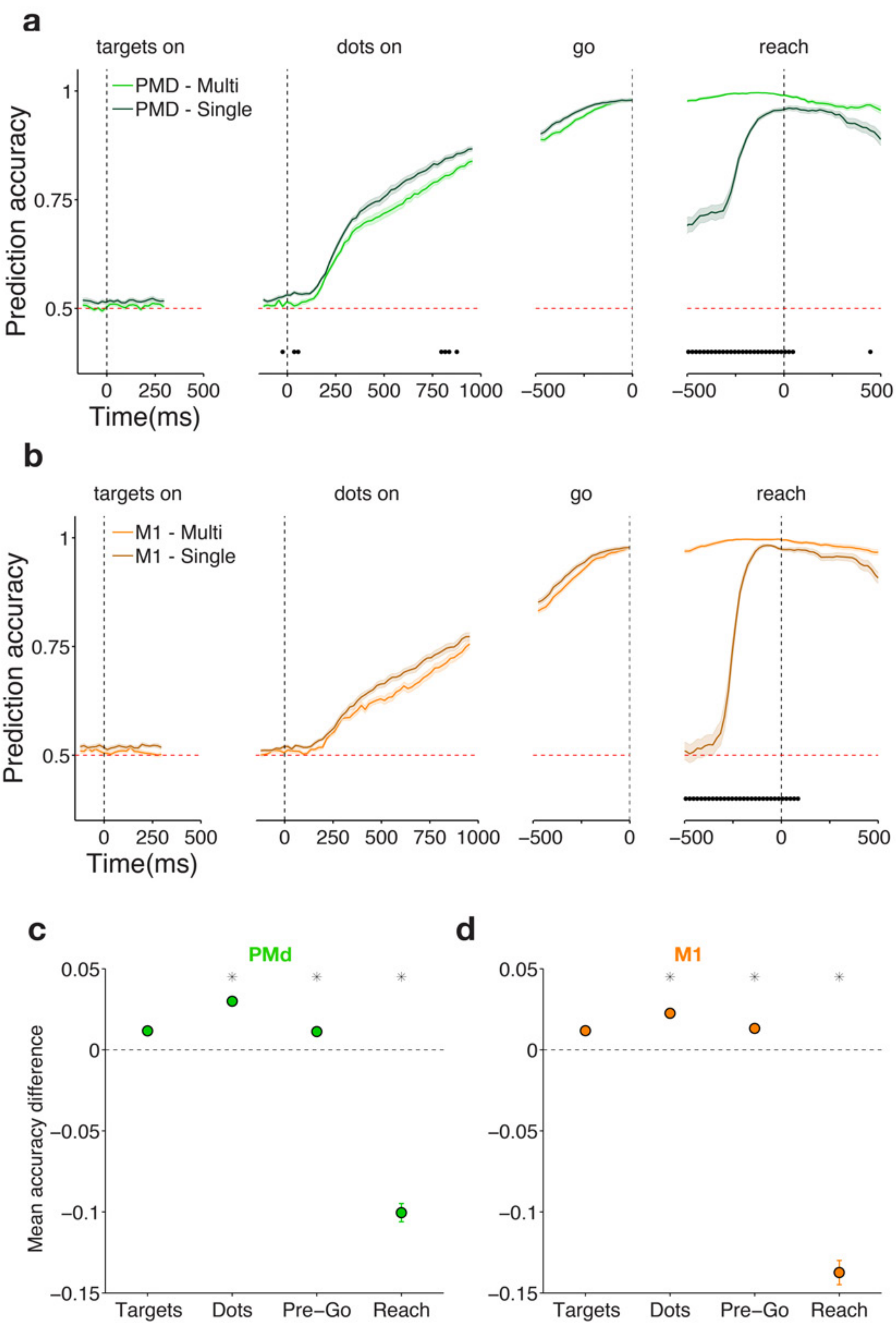


**Supp. Figure 18 - Neural population choice prediction accuracy on single trials in the fixed duration task: multiple vs single classifiers (pooled results across 2 monkeys).**

**a) Single and multiple classifiers result in similar performance for targets, dots and pre-go epochs but not for reach epoch for PMd.** Average prediction accuracy (see Methods) over time +- SEM for both monkeys PMd. Single (multiple) classifier results are plotted in dark (light) green. Data for multiple classifiers are re-plotted from Figure 2a. Black dots denote time bins for which the prediction accuracy was significantly different between the two areas (Wilcoxon signed-rank $p<0.05$ Holm-Bonferroni correction for multiple comparisons). Single Classifier does slightly better for targets, dots and pre-go periods and much worse than multiple classifiers for reach period.

**b) Single and multiple classifiers result in similar performance for targets, dots and pre-go epoch but not the reach epoch for M1.** Equivalent to **a)** but for M1. Same conventions apply.

**c) Summary of performance difference between single and multiple classifiers within each epoch.** Average performance difference between single and multiple classifiers (accuracy difference in percentage correct) for each of the epochs plotted in **a)**. Positive number numbers correspond to better single classifier performance and negative numbers to better multiple classifier performance. Black asterisks correspond to windows for which the coherence effects were significantly larger than zero (Wilcoxon signed-rank test, $P<0.001$).

**d**) Same as **c)** for M1. For both areas ( **c)** and **d**) ) the difference of choice prediction accuracies between the single and the multiple classifiers was small and positive for the target, dots and pre-go epochs, demonstrating substantial choice representation stability in these periods (between 1% ±0.15% and 3% ±0.26%). In contrast, for the peri-movement period, the difference in prediction accuracies was strongly negative and significantly different from the dots and delay epochs (Wilcoxon signed-rank test $p<10^{-3}$), confirming choice representation instability (-10% ±0.56% /-14% ±0.75% for PMD/M1, respectively).

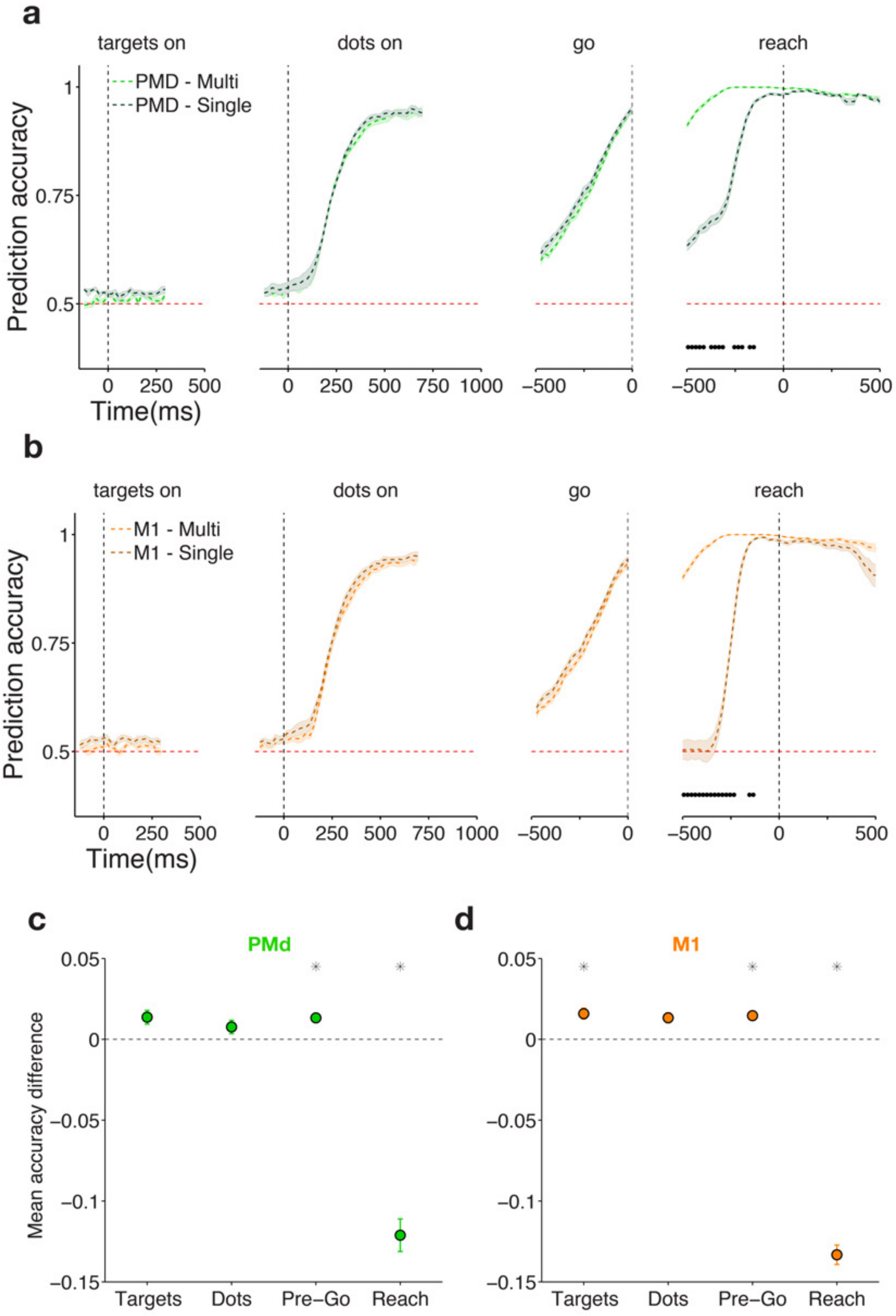


**Supp. Figure 19 - Neural population choice prediction accuracy on single trials in the variable duration task: multiple vs single classifiers (pooled results across 2 monkeys).**

**a)-d**) same as **Supp. Figure 18 a)-d)** but for the variable duration task**.**

For both areas ( **c)** and **d**) ) the difference of choice prediction accuracies between the single and the multiple classifiers was small and positive for the target, dots and pre-go epochs, demonstrating substantial choice representation stability in these periods (between 0.8% ±0.42% to 1.6% ±0.34%). In contrast, for the peri-movement period, the difference in prediction accuracies was strongly negative and significantly different from the dots and delay epochs (Wilcoxon signed-rank test $p<10^{-3}$), confirming choice representation instability (-12%±1.01% / -13%±0.46% for PMD/M1, respectively).

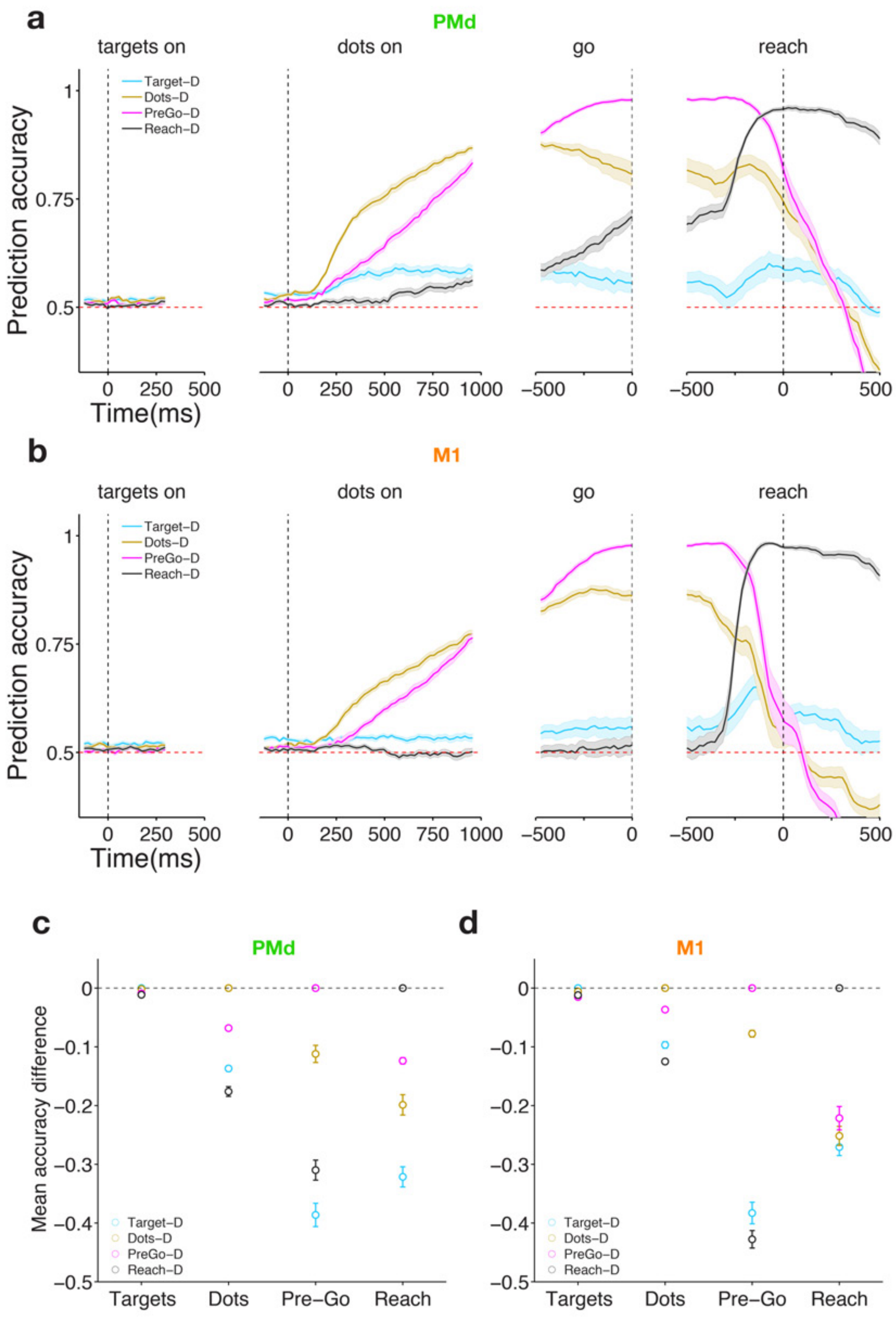


**Supp. Figure 20 - Neural population choice prediction accuracy on single trials in the fixed duration task when applying classifiers across epochs (pooled results across 2 monkeys).**

**a) Only dots and pre-go classifiers perform well across epochs in PMd.** Average prediction accuracy (see Methods) over time +- SEM for both monkeys for decoders trained in the targets (cyan), dots (dark yellow), pre-go (magenta) and reach (black) periods. If the choice subspaces for two independent epochs are similar, the decoder from one epoch ought to accurately predict choice in the other epoch. Dots decoder performs well during pre-go period and vice-versa. Targets and reach decoders perform poorly across other epochs.

**b) Only dots and pre-go classifiers perform well across epochs in M1.** Equivalent to **a)** but for M1. Same conventions apply.

**c) Summary of performance difference between single and multiple classifiers within each epoch.** Average performance difference between within-epoch classifier and across-epoch classifiers for each of the epochs plotted in **a)**. Error bars correspond to +- SEM across sessions. Zero difference corresponds to the performance of the classifier trained and tested within the same epoch.

**d)** Same as **c)** for M1. For both PMd **(c)** and M1 (**d**) in the dots and pre-go periods the loss in decoding accuracy across epochs was fairly small (pre-go decoder during dots: -6.8% ±0.34% /-3.7% ±0.25%, dots decoder during pre-go: -11% ±1.46% /-7.8% ±0.57%, for PMd/M1), but not for other pairs of epochs (e.g., reach decoder during pre-go: -31.0% ±1.71% /-42.8% ±1.5% for PMd/M1). The small negative values for dots and pre-go epochs suggest that while the subspaces were largely overlapping they were not perfectly identical.

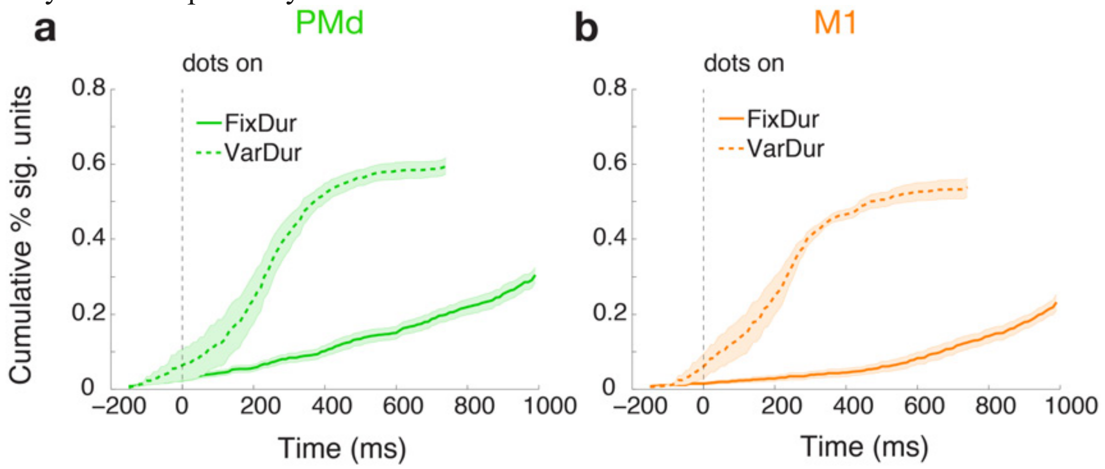


**Supp. Figure 21 – Recruitment of choice predictive cells is accelerated for both brain areas under uncertainty conditions. a-b)** Same as Figure 7 **a-b)** for monkey H.

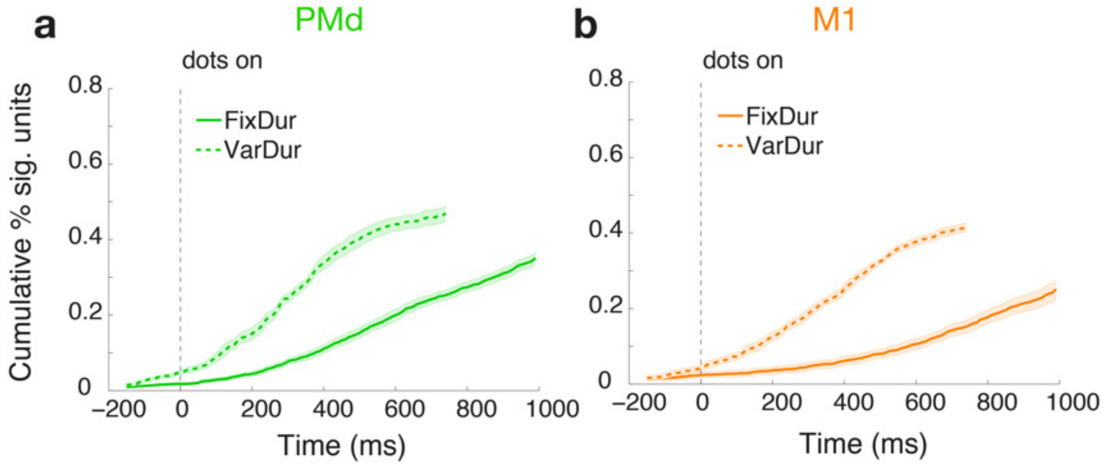


**Supp. Figure 22 – Recruitment of choice predictive cells is accelerated for both brain areas under uncertainty conditions. a-b)** Same as Figure 7 **a-b)** for monkey F.

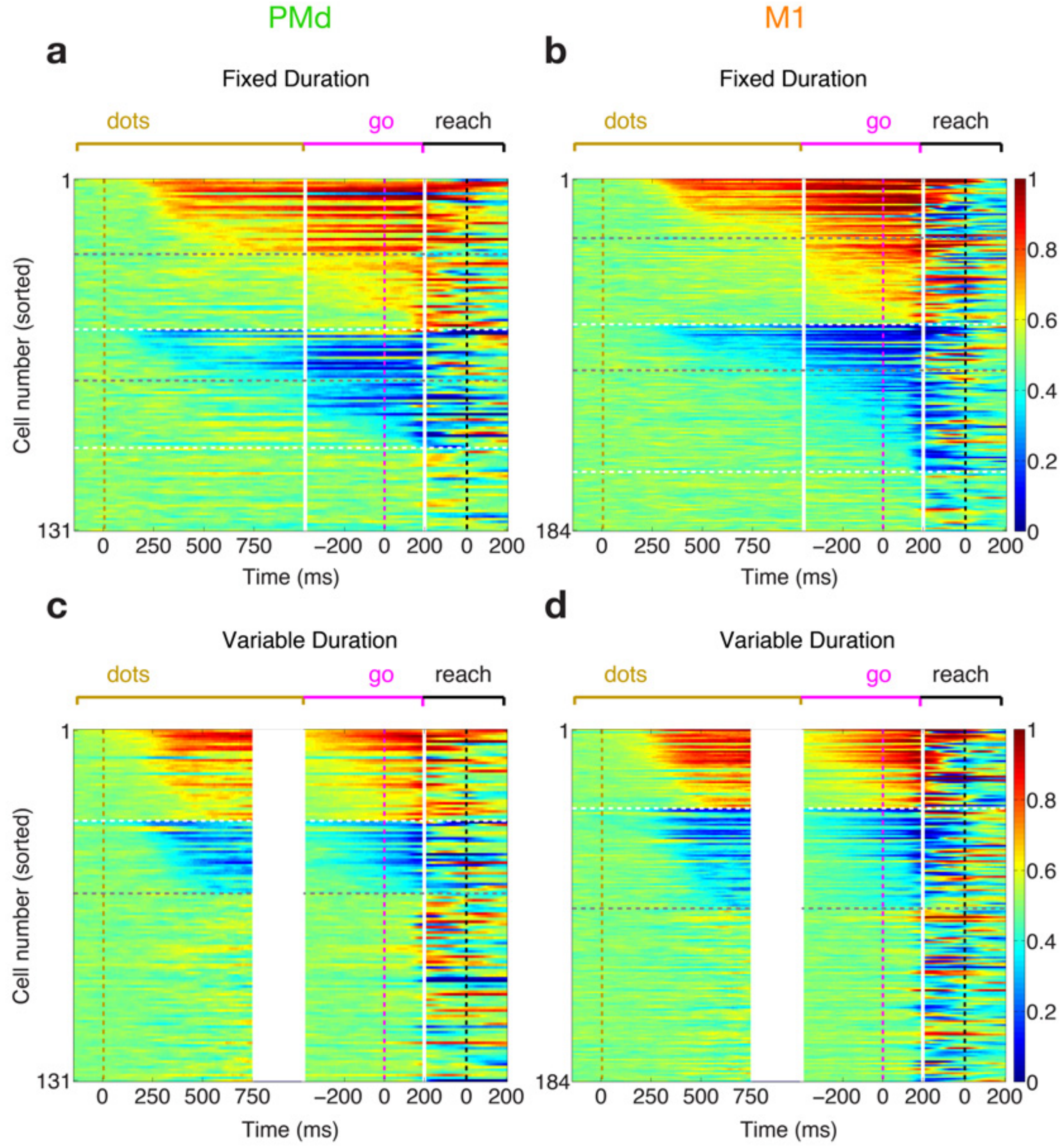


**Supp. Figure 23 – Side preference of choice predictive cells is largely maintained**

**during stimulus and pre-go periods for both brain areas and tasks. a) Individual unit choice predictive activity is stable during dots presentation and builds up slower in the fixed duration task.** Area under ROC traces for all units recorded in one session in PMd (same units as **Fig.7c**). Traces (one row for each unit) were sorted by onset of significant choice modulation during the dots presentation for right preferring units (red traces) and left preferring units (blue traces). White solid lines denote the separation of epochs (dots end, and go cue +200 ms); golden, magenta and black dashed lines mark the dots onset, go cue and reach initiation, respectively. Top horizontal dashed white line separates right from left preferring units and bottom horizontal dashed white line separates the latter from the remainder of the population (below). Horizontal dashed gray line separates cells with significant choice modulation during dots (above) from cells with significant choice modulation starting in the delay period. Data from Monkey F. **b)** Same as **a)** for M1 (same units as **Fig.7d**).

**c) Individual unit choice predictive activity is stable during dots presentation and builds up faster in the variable duration task.** Figure conventions as in **a)**. Data from Monkey F (same units as **Fig.7e**). **d)** Same as **c)** for M1 (same units as **Fig.7f**).

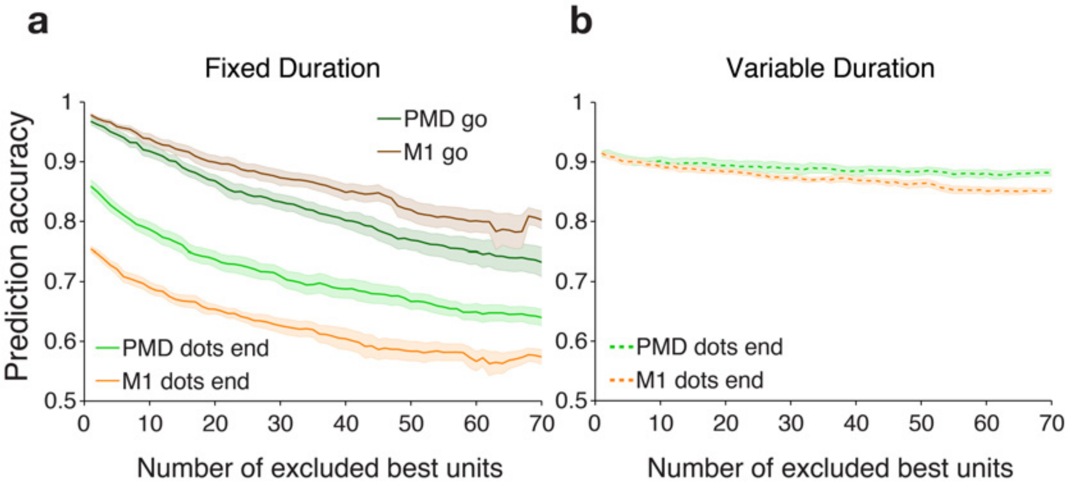


**Supp. Figure 24 - Choice signal is robust and distributed across the population of cells in both areas and both tasks. a-b)** Same as figure 8 **a-b)** for Monkey H.

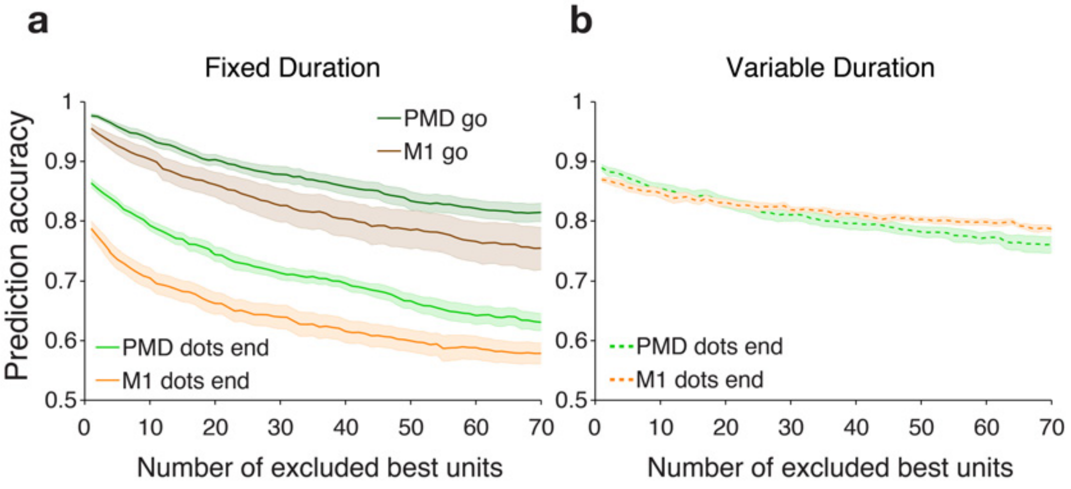

**Supp. Figure 25 - Choice signal is robust and distributed across the population of cells in both areas and both tasks. a-b)** Same as figure 8 **a-b)** for Monkey F.
